# Fast-conducting mechanonociceptors uniquely engage reflexive and affective pain circuitry to drive protective responses

**DOI:** 10.1101/2025.11.11.687663

**Authors:** Karina Lezgiyeva, Jingyi Liu, Karen Nguyen, Michelle M. DeLisle, Frank C. Ko, Spencer Fullam, Alia M. Obeidat, Josef Turecek, Ilayda Alkislar, Brendan P. Lehnert, Rosa I. Martinez-Garcia, Riya Sivakumar, Jinheon Choi, Ofer Mazor, Lilit Garibyan, Nikhil Sharma, Alan J. Emanuel, Anne-Marie Malfait, Rachel E. Miller, David D. Ginty

## Abstract

Nociceptors detect damaging stimuli and evoke pain in healthy animals. We conducted an optogenetic activation screen to identify genetically defined nociceptor populations that elicit place aversion and nocifensive behaviors in response to stimulation. *Smr2^Cre^-* and *Bmpr1b^Cre^-*labeled Aδ high-threshold mechanoreceptors (HTMRs) emerged as two of the few nociceptor populations, and we focused on investigating their physiological, morphological, functional, and synaptic properties. These neurons densely innervate skin and other organs, are activated only by intense, potentially damaging stimuli, and are necessary for protective responses to sharp mechanical stimuli. Centrally, Aδ-HTMRs projections span multiple spinal segments and terminate across spinal cord laminae, forming strong, monosynaptic connections onto anterolateral tract projection neurons, including antenna cells of the deep dorsal horn. Aδ-HTMRs also engage a local spinal reflex circuit enabling a remarkably rapid limb withdrawal. Thus, Aδ-HTMRs are myelinated nociceptors with unique properties that can be exploited for development of new analgesics.

## Introduction

Despite its evolutionary importance and clinical burden, the biological underpinnings of pain remain incompletely understood and mounting evidence suggests that different types of pain have different underlying mechanisms^1-8^. Nociceptive pain, such as that caused by acute physical trauma or exposure to extreme temperatures, alerts us to harmful or damaging environmental stimuli and promotes avoidance. The neural signals underlying nociceptive pain originate in primary somatosensory neurons housed in dorsal root ganglia (DRG) or trigeminal ganglia and called nociceptors. Nociceptors can be categorized based on soma size and conduction velocity as large, fast-conducting, myelinated A-fiber neurons and small, slow, unmyelinated C-fiber neurons^9,10^. Nociceptors have been physiologically identified in every mammalian species studied, including mouse and human^9,11-18^, and more recent RNA sequencing studies have confirmed the conservation of transcriptionally defined putative A- and C- fiber nociceptors across species^19-21^.

Since pain is challenging to assess in non-human animals, definitions of nociceptors in preclinical research can be contentious. While originally referring to neurons that respond exclusively to damaging stimuli, the definition is often expanded to include sensory neurons with lower activation thresholds and wide dynamic ranges^9^. Additionally, immunohistological and transcriptional features associated with nociceptors, including expression of TrkA, CGRP, Substance P, TrpV1, Na_v_1.8, and IB4 binding, among others, are often used in place of physiological characterization^21,22^. Finally, there are challenges in distinguishing the properties of sensory neurons in healthy, uninjured animals versus animals in an altered state of disease, such as chronic pain or inflammation^23-25,26^. To avoid ambiguity in describing sensory neuron functions, we propose a strict definition of nociceptors based on the original definitions of Charles Sherrington and Ed Perl^18,27^. Nociceptors are sensory neurons that respond to noxious or damaging stimuli and, when activated, evoke pain and nocifensive behaviors in healthy, naive animals.

There are well over a dozen distinct types of DRG sensory neurons, and mouse genetic tools now allow unprecedented access to most of these populations for observation and manipulation^23,28-43^. Over the preceding decade, most mouse genetics-enabled work has focused on small unmyelinated C fibers, and it is often suggested that most nociceptors are C-fiber neurons^10,31,36,37,44^. Comparatively few studies have reported gaining genetic access to A-fiber high-threshold mechanoreceptors (A-HTMRs)^23,45,46^. Thus, since their discovery over 50 years ago^18^, the morphologies, peripheral targets, functional properties, central projection patterns, and synaptic targets of A-fiber nociceptors remain incompletely understood. Here, we leveraged recently developed mouse genetic tools for a comparative behavioral analysis of most DRG sensory neuron types of the mouse to determine genetically defined A-fiber and C-fiber nociceptor populations. Two transcriptionally distinct populations stood out as myelinated nociceptors and we undertook a detailed characterization of these A-fiber nociceptors to better understand their contributions to nociceptive pain.

## Results

### Two Aδ-HTMR populations drive uniquely fast and robust nocifensive behaviors and aversion

We first sought to identify nociceptor populations innervating glabrous skin of mice. For purposes of this study, we define a nociceptor as a primary somatosensory neuron that fires action potentials in response to damaging or potentially damaging stimuli and, when activated, promotes both a nocifensive or protective behavioral response and an aversion to the activating stimulus.

To identify sensory neurons that meet our behavioral definition of a nociceptor, we used mouse genetic tools for select sensory neuron subtype access and performed an optogenetic activation behavioral survey across 10 distinct cutaneous A-fiber and C-fiber DRG neuron subtypes, as summarized in Table 1. Most LTMR subtypes were excluded from this analysis because they are maximally responsive to innocuous stimuli^23^ and, when activated, they do not evoke nocifensive behaviors and, in the case of C-LTMRs and Aδ-LTMRs, they do not innervate glabrous skin^47^.

To assess behavioral responses to minimal activation of the different DRG sensory neuron subtypes, we optically stimulated the hindpaws of mice expressing the light-gated ion channel ReaChR in different sensory neuron populations and recorded their behavioral responses using high-speed videography (Figure 1A). We used short 5-ms light pulses as they generate only a single action potential in *Smr2^Cre^*-labeled Aδ-HTMR neurons, as demonstrated using *in vivo* electrophysiological recordings (Figure 1A). Strikingly, when either *Smr2^Cre^*-labeled or *Bmpr1b^Cre^*-labeled Aδ-HTMR neurons were targeted, this stimulus reliably generated a rapid (∼20 ms latency) and robust response characterized by paw withdrawal, shaking, and jumping, along with occasional guarding (Figure 1B-C, Video S1). Activation of C-heat thermoreceptors also evoked a robust response, although with a much longer latency (200-400 ms), while activation of C-cold thermoreceptors typically evoked a only subtle limb withdrawal but no paw shaking, guarding, kicking, or jumping. The four C-fiber polymodal sensory neuron subtypes and Aβ RA-LTMRs did not evoke an observable behavioral response upon activation. This was not due to a failure of opsin activation since light-evoked action potentials could be detected in the spinal cord upon stimulation of all neuron types that did not evoke a behavioral response, albeit with variable reliability^48^ (Figure S1).

**Figure 1.**
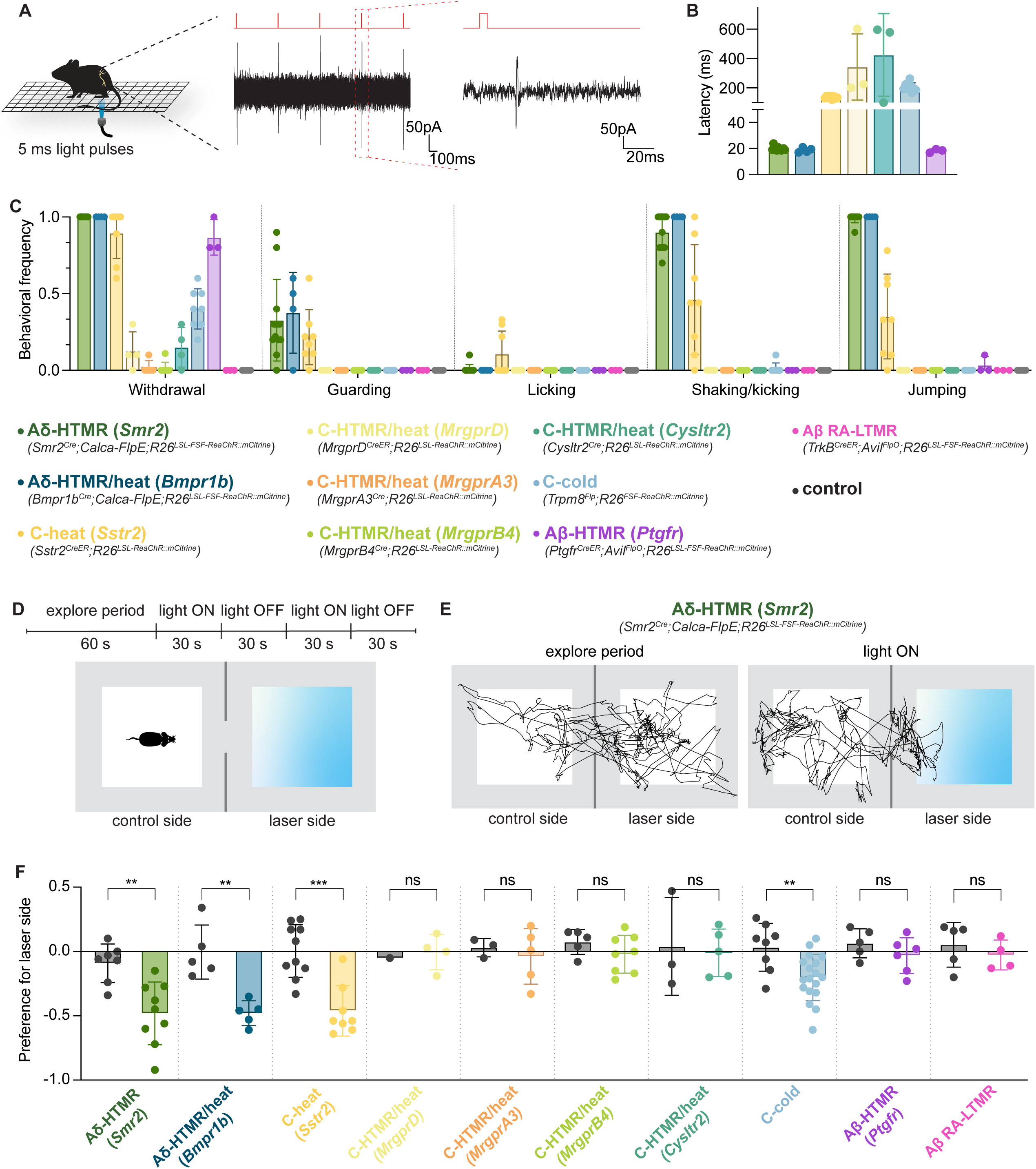
An optogenetic activation behavioral screen of DRG sensory neuron types. (A) Left: Optogenetic stimulation paradigm. Right: *In vivo* loose patch electrophysiology reveals that 5-ms pulses trigger single action potentials in *Smr2^Cre^*-labeled neurons. Right trace is a zoomed-in version of the light-evoked spike outlined in red on the left trace. (B) Average latency (mean ± SD) of behavioral response to 5-ms light stimulation of glabrous hindpaw in mice expressing the opsin ReaChR in different sensory neuron populations. Each dot represents one animal (n = 12, 4, 8, 3, 3, 8, 3 from left to right). (C) Average frequency (mean ± SD) of select behavioral responses to 5-ms light stimulation of different sensory neuron populations. See Video S1 for a representative example of Aδ-HTMR-driven behavior. Littermate controls were animals that lacked either one of the driver alleles or the ReaChR allele. Each dot represents one animal (n = 12, 4, 8, 4, 5, 8, 4, 8, 3, 3, 3 from left to right). (D) Optogenetic real-time place preference paradigm. (E) Representative locomotion traces from one animal expressing ReaChR in Aδ-HTMRs (*Smr2^Cre^; Calca-FlpE; R26^LSL-FSF-ReaChR::mCitrine^*) during the exploration (left) and the stimulation (right) periods. (F) Average relative preference (mean ± SD) for the laser side in mice expressing ReaChR in different sensory neuron populations and littermate controls. Negative values indicate avoidance of the laser side and positive values indicate preference for the laser side. Littermate controls were animals that lacked either one of the driver alleles or the ReaChR allele. For *MrgprB4^Cre^; R26^LSL-ReaChR::mCitrine^* animals, wild-type CD1 mice tested alongside the mutants were used as controls. Each dot represents one animal (n = 7, 9, 5, 5, 10, 8, 1, 4, 3, 5, 5, 8, 3, 5, 8, 18, 5, 6, 5, 4 from left to right, *\*\*p* ≤ 0.01, *\*\*\*p* ≤ 0.001, unpaired t test).

We also tested a mostly undescribed population of glabrous skin innervating neurons, which we refer to as Aβ-HTMRs. Aβ-HTMRs were first physiologically identified as moderate pressure receptors in 1968^11^ but have not been well characterized. This population of sensory neurons can be labeled with the *Ptgfr^CreER^* mouse line and resembles the Aβ Field-LTMRs previously described in hairy skin^41^. The *Ptgfr*, *Smr2*, and *Bmpr1b* transcripts are largely non-overlapping (Figure S1A), suggesting the existence of three unique A-fiber HTMR populations. The *Ptgfr^CreER^*-based labeling strategy is moderately selective and very efficient (Figure S2C-E), and the labeled neurons have large soma diameters and express NFH and *Ntrk3* but not CGRP (Figure S2A, B). Moreover, while most *Ptgfr^+^* neurons do not coexpress Na_v_1.8 (*Scn10a*) (Figure S2B), over a quarter of the labeled neurons do express this Na^+^ channel (Figure S2H), suggesting that a recently described population with fast HTMR properties^49^ may at least partially overlap with the genetically labeled Aβ-HTMRs described here. Physiological recordings revealed that these neurons conduct action potentials in the Aβ velocity range and have indentation thresholds higher than those of Aβ-LTMRs but lower than Aδ-HTMRs (Figure S2I-L). In addition, like other Aβ fiber DRG neurons, Aβ-HTMRs have one central branch projecting directly to the dorsal column nuclei (DCN) and other branches that terminate densely in the LTMR recipient zone of the deep dorsal horn (Figure S2M, N). However, unlike other Aβ neuron subtypes, the peripheral axons of these large diameter CGRP-negative Aβ-HTMRs exhibit expansive arbors in the skin that penetrate the epidermis and form free nerve endings (Figure S2O, P). Interestingly, human glabrous skin also contains CGRP-negative epidermal endings that are NFH-positive in the dermis, and these human neurons may correspond to the genetically labeled mouse Aβ-HTMRs described here (Figure S2R). When Aβ-HTMRs were optogenetically activated, animals exhibited only a mild hindlimb withdrawal response, with no paw shaking, guarding, kicking, or jumping (Figure 1C).

To measure behavioral avoidance, we devised a real-time place preference paradigm. Animals expressing ReaChR in one of the ten sensory neuron subtypes were placed into a two-sided chamber with transparent floors, where one of the sides was illuminated from below with laser light (Figure 1D). We found that mice with ReaChR expressed in either of the two Aδ-HTMR subtypes or C-heat thermoreceptors, but not littermate controls, robustly and consistently avoided the laser side during illumination periods (Figure 1E, F). The mice also exhibited an aversion to the stimulation of C-cold thermoreceptors, consistent with prior reports of mice avoiding cooler temperatures in a Trpm8-dependent manner^50^. In contrast, stimulating any of the other six sensory neuron subtypes did not evoke an aversion to the stimulation side of the chamber (Figure 1F). This included Aβ-HTMRs and the four polymodal C-fiber types, often referred to as nonpeptidergic nociceptors.

Thus, of the 10 cutaneous sensory neuron types tested, only three populations – two Aδ-HTMRs and C-heat thermoreceptors – meet our behavioral criteria of a nociceptor, whereas none of the C-HTMRs examined here met the criteria. Two additional C-fiber populations (*Oprk1^+^* and *Calca^-^/TrpV1^high^* DRG neurons) were not tested here because genetic labeling tools for these neurons are not available, and these populations exhibit robust *TrpV1* expression but low or undetectable *Piezo2* expression^28^, suggesting thermal responsiveness. Together, our observations indicate that acute mechanonociception is mediated primarily by fast conducting mechanosensory neurons. Thus, we focused our physiological, anatomical, and circuit-level analyses on these two Aδ-HTMR populations to gain insight into the basis of mechanical nociceptive pain.

### Neurons labeled by *Smr2^Cr^*^e^ and *Bmpr1b^Cre^* mouse lines are Aδ-HTMRs in mouse glabrous skin

We used *in vivo* calcium imaging to observe and compare responses of *Smr2^Cre^*- and *Bmpr1b^Cre^*-labeled neurons to mechanical and thermal stimuli applied to glabrous skin (Figure 2A). As previously observed in hairy skin^23,46^, both populations exhibited high mechanical force thresholds, showing no responses to brush and vigorous responses to higher forces, including skin indentation with stiff von Frey filaments, poking with closed forceps, and pinching (Figure 2B, C, Figure S3A). *Bmpr1b^Cre^*-labeled neurons have somewhat lower mechanical thresholds compared to *Smr2^Cre^*-labeled neurons (Figure 2C, Figure S3A). The difference was more pronounced when using von Frey filaments for stimulation compared to the 200-μm indenter tip (Figure S3A), suggesting that sharpness, in addition to force, may be a distinguishing feature for activating these two genetically defined Aδ-HTMR populations. We also observed that neurons of both Aδ-HTMR populations have overlapping physiological receptive fields, as multiple neurons simultaneously responded to indentations delivered to a single spot with the 200-µm probe tip (Figure 2D).

**Figure 2.**
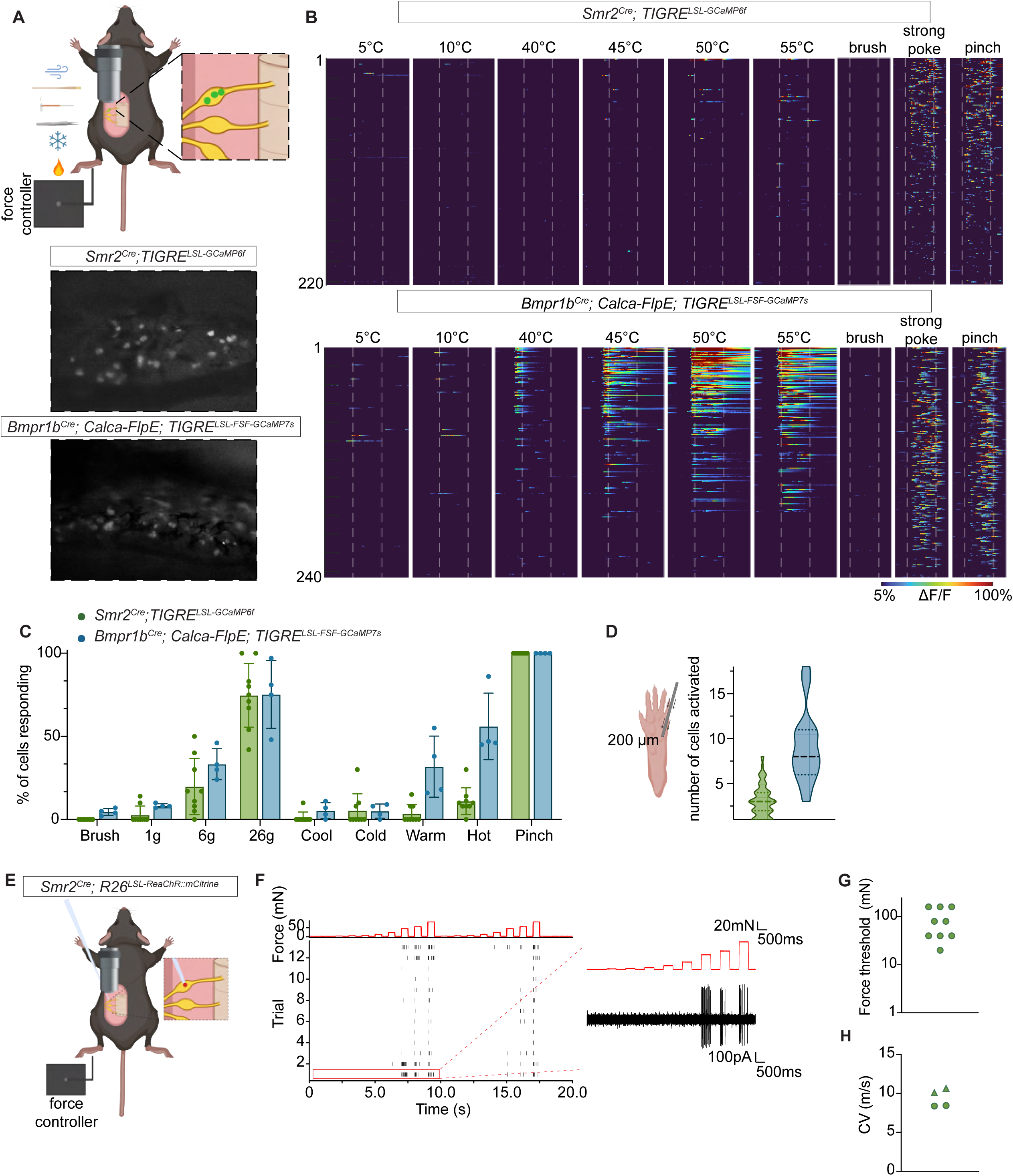
Physiological characterization of glabrous skin innervating neurons labeled by the *Smr2^Cre^* and *Bmpr1b^Cre^* mouse lines. (A) *In vivo* epifluorescent calcium imaging. Stimuli were presented to the glabrous hindpaw, and calcium responses of genetically labeled sensory neurons were recorded in the cell bodies of lumbar level 4 DRGs. Representative fields of view are shown. (B) Calcium indicator responses of *Smr2^Cre^*- and *Bmpr1b^Cre^*-labeled neurons to select stimuli (n = 223 neurons across 9 animals for *Smr2^Cre^*-labeled neurons, 246 neurons across 3 animals for *Bmpr1b^Cre^*-labeled neurons). (C) Average percentage (mean ± SD) of neurons in a field of view responding to select stimuli. The number of neurons responding to pinch is used as a denominator. 1 g, 6 g, and 26 g are the weights of von Frey filaments. Cool is defined as 20 and 25°C, cold as 5, 10, and 15°C, warm as 35 and 40°C, and hot as 45, 50, and 55°C. Each dot represents an animal and an independent recording session (n = 9 for *Smr2^Cre^*-labeled neurons and 4 for *Bmpr1b^Cre^*-labeled neurons). (D) Number of cells in a field of view simultaneously responding to an indentation with a 200-µm probe tip across a range of forces (10 – 400 mN) (n = 84 indentation spots across 16 animals for *Smr2^Cre^*-labeled neurons, and 22 indentation spots across 4 animals for *Bmpr1b^Cre^*-labeled neurons). (E) *In vivo* targeted loose patch electrophysiological recordings. Stimuli were presented to the glabrous hindpaw, and somatic extracellular currents of genetically labeled sensory neurons were recorded. (F) An example raster plot of indentation responses of an *Smr2^Cre^*-labeled neuron. A current trace of the responses outlined with a red line is shown on the right. (G) Force thresholds of *Smr2^Cre^*-labeled neurons. See Supplemental Figure 1 for a comparison between more broadly labeled sensory neuron populations. Each dot represents a unit recorded from a single animal (n = 9 units from 9 animals). (H) Conduction velocities of *Smr2^Cre^*-labeled neurons. Each dot and triangle represents a unit (n = 4 units from 3 animals, dots represent data acquired with loose patch configuration, triangles represent data acquired with sharp glass electrodes).

Consistent with the very high mechanical thresholds of *Smr2^Cre^*-labeled neurons, we found that the mechanotransduction channel Piezo2 is dispensable for their responses (Figure S3B-D). This is in line with findings from both transcriptional and *in situ* hybridization analyses indicating that *Piezo2* expression is low in the *Smr2^+^* population^28^ (Figure S3E) compared to other mechanosensory neuron types. Similarly, our previous work revealed that Piezo2 contributes to, but is not required for, mechanically evoked responses of *Bmpr1b^Cre^*-labeled neurons that innervate the colon^51^.

When applying thermal stimuli, we found that in hairy skin, a minority of the *Bmpr1b^Cre^*-labeled neurons respond to noxious cold, and about half of the *Smr2^Cre^*-labeled neurons respond to noxious heat (Figure S3F-G, see also^23^). In contrast, in glabrous skin, few neurons of both populations exhibit appreciable cold responses, about half of the *Bmpr1b^Cre^*-labeled neurons respond to noxious heat, and *Smr2^Cre^*-labeled neurons show minimal noxious heat responses (Figure 2B, C). Given the robust responses to heat in some *Bmpr1b^Cre^*-labeled neurons, the term Aδ-HTMR/heat is more appropriate for these neurons. The heterogeneous heat responses of glabrous skin-innervating *Bmpr1b^Cre^*-labeled neurons suggest the presence of subtypes within this population.

We also observed that while some glabrous-innervating *Sstr2^CreER^*-labeled neurons responded to pinch, most were tuned to noxious heat, vigorously responding to 50-55°C (Figure S3H). Thus, both Aδ-HTMR subtypes and *Sstr2^CreER^*-labeled C-heat thermoreceptors respond almost exclusively to noxious, potentially damaging stimuli, in alignement with the original nociceptor definition^18,27^.

We next used *in vivo* loose-patch electrophysiology to evaluate Aδ-HTMR action potential firing patterns, conduction velocities, and waveforms (Figure 2E-H. Figure S3I), focusing on the more homogeneous *Smr2^Cre^*-labeled population. *Smr2^Cre^*-labeled neurons produced action potentials with wide waveforms in response to high forces applied to the skin (Figure S3I). In addition, they fired repetitively and adapted slowly in response to ramp and hold mechanical stimuli (Figure 2F), although their firing patterns were much less stereotyped or reliable compared to those previously observed for Aβ-LTMRs^52^. Conduction velocity measurements of *Smr2^Cre^-*labeled neurons (Figure 2H) confirmed that they fall within the Aδ range, similar to a previous report of hairy skin-innervating *Bmpr1b^+^* neuron conduction velocity (Figure 2H; see^46^, neurons recorded there are likely *Bmpr1b^+^*; see also comparative conduction velocity measurements in Figure S2E – CGRP^+^ neurons recorded here are likely *Smr2^+^* or *Bmpr1b^+^* Aδ-HTMRs).

### Aδ-HTMRs correspond to transcriptionally defined putative nociceptor populations

The homogeneity of physiological response properties of genetically labeled Aδ-HTMRs was reflected in their molecular and soma size homogeneity (Figure S4). Both *Bmpr1b^Cre^*- and *Smr2^Cre^*-labeled populations exhibited medium/large-diameter cell bodies, falling on the larger end of the CGRP^+^ cell body diameter spectrum (Figure S4C). Each population comprises ∼45% of the medium/large diameter CGRP^+^ cells in lumbar DRGs (Figure S4D), suggesting that these two populations account for most if not all medium/large diameter CGRP^+^ neurons in lumbar DRGs. Virtually all Aδ-HTMRs of both types expressed CGRP and TrkA but did not bind IB4, nor did they express Calb, a marker of A-fiber LTMRs (Figure S4E). The majority of genetically labeled Aδ-HTMRs also expressed NFH, consistent with their medium/large soma size (Figure S4). Most Aδ-HTMRs did not express detectable levels of the heat-sensitive ion channel TrpV1 (Figure S4), and the few neurons that did express TrpV1 may account for the hairy skin-innervating *Smr2^Cre^*-labeled neurons and the glabrous skin-innervating *Bmpr1b^Cre^*-labeled neurons that exhibited pronounced responses to heat (Figure 2B, C, Figure S3F, G). Neither Aδ-HTMR population expressed the ATP receptor P2XR3, suggesting that P2XR3-dependent purinergic signaling does not underlie sensory transduction in these neurons (Figure S4E).

We also explored the transcriptional profile of genetically labeled Aδ-HTMRs. We first confirmed that the *Cre*-based genetic lineage labeling strategies employed here largely recapitulate *Smr2* and *Bmpr1b* marker gene expression of adult neurons (Figure S5A-B). We then considered whether transcripts commonly associated with nociceptor identity^19,21,53^ are expressed in these two genetically labeled populations. Our findings generally agree with predictions made based on transcriptomic data analyses: both populations coexpressed broad markers *Scn10a* (Na_v_1.8) and *Prdm12*, while *Kit* expression was biased towards the *Bmpr1b^Cre^*-labeled cells and *Chrna7* expression was biased towards the *Smr2^Cre^*-labeled cells (Figure S5C-D).

Thus, *Smr2^Cre^*- and *Bmpr1b^Cre^*-labeled DRG neurons are transcriptionally and physiologically distinct populations of Aδ-HTMRs, and together these two Aδ-HTMR populations account for most, if not all, medium/large diameter CGRP^+^, TrkA^+^ DRG neurons in the mouse.

### Aδ-HTMRs are required for normal responses to damaging mechanical stimuli

Since optical activation of either Aδ-HTMR population produced intense nocifensive responses and place aversion, we next asked whether one or both Aδ-HTMR subtypes are necessary for behavioral responses to noxious mechanical or thermal stimuli. To achieve Aδ-HTMR ablation, we generated mice expressing the human diphtheria toxin receptor in one or both populations and then delivered diphtheria toxin via intraperitoneal injections (Figure 3A). Strikingly, before any behavioral testing was performed, we observed that animals that had one or both Aδ-HTMR populations ablated developed facial and bodily wounds, while their co-housed littermate controls were unaffected (60% of Aδ-HTMR-ablated animals vs 0% of controls; Figure 3B-C, Figure S5E). This finding is reminiscent of tissue damage observed in human patients with loss-of-function pain disorders^54-57^ and suggests that a lack of fast mechanical pain signaling may lead to aberrant behaviors resulting in tissue damage.

**Figure 3.**
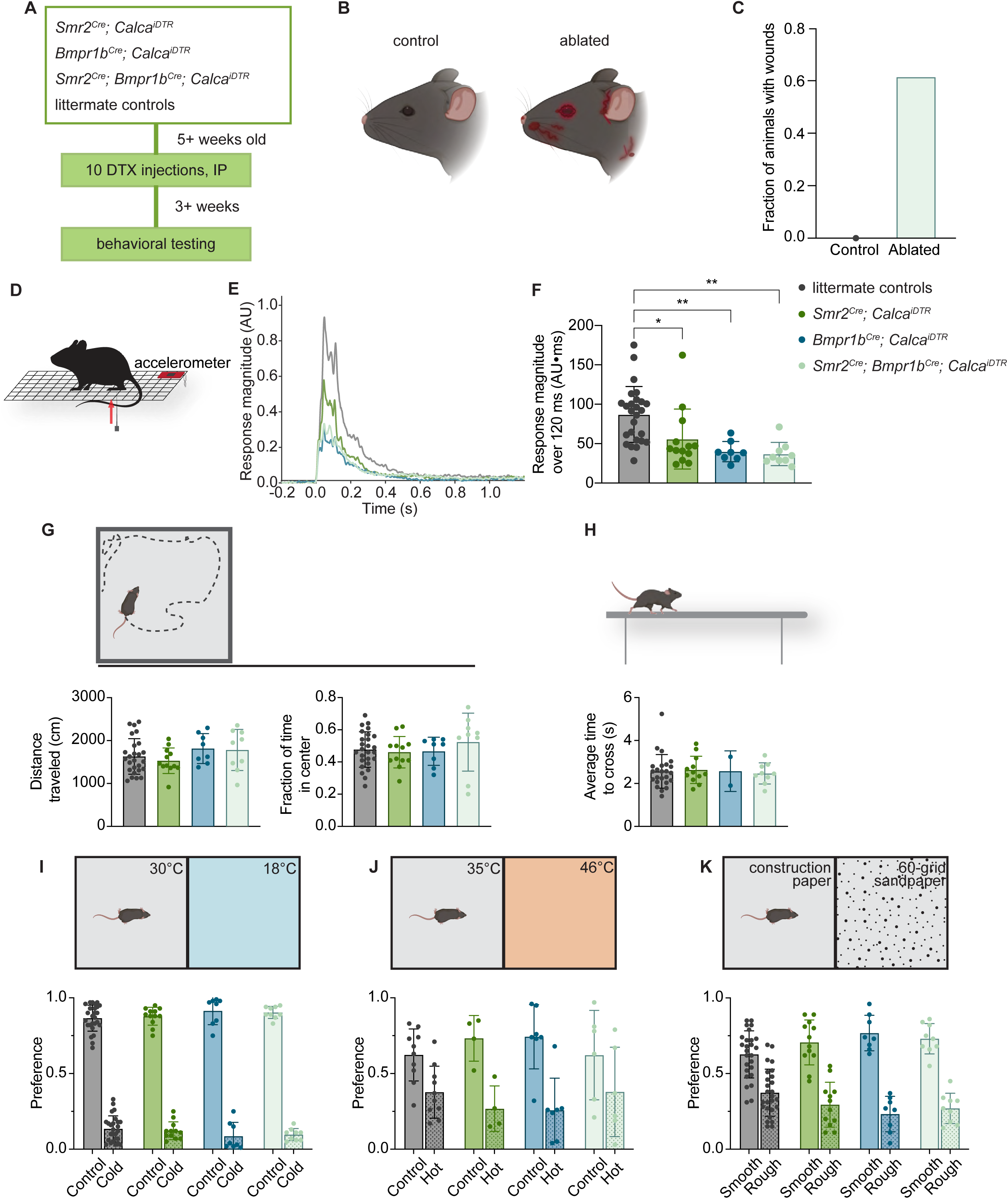
Aδ-HTMRs mediate behavioral responses to damaging mechanical stimuli. (A) Schematic of the chemogenetic ablation paradigm. Adult mice expressing human diphtheria toxin receptor in one (*Smr2^Cre^; Calca^iDTR^* or *Bmpr1b^Cre^; Calca^iDTR^*) or both (*Smr2^Cre^; Bmpr1b^Cre^; Calca^iDTR^*) Aδ-HTMR populations and littermate controls were treated with diphtheria toxin for 10 days. Behavioral testing was done three weeks after the last injection, or later. (B-C) Animals that had their Aδ-HTMRs ablated, but not littermate controls, developed facial and bodily wounds (0/12 controls and 16/26 ablated). (D) Pinprick behavior paradigm. Animals were placed on a wire rack, and a sharp pin was pushed into their hindpaws from below. Their movements were recorded using an accelerometer coupled to the wire rack. See Video S2 for an example of the behavior. (E-F) Mice with Aδ-HTMR ablation exhibited diminished responses to pinprick stimulation. Animals that lacked either the *Cre* alleles or the *Calca^iDTR^* allele and underwent toxin treatment were used as littermate controls. (E) shows average accelerometer responses from the different groups of mice. Each trace represents an average of multiple trials from several mice (n = 27 animals for controls, 14 for *Smr2^Cre^; Calca^iDTR^*, 8 for *Bmpr1b^Cre^; Calca^iDTR^*, 10 for *Smr2^Cre^; Bmpr1b^Cre^; Calca^iDTR^*). (F) shows response magnitude (mean ± SD) during the 120-ms period after stimulus onset. Each dot represents an animal (as in (E), n = 27, 14, 8, 10 from left to right; *\*p* ≤ 0.05, *\*\*p* ≤ 0.01, one-way ANOVA with Tukey’s multiple comparisons post-hoc test). (G) Chemogenetic ablation of Aδ-HTMRs does not affect animal locomotion, as evidenced by normal performance in the open field assay. The distance traveled (mean ± SD) and fraction of time spent in the center of the field (mean ± SD) are plotted. Each dot represents an animal (n = 26, 12, 8, 9 from left to right; no statistical difference between the groups, one-way ANOVA). (H) Chemogenetic ablation of Aδ-HTMRs does not affect animal locomotion and sensory-motor coordination, as evidenced by normal performance in the balance beam assay. Average times to cross the beam (mean ± SD) are plotted. Each dot represents an animal (n = 26, 12, 2, 6 from left to right; no statistical difference between the groups, one-way ANOVA). (I-K) Chemogenetic ablation of Aδ-HTMRs does not affect animals’ aversion to moderately cold (18°C) and hot (46°C) temperatures and rough texture. Preference is defined as the fraction of testing time spent on a given flooring, i.e., interacting with a given stimulus. Each dot represents an animal (for (I) and (K), n = 26, 12, 8, 9 from left to right; for (J), n = 10, 4, 7, 6 from left to right; no statistical difference between the groups, one-way ANOVA).

We next asked whether Aδ-HTMRs are required for behavioral responses to noxious mechanical stimuli, as suggested by earlier work^45,58^. To do this, sharp pinpricks were delivered to animals’ hindpaws, and their movements were recorded using an accelerometer attached to the wire rack that they stood on (Figure 3D, E, Video S2). We found that ablating either or both Aδ-HTMR populations was sufficient to significantly reduce the animals’ responses to pinprick, despite incomplete ablation (Figure 3E, F, Figure S5F). Ablation of both populations led to a ∼60% decrease in response magnitude. The diminished response to pinprick was not due to a general motor or sensory deficit, as mutant and control mice performed similarly on both open-field and balance beam tests (Figure 3G, H). Animals’ aversion to moderate cold (18°C) and hot (46°C) temperatures was also unaffected by Aδ-HTMR ablation, consistent with our findings of a lack of physiological responses to cold and limited responses to heat (<45°C) in these populations (Figure 3I, J). All ablated groups also exhibited normal aversion to sandpaper flooring, suggesting that other DRG mechanosensory neuron types mediate rough texture aversion (Figure 3K). Together, these findings indicate that Aδ-HTMRs mediate protective responses to sharp and acute mechanical stimuli and are not necessary for responses to at least some other aversive stimuli.

### Two Aδ-HTMR populations densely innervate glabrous skin, forming a large overlapping meshwork of free nerve endings that penetrate the epidermis

To better understand how Aδ-HTMRs respond to noxious mechanical stimuli applied to glabrous skin, we next sought to describe their peripheral arborization and termination patterns. We observed that both Aδ-HTMR populations innervate glabrous skin of the paws densely and uniformly (Figure S6A).

To visualize individual neurons of both Aδ-HTMR populations, sparse labeling experiments were done by injecting a low titer of CAG-CreON-FlpON-SEAP AAV9 virus into the glabrous hindpaw skin of *Smr2^Cre^; Calca-FlpE* and *Bmpr1b^Cre^; Calca-FlpE* animals (Figure 4A). This virus expresses alkaline phosphatase in a Cre and Flp recombinase-dependent manner. Consistent with our previous findings in hairy skin^23^, individual *Smr2^Cre^*- and *Bmpr1b^Cre^*-labeled neurons form large, expansive arbors in glabrous skin, with the *Bmpr1b^Cre^*-labeled neurons branching more elaborately than *Smr2^Cre^*-labeled neurons but less so, by comparison, than *Ptgfr^CreER^*-labeled Aβ-HTMRs (Figure 4B). Aδ-HTMR arbors appear larger than those of Aβ RA-LTMRs and comparable in size to Aβ-HTMR arbors. Inspired by the observation that 200 µm indentations activate multiple Aδ-HTMRs (Figure 2D), we also compared individual and population terminal densities to estimate the extent of homotypic overlap and found that 3-10 Aδ-HTMRs overlap within any given area of glabrous skin (Figure 4C, S6A). This overlapping innervation pattern contrasts with that of Aβ-LTMRs involved in fine discriminative touch^52,59^ and may provide better signal reliability at the expense of spatial acuity. Finally, both Aδ-HTMR populations were observed to penetrate the epidermis, where they lose NFH-immunoreactivity and form free nerve endings devoid of S100^+^ surrounding lamellar cells (Figure S6B). This is distinct from hairy skin-innervating *Bmpr1b^Cre^*-labeled neurons, which form circumferential endings around hair follicles in the dermis^23,46^. Thus, the two Aδ-HTMR nociceptor subtypes densely innervate glabrous skin where they terminate within the epidermis in a highly intermingled manner.

**Figure 4.**
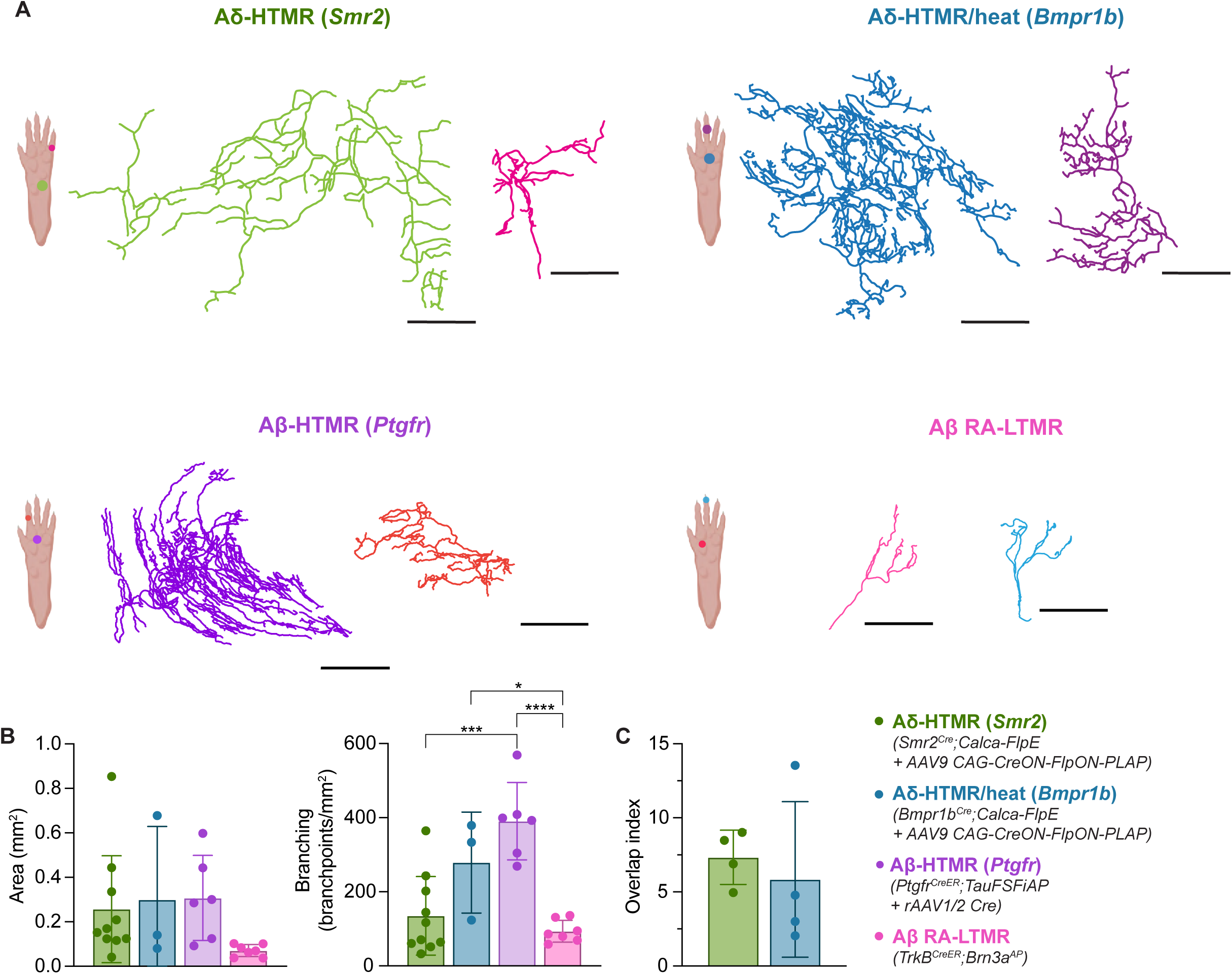
Aδ-HTMRs form expansive, highly overlapping terminal arbors in glabrous skin. (A) Representative reconstructions of glabrous skin whole-mount AP staining of single Aδ-HTMRs (labeled with *Smr2^Cre^* and *Bmpr1b^Cre^*), Aβ-HTMRs (labeled with *Ptgfr^CreER^*), and, for comparison, one Aβ-LTMR subtype (Aβ RA-LTMRs that form Meissner corpuscles, labeled with *TrkB^CreER^*). For Aβ-LTMRs, the data shown were acquired from previous studies^23,84^ and re-analyzed and plotted. Reconstructions are color-coded based on the arbor location. Scale bars are 300 μm. (B) Area (mean ± SD) and branching (mean ± SD) of glabrous skin arbors of the four sensory neuron populations tested. Each dot represents a reconstructed arbor (n = 10, 3, 6, 7 from left to right; no statistical difference in area measurements between the groups, *\*p* ≤ 0.05, *\*\*\*p* ≤ 0.001, *\*\*\*\*p* ≤ 0.0001, one-way ANOVA with Tukey’s multiple comparisons post-hoc test). (C) Aδ-HTMRs have overlapping anatomical receptive fields. The average overlap index (mean ± SD) of the two populations is plotted. Each dot represents a single density comparison (see Methods for details; n = 4 pairs for both populations).

### Aδ-HTMRs innervate many tissues and organ systems of the body

Inspired by observations of dramatic protective behaviors driven by skin-innervating Aδ-HTMRs, we asked whether *Smr2^Cre^*- and *Bmpr1b^Cre^*-labeled neurons innervate other, non-cutaneous organs and tissues, especially those that may be associated with mechanically evoked pain. We first examined bones and joints and found dense innervation by both Aδ-HTMR populations (Figure 5A-F), which is consistent with prior reports of CGRP^+^ Aδ-HTMRs in the bone^60,61^. In the knee joint, Aδ-HTMRs innervate the lateral synovium, the insertion of cruciate ligaments, infrapatellar fat pad (Hoffa’s fat pad), and the anterior horn of the meniscus (Figure 5A-D). Direct mechanical stimulation of these parts of the knee is associated with moderate to severe pain^62^, which may be mediated by Aδ-HTMRs.

**Figure 5.**
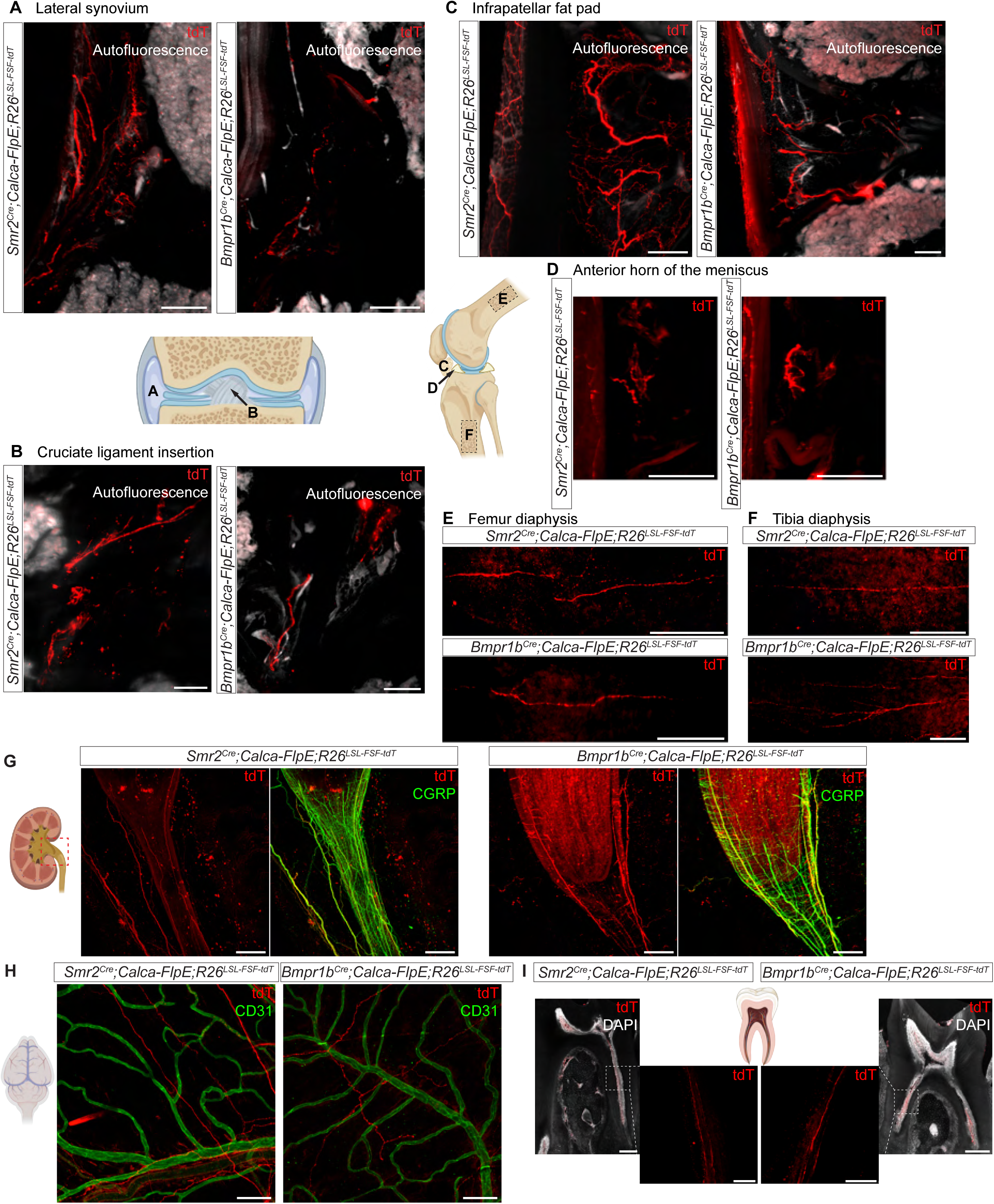
Aδ-HTMRs innervate many tissues and organ systems of the body. (A-F) Representative light-sheet microscopy images from *Smr2^Cre^; Calca-FlpE; R26^LSL-FSF-tdT^* and *Bmpr1b^Cre^; Calca-FlpE; R26^LSL-FSF-tdT^* intact whole-mount-stained knee joints (A-D, 200-µm maximum intensity projections), femurs, and tibias (E-F, 100-µm maximum intensity projections) collected from 10-week-old mice (n = 2 animals/reporter line). Scale bars are 200 µm. (G) Representative confocal images from *Smr2^Cre^; Calca-FlpE; R26^LSL-FSF-tdT^* and *Bmpr1b^Cre^; Calca-FlpE; R26^LSL-FSF-tdT^* whole-mount-stained kidneys. tdTomato-labeled axons are prominent in the renal pelvis and ureter regions. Scale bars are 200 µm. (H) Representative confocal images from *Smr2^Cre^; Calca-FlpE; R26^LSL-FSF-tdT^* and *Bmpr1b^Cre^; Calca-FlpE; R26^LSL-FSF-tdT^* whole-mount-stained cranial meninges. Blood vessels are labeled with CD31. Scale bars are 100 µm. (I) Representative confocal images from *Smr2^Cre^; Calca-FlpE; R26^LSL-FSF-tdT^* and *Bmpr1b^Cre^; Calca-FlpE; R26^LSL-FSF-tdT^* molar sections. Scale bars are 200 µm for whole-tooth images and 50 µm for insets.

Aδ-HTMR terminals of both types were also observed in the kidney and the walls of the urinary bladder (Figure 5G, Figure S6C), and previous work revealed innervation of the distal colon by *Bmpr1b^Cre^*-labeled but not *Smr2^Cre^*-labeled neurons^51^. In the kidney, most Aδ-HTMR innervation was concentrated in the calyces, the renal pelvis, and the ureters. These kidney structures can experience increased mechanical pressure during inflammation or obstruction, for example, by kidney stones, and this increased pressure can cause severe pain.

While our analyses focused on DRG Aδ-HTMRs innervating the body, the trigeminal ganglia (TG) house mostly the same transcriptionally defined populations of somatosensory neurons that supply innervation of the head^28,30^. We confirmed that neurons labeled by the *Smr2^Cre^* and *Bmpr1b^Cre^* lines in the TG are immunohistologically similar to those labeled in the DRG (medium/large diameter, CGRP^+^, IB^-^) (Figure S6D). In the head, trigeminal Aδ-HTMRs innervate both the meninges and tooth pulp (Figure 5H, I). These findings suggest that trigeminal Aδ-HTMRs may contribute to tooth pain and headache pathogenesis, including migraine.

Thus, Aδ-HTMRs innervate a range of non-cutaneous tissues across the body, where they likely function as fast mechanonociceptors underlying nociceptive pain.

### Aδ-HTMRs terminate across multiple dorsal horn laminae and form nest-like structures in laminae III-IV

The unique physiological, morphological, and functional properties of the two Aδ-HTMR populations prompted a detailed characterization of their central connectivity patterns with the goal of defining central circuits that these nociceptors engage to mediate rapid nocifensive responses and pain perception.

The spinal cord termination patterns of Aδ-HTMRs were found to be highly distinct from other sensory neuron types. C-HTMRs, C-cold and C-Heat neurons^23^ terminate mainly in the superficial spinal cord dorsal horn, whereas Aβ-LTMRs^43,47,63^ and Aβ-HTMRs (Figure S2N) terminate in the deep dorsal horn. Remarkably, Aδ-HTMRs terminate in both the superficial and deep laminae of the dorsal horn, forming clusters of axons in the latter (Figure 6A). Moreover, two observations support the idea that the two Aδ-HTMR subtypes provide most, if not all, CGRP^+^ sensory endings in the deep dorsal horn. First, when the two Aδ-HTMR populations were simultaneously labeled with Cre lines that are 73-95% efficient^23^ (Figure S5B), most CGRP immunoreactivity in laminae III-V was observed to overlap with the Aδ-HTMRs (73%) (Figure S7A, C). Second, when both Aδ-HTMRs populations were ablated, albeit incompletely (Figure S5B), much of deep CGRP immunoreactivity was lost (Figure S7B, D). When viewed in sagittal sections, in lamina I and II, a dense band of Aδ-HTMR terminals and fibers was observed (Figure 6B). In contrast, in the deep dorsal horn, clusters of Aδ-HTMR terminals appeared discontinuous, potentially converging on sparsely distributed synaptic partners. This clustered pattern of deep Aδ-HTMR terminals was especially pronounced in the *Smr2^+^* population, perhaps reflecting distinguishing central connectivity of this Aδ-HTMR population. These genetic labeling experiments confirm and expand on earlier insights into Aδ-HTMR central anatomy achieved with single-axon reconstructions of physiologically defined neurons^17^.

**Figure 6.**
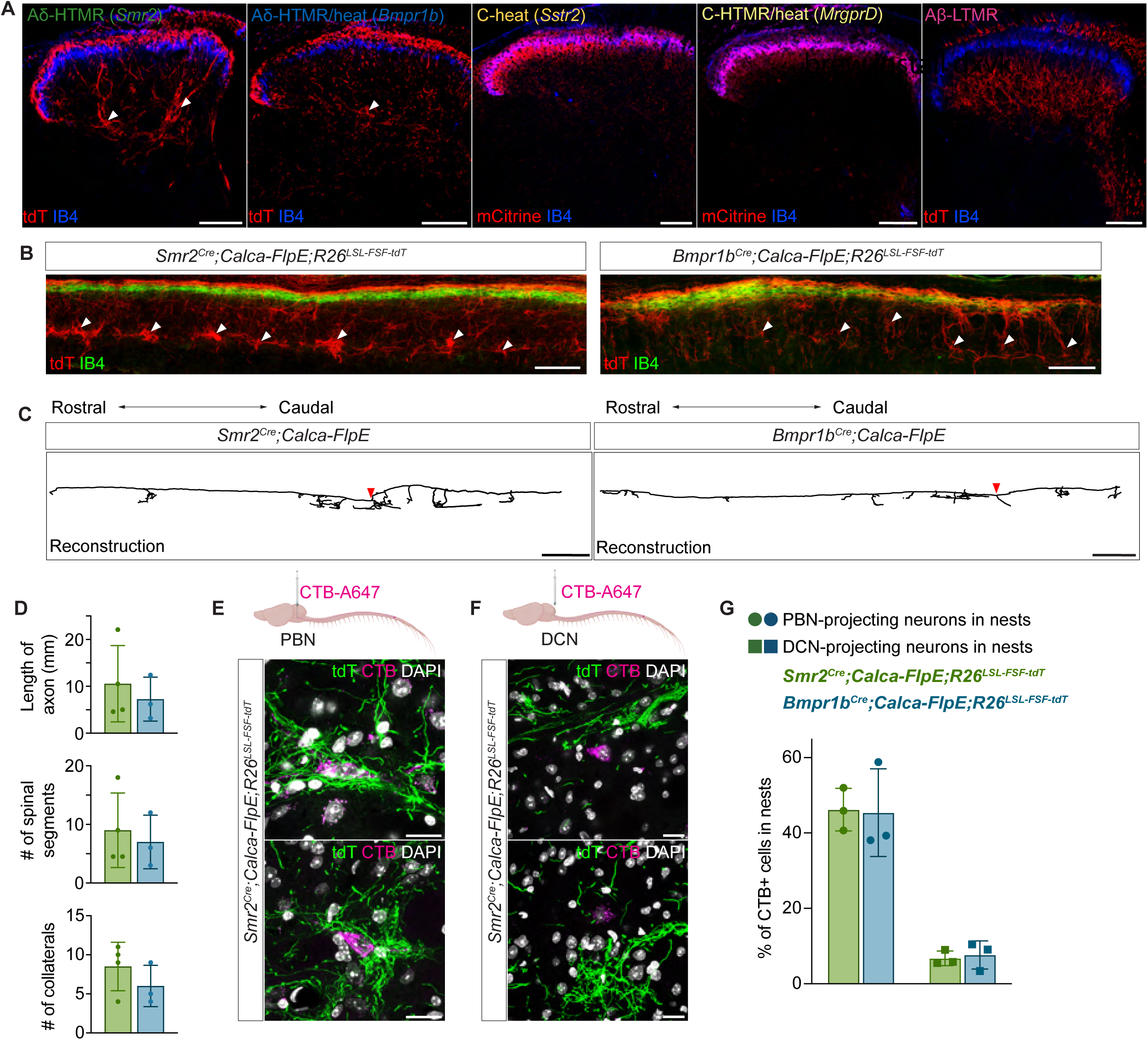
Aδ-HTMRs uniquely terminate across several laminae of the spinal cord dorsal horn and form collaterals that span multiple spinal segments. (A) Aδ-HTMR spinal cord termination patterns are unique among DRG sensory neuron types. Representative confocal images of the two Aδ-HTMR (*Smr2^Cre^; Calca-FlpE; R26^LSL-FSF-tdT^* and *Bmpr1b^Cre^; Calca-FlpE; R26^LSL-FSF-tdT^*), C-heat thermoreceptor (*Sstr2^CreER^; R26^LSL-ReaChR::mCitrine^*), C-HTMR/heat receptor (*MrgprD^CreER^; R26^LSL-ReaChR::mCitrine^*), and Aβ RA-LTMR (*TrkB^CreER^; Avil^FlpO^; R26^LSL-FSF-tdT^*) axons in the dorsal horn, taken from transverse sections of lumbar spinal cord. Arrowheads point at clusters of Aδ-HTMR axons in the deep dorsal horn. Scale bars are 100 µm. (B) Both Aδ-HTMR subtypes exhibit discontinuous terminals in the deep dorsal horn. Representative confocal images of *Smr2^Cre^; Calca-FlpE; R26^LSL-FSF-tdT^* and *Bmpr1b^Cre^; Calca-FlpE; R26^LSL-FSF-tdT^* spinal cord sagittal sections. Arrowheads point at clusters of Aδ-HTMR axons in the deep dorsal horn. Scale bars are 200 µm. (C) Aδ-HTMRs have long axons with many collaterals that span multiple segments of the spinal cord. Representative reconstructed axons of sparsely labeled Aδ-HTMRs. Red arrowheads indicate points of entry into the spinal cord. Scale bars are 1 mm. (D) Quantification of Aδ-HTMR central axon length along the rostrocaudal axis, number of spinal segments spanned, and number of collaterals (mean ± SD), related to examples shown in (C). Each dot represents an axon (n = 4 for *Smr2^Cre^*-labeled neurons and 3 for *Bmpr1b^Cre^*-labeled neurons). (E-F) Aδ-HTMRs form “nests” around antenna cell projection neurons of the anterolateral tract (ALT) (E) but not postsynaptic dorsal column neurons (PSDCs) (F). Representative confocal images from *Smr2^Cre^; Calca-FlpE; R26^LSL-FSF-tdT^* and *Bmpr1b^Cre^; Calca-FlpE; R26^LSL-FSF-tdT^* spinal cord sections. The projection neurons are retrogradely labeled using injections of the tracer CTB-A647 into the PBN (E) and DCN (F). Scale bars are 20 µm. (G) Percentage (mean ± SD) of CTB-labeled cells found within Aδ-HTMR nests. Quantification related to examples shown in (E-F). Each dot represents an animal (n = 3 for all groups).

We also performed sparse genetic labeling experiments to determine the spinal cord innervation patterns of individual Aδ-HTMRs (Figure 6C). Unlike Aδ-LTMRs, C-LTMRs, and other C-fiber neuron types, which form a single terminally branched arbor in a rostrocaudally restricted manner^59,64^, individual Aδ-HTMRs produce multiple collateral arbors (5+) along the rostrocaudal axis, reminiscent of all Aβ-LTMR subtypes and proprioceptors^59^ (Figure 6C, D). Additionally, axons of individual neurons were observed in both the superficial and deep lamina. These distinguishing features of Aδ-HTMRs suggest that they engage distinct postsynaptic partners along the rostro-caudal and dorsal-ventral axis of the spinal cord.

In both transverse and sagittal sections, the deep projections of Aδ-HTMRs appeared to envelop one or several cell bodies (Figure 6A, B), and this was especially apparent when counterstained with NeuN (Figure S7E). These striking anatomical structures, which we refer to as deep dorsal horn Aδ-HTMR “nests”, may hold clues to the identity of central mediators of the unique behaviors evoked by Aδ-HTMRs. Therefore, we next sought to identify spinal cord neurons that reside within these deep dorsal horn Aδ-HTMR nests. There are several different neuronal populations in this laminar location, including two projection neuron types with large cell bodies – antenna cells of the anterolateral tract (ALT) that project to the parabrachial nucleus (PBN) and other brain targets, and post-synaptic dorsal column neurons (PSDCs) that project to the dorsal column nuclei (DCN)^63,65^. To ask whether either or both projection neuron types are localized within Aδ-HTMR nests, chrolera toxin B (CTB) retrograde tracer was injected bilaterally into either the PBN to label antenna cells or the DCN to label PSDCs (Figure 6E, F). We found that Aδ-HTMRs form nests that surround spinoparabrachial antenna cells but rarely PSDCs (Figure 6G). Since antenna cells are sparse^65^ and Aδ-HTMR nests often envelop more than one cell body (Figure S7E), we suspect that, in addition to antenna cells, these nests surround additional unknown neuronal populations. It is also noteworthy that the superficial Aδ-HTMR terminals reside in close proximity to lamina I spinoparabrachial neurons (Figure S7F).

### Aδ-HTMRs directly synapse onto distinct ALT projection neuron populations

Because of the close proximity of Aδ-HTMR terminals to both superficial and deep ALT projection neurons, we hypothesized that Aδ-HTMRs form monosynaptic connections upon them. To test this idea, we performed whole-cell patch clamp electrophysiological recordings of retrogradely labeled ALT neurons in spinal cord slices while optically activating ReaChR-expressing *Smr2^Cre^*- and *Bmpr1b^Cre^*-labeled Aδ-HTMRs (Figure 7A, B). These experiments revealed that 100% (13 out of 13) of recorded antenna cells received strong monosynaptic inputs from genetically labeled Aδ-HTMRs (Figure 7C). Post hoc histological analysis of the biocytin-filled antenna cells revealed CGRP^+^ axons surrounding both the cell body and the dorsally directed dendrites of antenna cells (Figure 7D). In the superficial dorsal horn, 10 of 14 recorded spinoparabrachial neurons received Aδ-HTMR input which proved to be monosynaptic (Figure 7C). This heterogeneity of superficial spinoparabrachial neuron responses may reflect the presence of distinct ALT populations in the superficial dorsal horn^66-70^. Overall, these findings show that Aδ-HTMRs have highly unique central projections that form strong monosynaptic connections onto both superficial and deep dorsal horn ALT projection neurons that extend to the PBN and likely other target regions of the brain.

**Figure 7.**
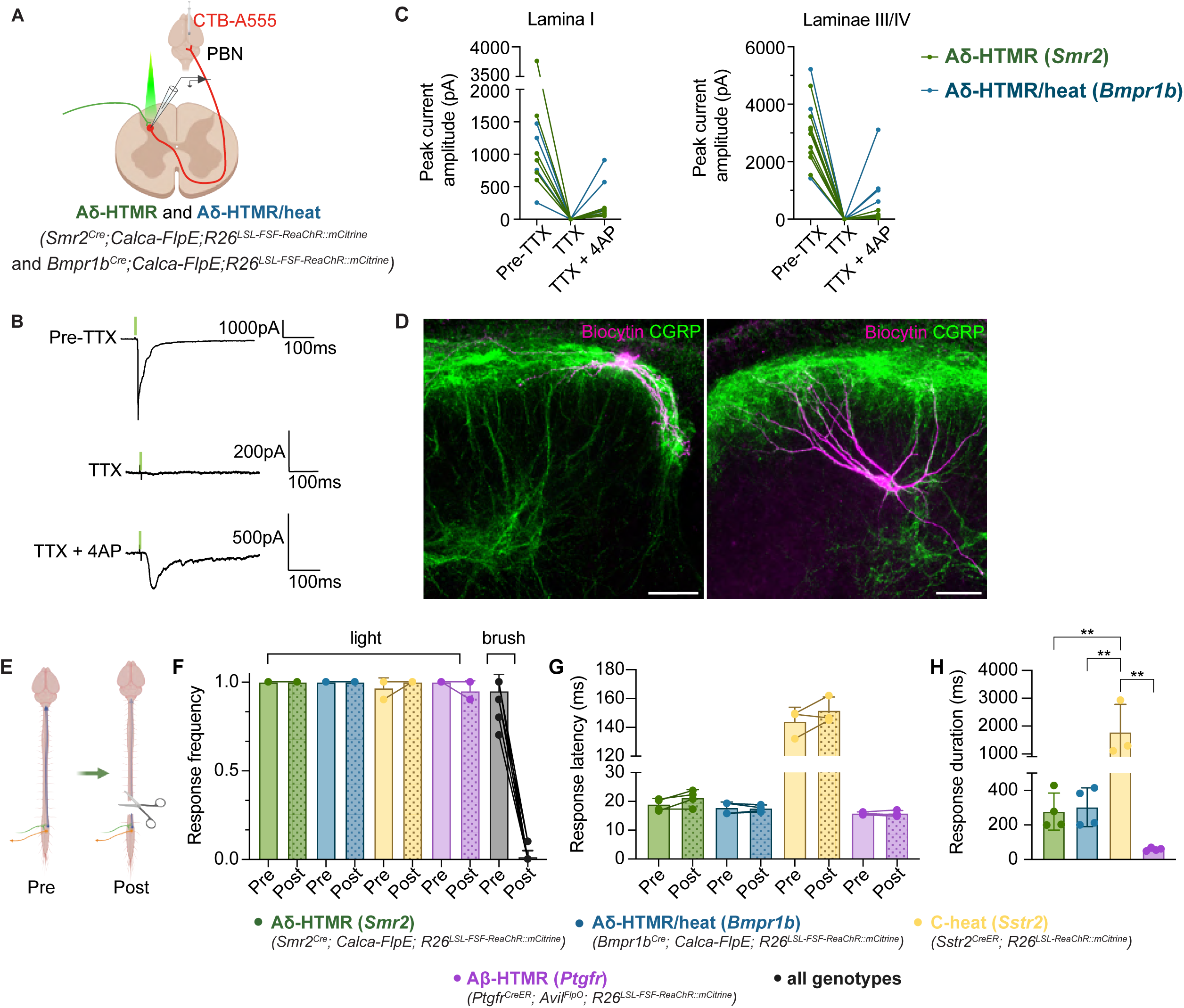
Aδ-HTMRs engage both a supraspinal circuit and a local reflexive circuit. (A) Whole-cell patch clamp recordings of retrogradely labeled ALT projections neurons with optical stimulation of Aδ-HTMR axons in *Smr2^Cre^; Calca-FlpE; R26^LSL-FSF-ReaChR::mCitrine^* and *Bmpr1b^Cre^; Calca-FlpE; R26^LSL-FSF-ReaChR::mCitrine^* spinal cord sections (300 µm). (B) Aδ-HTMRs form strong, monosynaptic connections onto ALT projection neurons. Representative traces of light-activated currents in a lamina III-IV ALT projection neuron. (C) Peak current amplitude of light-activated currents in superficial (lamina I) and deep (laminae III-IV) ALT projection neurons. Each dot represents a recording from one cell (n = 6 cells across 4 *Smr2^Cre^; Calca-FlpE; R26^LSL-FSF-ReaChR::mCitrine^* animals and 4 cells across 4 *Bmpr1b^Cre^; Calca-FlpE; R26^LSL-FSF-ReaChR::mCitrine^* animals for superficial cells; n = 9 cells across 4 *Smr2^Cre^; Calca-FlpE; R26^LSL-FSF-ReaChR::mCitrine^* animals and 4 cells across 4 *Bmpr1b^Cre^; Calca-FlpE; R26^LSL-FSF-ReaChR::mCitrine^* animals for deep cells). Importantly, of 13 deep cells recorded, all exhibited substantial light-evoked EPSCs that persisted upon TTX + 4-AP application. On the other hand, of 14 superficial cells recorded, four exhibited only weak light-evoked EPSCs (two per genotype); the latter four cells are not plotted. (D) Representative images of biocytin-filled superficial (left) and deep antenna (right) spinoparabrachial neurons. Cells were filled at the end of the recordings summarized in (A-C). Scale bars are 50 µm. (E) Schematic of spinal transection experiment. The spinal cord was cut above spinal segment T9, and mice were allowed to recover for 6 hours before testing. (F) Optical activation of Aδ-HTMRs promoted hindpaw withdrawal in spinalized animals, while brush-evoked responses were lost. 5-ms light pulses and brush stimuli were delivered to the hindpaws of *Smr2^Cre^; Calca-FlpE; R26^LSL-FSF-ReaChR::mCitrine^*, *Bmpr1b^Cre^; Calca-FlpE; R26^LSL-FSF-ReaChR::mCitrine^*, *Sstr2^CreER^; R26^LSL-ReaChR::mCitrine^*, and *Ptgfr^CreER^; Avil^FlpO^; R26^LSL-FSF-ReaChR::mCitrine^* animals before (Pre) and after (Post) spinalization. See Videos S3 and S4 for representative examples of Aδ-HTMR and C-heat thermoreceptor-driven behaviors after spinalization. Each dot represents an animal (n = 4, 4, 4, 14 from left to right). (G) Spinal transection does not alter the light-evoked response latency. Each dot represents an animal, the same as (F). (H) Optical activation of physiologically distinct sensory neuron types evokes reflexive responses of varying duration. See Video S5 for an example of prolonged behavioral response to activation of C-heat thermoreceptors. The assay was performed in spinalized animals. Each dot represents an animal (n = 4, 4, 3, 4 from left to right, *\*\*p* ≤ 0.01, one-way ANOVA with Tukey’s multiple comparisons post-hoc test).

### Aδ-HTMRs are the primary drivers of an ultrafast, spinally-mediated limb withdrawal reflex

Motivated by the observation that optical activation of ReaChR expressing Aδ-HTMRs evokes invariable limb withdrawal with a short, 20-25 ms latency, we next asked whether this response is mediated by an ascending signal or a local spinal reflex. For this, the spinal cord of animals expressing ReaChR in their Aδ-HTMRs was transected at thoracic segment 9 (T9), leaving a potential local lumbar spinal reflex circuit intact (Figure 7E). A complete loss of motor responses to brushing the hindpaws was observed in the spinalized animals. However, when light pulses were applied to the paw, a fast withdrawal and paw shaking behavior was consistently observed, although, unsurprisingly, the response did not involve whole-body movement as compared to control mice (Figure 7F, G, Video S3). This finding indicates that Aδ-HTMR activation is sufficient to drive short-latency limb withdrawal and paw shaking through a spinal reflex. We also observed that activating *Sstr2^CreER^*-labeled C-heat thermoreceptors and *Ptgfr^CreER^*-labeled Aβ-HTMRs in spinalized animals evoked a reflexive paw movement (Figure 7F, G). Interestingly, the movement pattern, magnitude, and duration differed from those evoked by Aδ-HTMR activation, suggesting that these physiologically distinct sensory neurons engage distinct spinal reflex motor circuits (Figure 7H, Videos S4, S5). Importantly, although Aβ-HTMRs evoked a modest reflexive paw withdrawal when optically activated, these neurons are not classified as nociceptors based on behavioral and physiological criteria, underscoring the notion that fast reflexes cannot be equated to nociception or pain and are mediated by distinct circuits. Thus, Aδ-HTMRs engage both fast local spinal circuits and slower ascending pathways to drive nocifensive behaviors and aversion.

## DISCUSSION

We conducted an optogenetic activation screen of somatosensory neuron types in naive healthy mice to identify genetically labeled nociceptor populations based on a physiological and behavioral definition rooted in early work of Sherrington^27^ and Perl^18^. Two Aδ-HTMR populations and a C-heat thermoreceptor population emerged from our analyses as nociceptors. It is possible that a 5-ms light pulse was not an optimal stimulus for all studied sensory neuron populations and this is a limitation of our study. Nevertheless, the stimulus was sufficient to evoke electrophysiological responses in the spinal cord, and the differential responses to this stimulus might reflect different levels of nociceptive circuitry engagement by the afferents. It is also revealing that the three nociceptor populations identified in our behavioral screen had mechanical and thermal thresholds that were so high that continued application of such stimuli would result in tissue damage^23^. Importantly, our findings highlight that other commonly cited sensory neuron features are poor predictors of nociceptor identity, including cell body diameter, broad molecular markers such as Na_v_1.8 expression or IB4 binding, and an “HTMR” designation. It is worth emphasizing that we do not argue that nociceptors are the only sensory neurons capable of driving pain. Instead, we propose that the term “nociceptor” is most useful when studying acute nociceptive pain in healthy animals. Indeed, activation of LTMRs and other sensory neuron types may evoke nocifensive reflexes and the perception of pain following nerve injury, upon inflammation, and in other disease states^26,71,72^.

Here, we characterized and compared three distinct A-fiber HTMR populations, two of which are nociceptors. Given that several nomenclature systems are used in the field^19,21,23,28,53^, we confirmed *in situ* that the *Smr2^Cre^*-labeled Aδ-HTMRs correspond to the PEP2 or A-PEP.CHRNA7 transcriptional clusters and *Bmpr1b^Cre^*-labeled Aδ-HTMRs to the PEP3 or A-PEP.KIT transcriptional clusters^19,53,73,74^. More work is needed to characterize the *Ptgfr^CreER^*-labeled Aβ-HTMRs but they likely correspond to the Aβ-LTMR.NSG, PEP2.2, Aβ Field-LTMR, and Ntrk3^high^+/S100a16 clusters^19,53,73,74^. Importantly, while mouse Aδ-HTMRs are nociceptors and Aβ-HTMRs are not, a conduction velocity distinction is likely to be less predictive of function in humans^14,19,75^. In the mouse, we find that the three genetically labeled A-fiber HTMRs have distinct but overlapping physiological and morphological properties. However, their central anatomy and, presumably, synaptic partners are distinct. While Aβ-HTMRs terminate broadly in the deep dorsal and project to the DCN, Aδ-HTMRs terminate within both the superficial and deep dorsal horn where they form “nests” around and synapse onto the ALT projection neurons. These findings emphasize the significance of afferents’ central connectivity to their function.

Both Aδ-HTMR populations drive a remarkably fast local spinal reflex, and establishing genetic access to first-order neurons in this nociceptive withdrawal reflex circuit can now enable identifying the cellular components of this long-recognized but unresolved circuit^27,76^. Aδ-HTMRs also provide strong monosynaptic inputs to ALT projection neurons across dorsal horn laminae. The role of the ALT in pain processing is well established, although most work in the spinal cord has focused on the ALT projection neurons of the superficial dorsal horn^15,67,68,70,77,78^. The antenna cells of the deep dorsal horn have distinct intrinsic electrophysiological properties^79^, and they are poised to receive different synaptic inputs compared to their superficial counterparts based on their laminar location. Indeed, Aδ-HTMRs are unique in forming nests around antenna cell bodies, presumably enabling their remarkably strong synaptic connections. How Aδ-HTMR signals are integrated with other sensory inputs to shape ALT projection neuron firing patterns is an exciting area of future research.

Aδ-HTMRs densely innervate skin and other organs. While further analysis of the properties of these neurons in non-cutaneous tissues is warranted, in long bones^60^, kidneys^80^, bladder^81^, cranial meninges^82^, and teeth^83^, Aδ fibers, and Aδ-HTMRs in particular, have been electrophysiologically identified, making it likely that, at least in those tissues, *Smr2^Cre^*- and *Bmpr1b^Cre^*-labeled neurons have physiological properties and functions similar to their cutaneous counterparts. Indeed, in the distal colon, *Bmpr1b^Cre^*-labeled neurons function as Aδ-HTMRs^51^. Across organs, Aδ-HTMRs may contribute both to normal tissue homeostasis and protection and to excessive and maladaptive pain in disease, and a genetic handle on these neurons will allow for deeper understanding of their contributions.

In conclusion, work presented here establishes two genetically labeled Aδ-HTMRs as myelinated nociceptors with unique physiological, morphological, and synaptic features distinguishing them from C-fiber nociceptors and Aβ-HTMRs. These findings open avenues for advancing our understanding of the neurobiological underpinnings of pain in health and disease and the development of new analgesics.

## Resource availability

### Lead contact

Further information and requests for resources should be directed to the lead contact, David D. Ginty.

### Materials availability

Requests for *Ptgfr^CreER^* mouse line generated in this study should be directed to and will be fulfilled by the lead contact upon request.

### Data and code availability

Data reported in this paper are available from the lead contact upon request.

## Acknowledgements

We thank Andrew Shuster, Xiangsunze Zeng, David Paul, Hankyul Kwak, Aindrila Saha, Anna Lebedeva, and Kitwa Ng for comments on the manuscript and all members of the Ginty lab for helpful discussions. We thank Caiying Guo at the Janelia Gene Targeting and Transgenic Facility for generating the *Pgtfr^CreER^* mouse line. We thank Lijun Qi for his guidance on *in vivo* calcium imaging and the code used for analysis. We thank Hankyul Kwak and Zoe Sarafis for guidance on conduction velocity measurement experiments. We thank Andrew J. Todd for helpful discussions on anterolateral tract and antenna cells. Schematics were created in BioRender under Harvard University license. This work was supported by NIH grants R35 5R35NS097344-05 (D.D.G.), R35 5R35NS132196-02 (D.D.G.), 1RO1AT011447 (D.D.G.), R01AR064251 (A.M.M.), R01AR060364 (A.M.M.), P30AR079206 (A.M.M.), UC2AR082186 (A.M.M.), R01AR077019 (R.E.M.), the Yang Tan Collective at Harvard Medical School (D.D.G.), K. Lisa Yang Brain Body Center at Harvard Medical School (D.D.G.), NEI P30 Core Grant for Vision Research no. EY012196 (O.M.), Mahoney Postdoctoral Fellowship (J.T.), Gordon Postdoctoral Fellowship (J.T.), William Randolph Hearst Fund (R.I.M.-G.), and Harvard Medical School Shapiro predoctoral fellowship (K.L.). D.D.G. is an investigator of the Howard Hughes Medical Institute (HHMI). This article is subject to HHMI’s Open Access to Publications policy. HHMI lab heads have previously granted a nonexclusive CC BY 4.0 license to the public and a sublicensable license to HHMI in their research articles. Pursuant to those licenses, the author-accepted manuscript of this article can be made freely available under a CC BY 4.0 license immediately upon publication.

## Author contributions

K.L. and D.D.G. conceived the study. K.L. performed all behavioral experiments, except for the real-time place preference. M.M.D. developed the real-time place preference assay with help from O.M. M.M.D. performed the real-time place preference and data analysis for open field, balance beam, temperature and texture preference assays. K.L. developed the pin-prick assay with help from J.T. K.L. performed *in vivo* loose patch electrophysiology with guidance from A.J.E. K.L. performed the calcium imaging experiments. K.N. and K.L. performed immunohistological experiments. N.S. provided plasmids used in the sparse labeling experiments, and K.N. performed the experiments. K.L., K.N., and J.C. performed RNAScope™ and related data analyses. R.S. performed immunostaining of the meninges. F.C.K., S.F., and A.M.O. performed immunostaining and imaging of bones and joints, with guidance from A.M.M. and R.E.M. J.L. performed slice electrophysiology and spinal cord MEA recordings. R.I.M.-G. provided guidance on slice electrophysiology experiments. J.T., I.A., and B.P.L. characterized the *Ptgfr^CreER^* mouse line and the anatomical and physiological properties of Aβ-HTMRs. L.G. collected human skin biopsies. K.L. and D.D.G. wrote the manuscript with input from all authors.

## Supplemental Figure Legends

**Figure S1.**
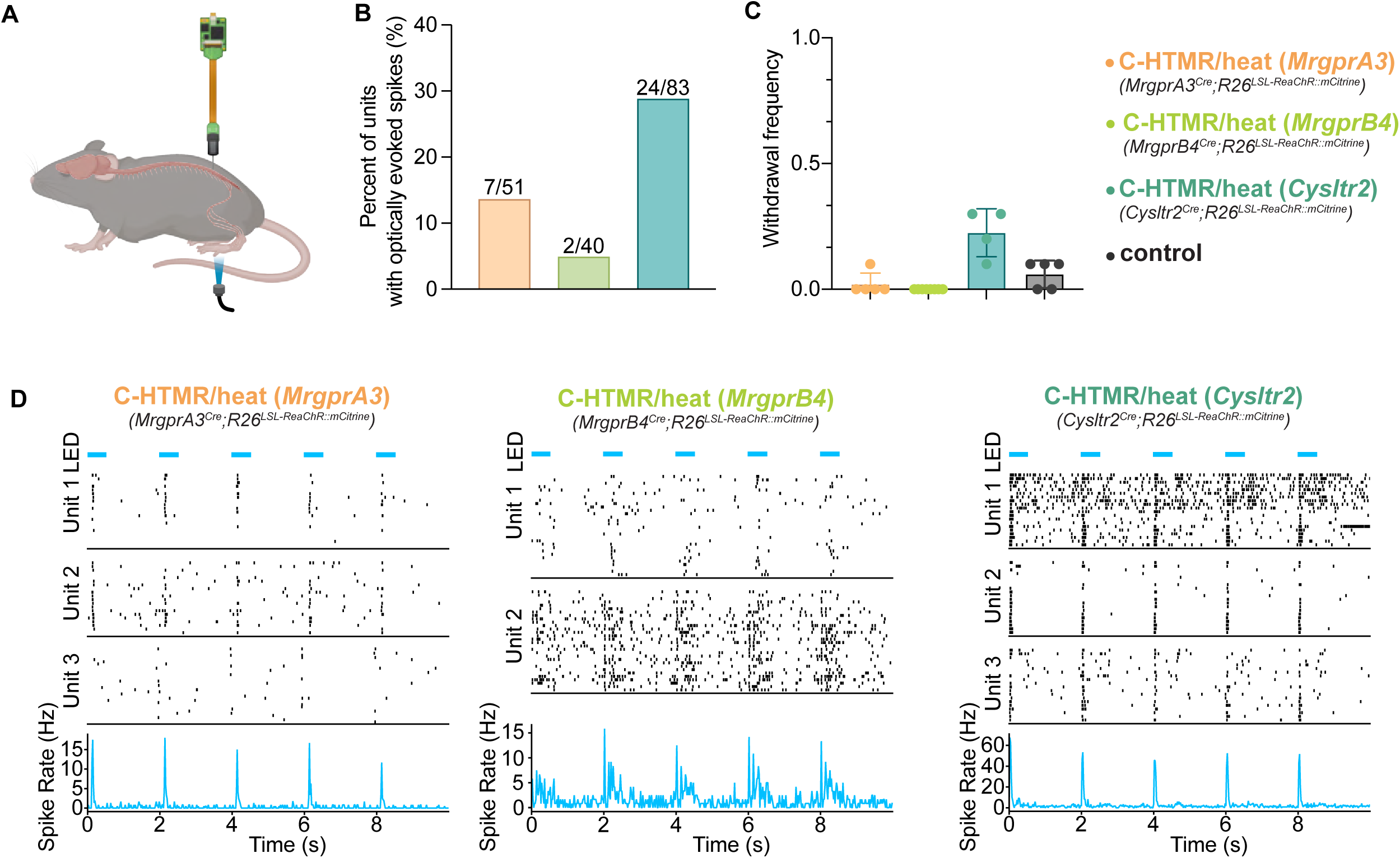
Optically evoked neural activity in the spinal cord in the absence of an observable behavioral response, related to Figure 1. (A) *In vivo* multielectrode array recordings from lumbar spinal cord neurons. Sensory neurons are stimulated with light pulses on the glabrous skin of animals expressing ReaChR in different DRG populations. (B) Percentage of recorded units with light-evoked responses. Units are summed across multiple animals (n = 2, 1, 3 animals from left to right). (C) Light stimulation of the paw does not evoke reliable withdrawal. Average (mean ± SD) withdrawal frequency plotted. Longer stimulation (500 ms 0.2 Hz pulses for *MrgprA3^Cre^; R26^LSL-ReaChR^*, *Cysltr2^Cre^; R26^LSL-ReaChR^* and littermate controls that lack a driver or ReaChR allele, continuous light for *MrgprB4^Cre^; R26^LSL-ReaChR^*) does not increase withdrawal frequency (see Figure 1C for 5-ms pulse stimulation). Each dot represents an animal (n = 5, 8, 4, 5 from left to right). (D) Example spike rasters and peristimulus time histograms (PSTHs) from three representative units from *MrgprA3^Cre^; R26^LSL-ReaChR^* and *Cysltr2^Cre^; R26^LSL-ReaChR^* animals, 2 units from an *MrgprB4^Cre^; R26^LSL-ReaChR^* animal. Stimulus is 500 ms light pulse.

**Figure S2.**
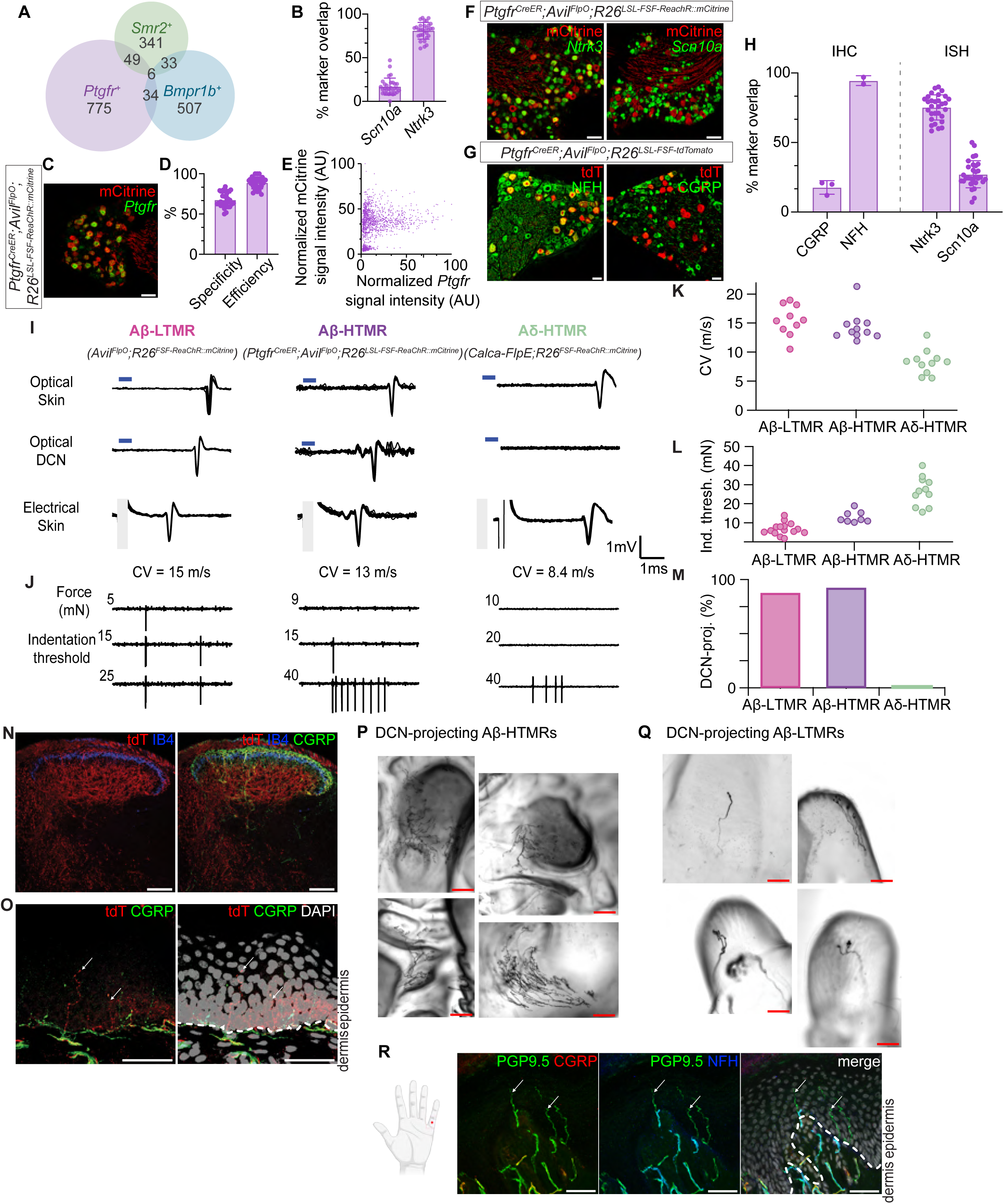
Characterization of Aβ-HTMRs labeled by the *Ptgfr^CreER^* mouse line, related to Figure 1. (A) Quantification of overlap between cell type markers of three putative A-fiber HTMR populations: *Smr2*, *Bmpr1b*, and *Ptgfr*. Analysis performed based on *in situ* hybridization. Cells with robust signal only were considered “positive” for a given marker. The numbers represent cells. Total number of cells analyzed is 1745 across 30 DRG sections from 3 animals. (B) Average percentage (mean ± SD) of *Ptgfr*^+^ neurons that co-express a given marker as accessed by *in situ* hybridization. Each dot represents a DRG section (n = 30 sections across 3 animals for bor both markers). (C) *In situ* hybridization and immunostaining of lumbar DRG sections from mice with genetically labeled *Ptgfr^+^* neurons. Scale bar is 100 µm. (D) Quantification of specificity and efficiency of the *Ptgfr^CreER^*-based labeling strategy. Average (mean ± SD) values displayed as bars. Specificity is defined as the percentage of reporter-positive cells that are also positive for the cell type marker *Ptgfr*. Efficiency is defined as the percentage of marker-positive cells that are also positive for the reporter. Each dot represents a DRG section (n = 30 sections across 3 animals, same sections analyzed for both quantifications). (E) The relationship between reporter (mCitrine) and cell type marker (*Ptgfr*) expression. Expression mean gray values are normalized by the maximum value of the respective signal. Each dot represents a cell (n = 1456 cells across 30 DRG sections from 3 animals). (F) *In situ* hybridization and immunostaining of lumbar DRG sections from mice with genetically labeled *Ptgfr^+^* neurons. Scale bars are 100 µm. (G) Immunostaining of lumbar DRG sections from mice with genetically labeled *Ptgfr^+^* neurons. Note the large diameter of reporter^+^ cells. Scale bars are 50 µm. (H) Average percentage (mean ± SD) of reporter^+^ neurons that co-express a given marker, as accessed by immunohistochemistry (left) and *in situ* hybridization (right). For immunohistochemistry, each dot represents an average of several DRG sections from a single animal (n = 3, 2 mice from left to right). For *in situ* hybridization, each dot represents a DRG section (n = 30 sections across 3 animals for both markers). (I) Comparative characterization of Aβ-LTMRs (labeled broadly with *Avil^FlpO^*, units selected based on fast conduction velocity and low mechanical threshold), Aβ-HTMRs (labeled with *Ptgfr^CreER^*), and Aδ-HTMRs (labeled broadly with *Calca-FlpE*, only signals from A-fiber neurons are picked up with this technique) using *in vivo* electrophysiological recordings with sharp glass electrodes in anesthetized mice. Representative voltage responses to optical stimulation of the skin or the dorsal column nuclei (DCN), and electrical stimulation of the skin. (J) Representative responses of the three sensory neuron populations to mechanical stimulation of the glabrous paw skin with precise indentation using a 200 μm probe tip. (K) Conduction velocity measurements for the three sensory neuron populations. Each dot represents a single unit (n = 9 units from 6 animals for Aβ-LTMRs, 11 units from 5 animals for Aβ-HTMRs, 11 units from 4 animals for Aδ-HTMRs). (L) Indentation force threshold measurement for the three sensory neuron populations. Each dot represents a single unit (n = 15 units from 10 animals for Aβ-LTMRs, 8 units from 4 animals for Aβ-HTMRs, 11 units from 4 animals for Aδ-HTMRs). (M) Percentage of recorded units that have axons projecting to the DCN, determined by antidromic stimulation (7/8 units for Aβ-LTMRs, 12/13 units for Aβ-HTMRs, 0/9 units for Aδ-HTMRs). (N) Sections of spinal cord dorsal horn from a *Ptgfr^CreER^; Avil^FlpO^; R26^LSL-FSF-tdTomato^* animal. The genetically labeled Aβ-HTMRs terminate predominantly in the deep dorsal horn, below the IB4 band. The terminals are dense and diffuse, distinct from the clustered pattern of the CGRP^+^ axons. Scale bars are 100 µm. (O) Sections of glabrous paw skin from a *Ptgfr^CreER^; Avil^FlpO^; R26^LSL-FSF-tdTomato^* animal. The genetically labeled Aβ-HTMRs are CGRP^-^ and terminate in the epidermis. Scale bars are 50 µm. (P) Examples of whole-mount AP staining of peripheral arbors of DCN-projecting Aβ-HTMRs. AAV-Flp virus was injected into the DCN of *Ptgfr^CreER^; Tau^FSFiAP^* animals to achieve sparse labeling. Individual arbors are large compared to those in (K). Scale bars are 200 μm. (Q) Examples of whole-mount AP staining of peripheral arbors of DCN-projecting Aβ-LTMRs. AAV-Cre virus was injected into the DCN of *Brn3a^AP^* animals to achieve sparse labeling. LTMR identity of labeled neurons is inferred based on known morphology. Scale bars are 200 μm. (R) Sections of glabrous palm skin from a human donor. Arrows point to PGP9.5^+^CGRP^-^ endings in the epidermis that are NFH^+^ in the dermis, reflecting myelination. Scale bars are 50 µm.

**Figure S3.**
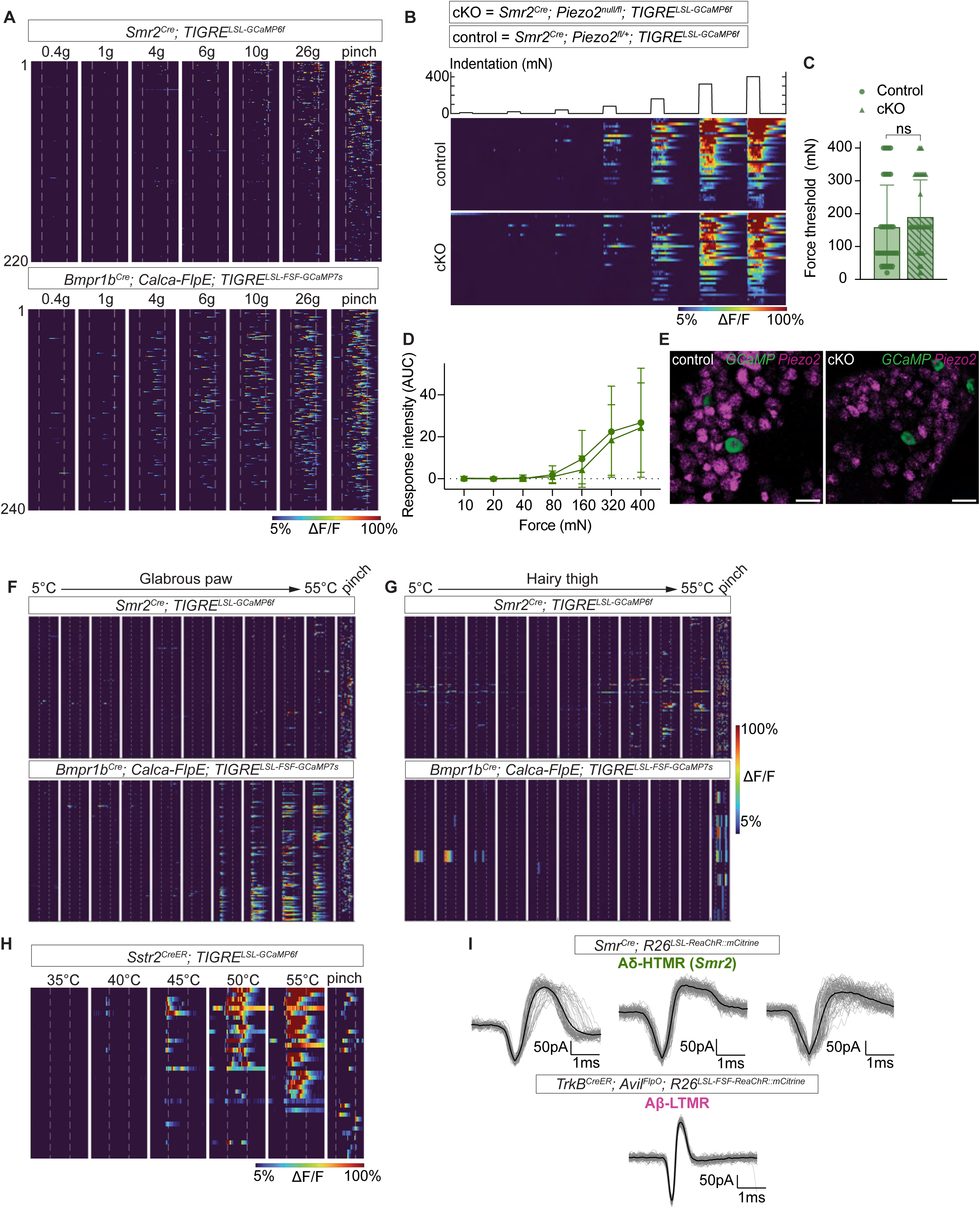
Additional physiological characterization of genetically labeled Aδ-HTMRs and C-heat thermoreceptors, related to Figure 2. (A) Calcium indicator responses of *Smr2^Cre^*- and the *Bmpr1b^Cre^*-labeled neurons to indentation of glabrous skin with von Frey filaments. While both populations only respond to relatively high forces, above 1 g, *Bmpr1b^Cre^*-labeled neurons are more sensitive than *Smr2^Cre^*-labeled neurons (n = 223 *Smr2^Cre^*-labeled neurons across 9 animals, 246 *Bmpr1b^Cre^*-labeled neurons across 3 animals). (B) Calcium responses of *Smr2^Cre^*-labeled neurons to controlled indentation of glabrous skin in Piezo2 conditional knockout (cKO) mice and littermate controls. *Smr2^Cre^; Piezo2^null/fl^; TIGRE^LSL-GCaMP6f^* mice were used as Piezo2 cKO, littermates with one floxed and one wild-type *Piezo2* allele were used as controls (n = 39 neurons across 5 animals for control, and 49 neurons across 6 animals for Piezo2 cKO). (C) Indentation force threshold (mean ± SD) of *Smr2^Cre^*-labeled neurons is unaffected by Piezo2 knockout. Each dot/triangle represents a cell (n same as in (B), Mann-Whitney test). (D) Indentation response intensity (mean ± SD) of *Smr2^Cre^*-labeled neurons is unaffected by Piezo2 knockout. A quantification of calcium responses shown in (B). Each dot/triangle represents an average intensity value from n cells (n same as in (B), permutation test on the mean difference, 10000 permutations). (E) *Piezo2* expression is very low in *GCaMP^+^ Smr2^Cre^*-labeled neurons compared to other DRG neurons in both Piezo2 cKO and control mice. Representative images of RNAScope™ on DRG sections. Scale bars are 50 µm. (F-G) Temperature responses of *Smr2^Cre^*- and the *Bmpr1b^Cre^*-labeled neurons vary by innervation target. Calcium responses of *Smr2^Cre^*- and the *Bmpr1b^Cre^*-labeled neurons to different temperatures presented to (F) glabrous paw and (G) hairy thigh skin. Pinch responses are also shown for comparison. The hairy skin-innervating *Smr2^Cre^*-labeled neurons are more heat-sensitive than their glabrous skin-innervating counterparts. On the other hand, the glabrous skin-innervating *Bmpr1b^Cre^*-labeled neurons are more heat-sensitive than their hairy skin-innervating counterparts (n = 110 glabrous skin-innervating and 126 hairy skin-innervating *Smr2^Cre^*-labeled neurons from 15 animals, 140 glabrous skin-innervating and 12 hairy skin-innervating *Bmpr1b^Cre^*-labeled neurons from 3 animals). (H) *Sstr2^CreER^*-labeled neurons are tuned to noxious heat. Calcium responses of *Sstr2^CreER^*-labeled neurons to select temperatures and pinch (n = 37 *Sstr2^CreER^*-labeled neurons from 3 animals). (I) Aδ-HTMRs have wide extracellular spike waveforms compared to Aβ-LTMRs. Example spike waveforms acquired with *in vivo* loose patch electrophysiology. Each waveform is from a different unit. The Aβ-LTMR example is replotted from Emanuel et al. 2021.

**Figure S4.**
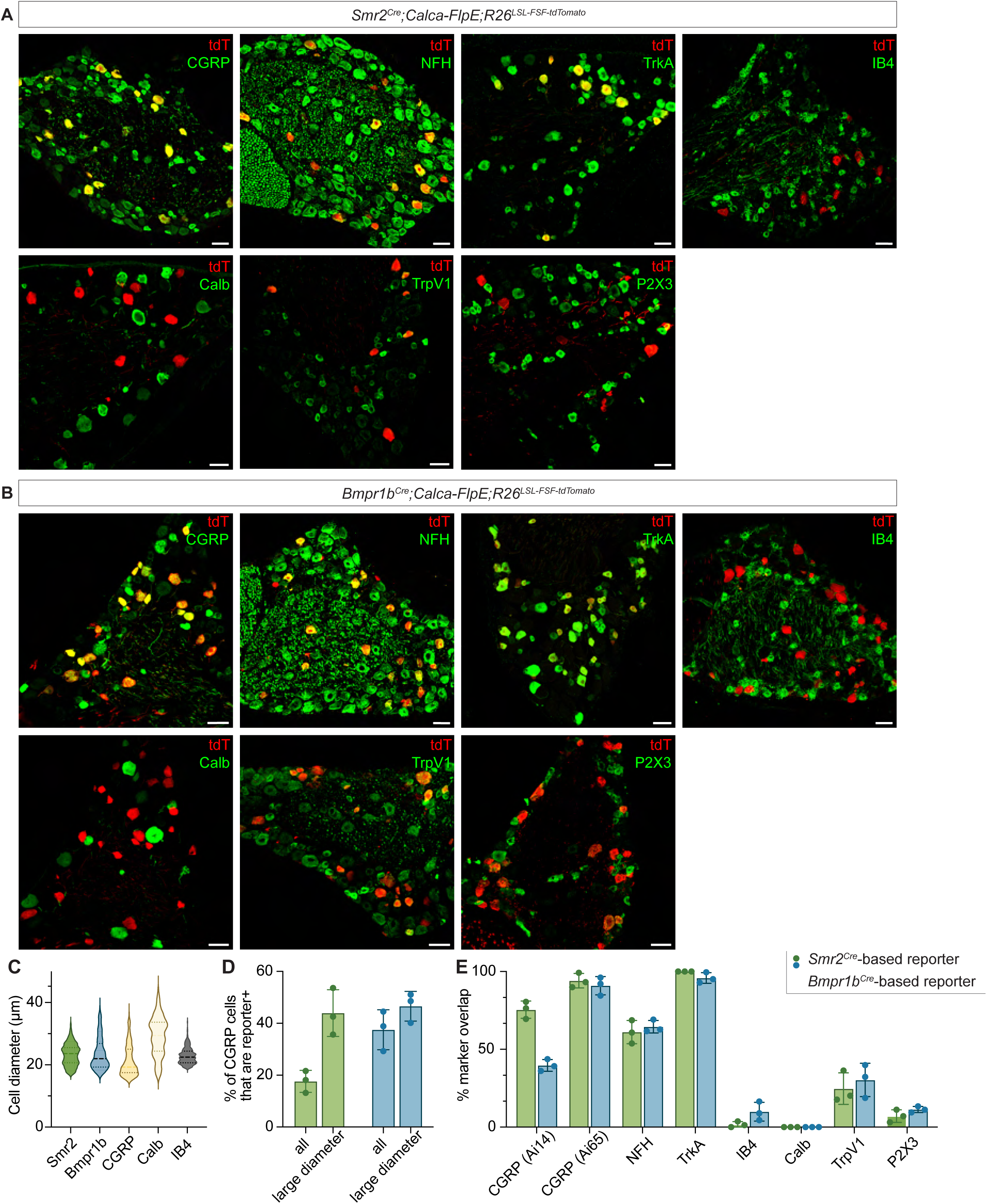
Molecular characterization of genetically labeled Aδ-HTMRs, related to Figure 2. (A-B) Immunostaining of lumbar DRG sections from mice with genetically labeled (A) *Smr2^+^* and (B) *Bmpr1b^+^* neurons. Scale bars are 50 μm. (C) *Smr2^Cre^*- and *Bmpr1b^Cre^*-labeled neurons have medium/large-diameter soma, larger than many other CGRP^+^ neurons. Diameters of reporter^+^ or marker^+^ soma are plotted (n = 237, 183, 369, 176, 580 cells across 2, 2, 1, 3, 3 animals from left to right). (D) Together, the *Smr2^Cre^*- and *Bmpr1b^Cre^*- labeled populations account for most large-diameter CGRP^+^ cells. Average percentage (mean ± SD) of all and large diameter CGRP^+^ cells that are *Smr2^Cre^*- or *Bmpr1b^Cre^*-labeled. Large diameter is defined as > 25 μm. Each dot represents an animal (n = 3 animals for all analyses). (E) Average percentage (mean ± SD) of reporter^+^ neurons that co-express a given marker. Each dot represents an average of 10 DRG sections from a single animal (n = 3 animals for all analyses).

**Figure S5.**
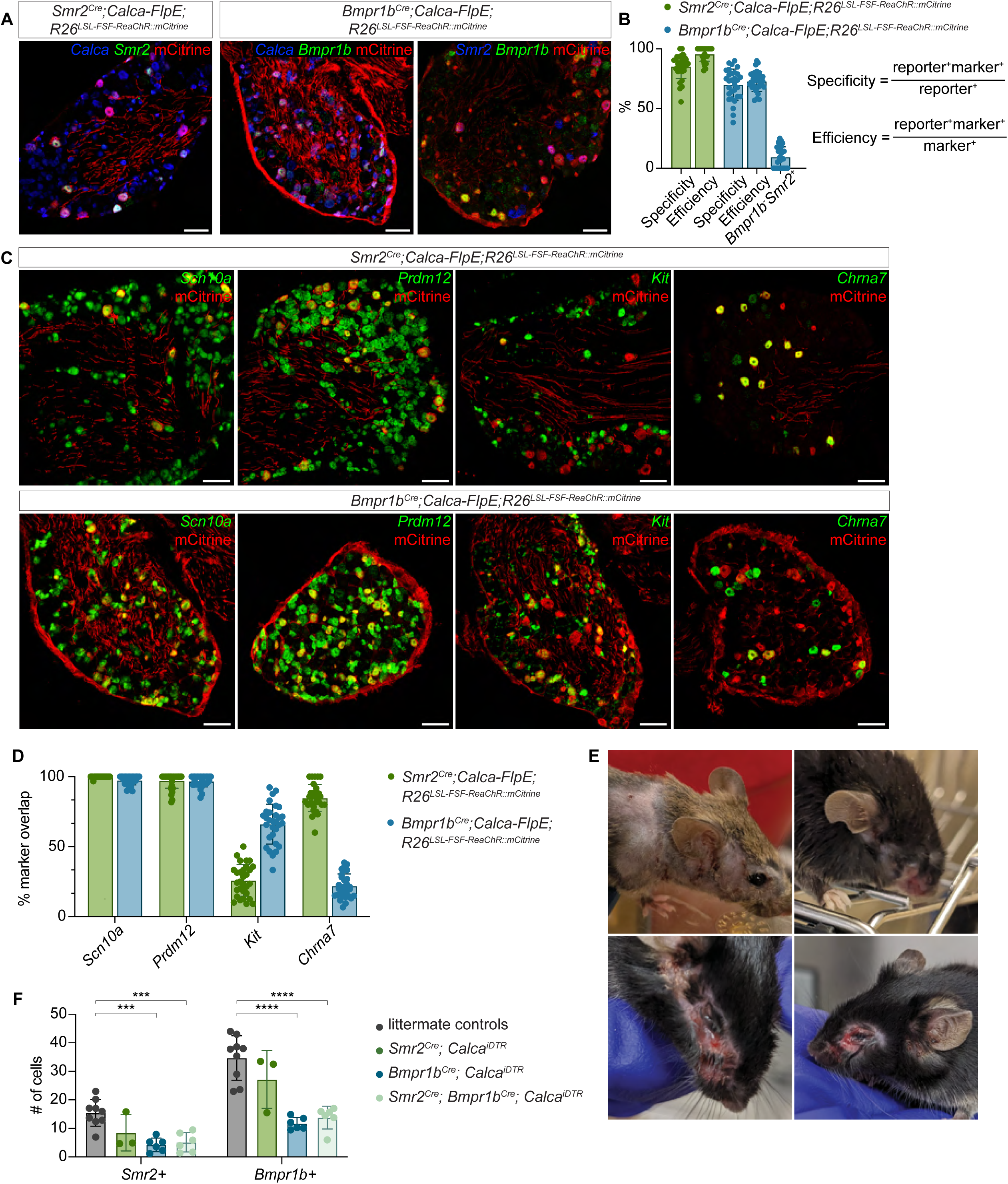
Further molecular characterization of Aδ-HTMRs and their chemogenetic ablation, related to Figures 2 and 3. (A) *In situ* hybridization and immunostaining of lumbar DRG sections from mice with genetically labeled *Smr2^+^* (first image) and *Bmpr1b^+^* (second and third images) neurons. Scale bars are 100 µm. (B) Quantification of specificity and efficiency of the *Smr2^Cre^; Calca-FlpE* and *Bmpr1b^Cre^; Calca-FlpE* based labeling strategies. Average (mean ± SD) values displayed as bars. Specificity and efficiency are defined as above. Marker-positive cells are those showing both *Smr2/Bmpr1b* and *Calca* expression. The rightmost bar quantifies the percentage of reporter-positive cells that were negative for *Bmpr1b* but positive for *Smr2*. Each dot represents a DRG section (n = 30 sections across 3 animals for all analyses). (C) *In situ* hybridization and immunostaining of lumbar DRG sections from mice with genetically labeled *Smr2^+^* (top row) and *Bmpr1b^+^* (bottom row) neurons. Scale bars are 100 µm. (D) Average percentage (mean ± SD) of reporter^+^ neurons that co-express a given marker. Each dot represents a DRG section (n = 30 sections across 3 animals for all markers). (E) Examples of facial wounds developed by mice whose Aδ-HTMRs were ablated. (F) Chemogenetic ablation of Aδ-HTMRs is ∼50% efficient. Average number (mean ± SD) of *Smr2^+^* and *Bmpr1b^+^* cells across experimental groups. Each dot represents an average of 10 DRG sections from a single animal (n = 9, 3, 6, 6 animals from left to right, *\*\*\*p* ≤ 0.001, *\*\*\*\*p* ≤ 0.0001, one-way ANOVA with Tukey’s multiple comparisons post-hoc test).

**Figure S6.**
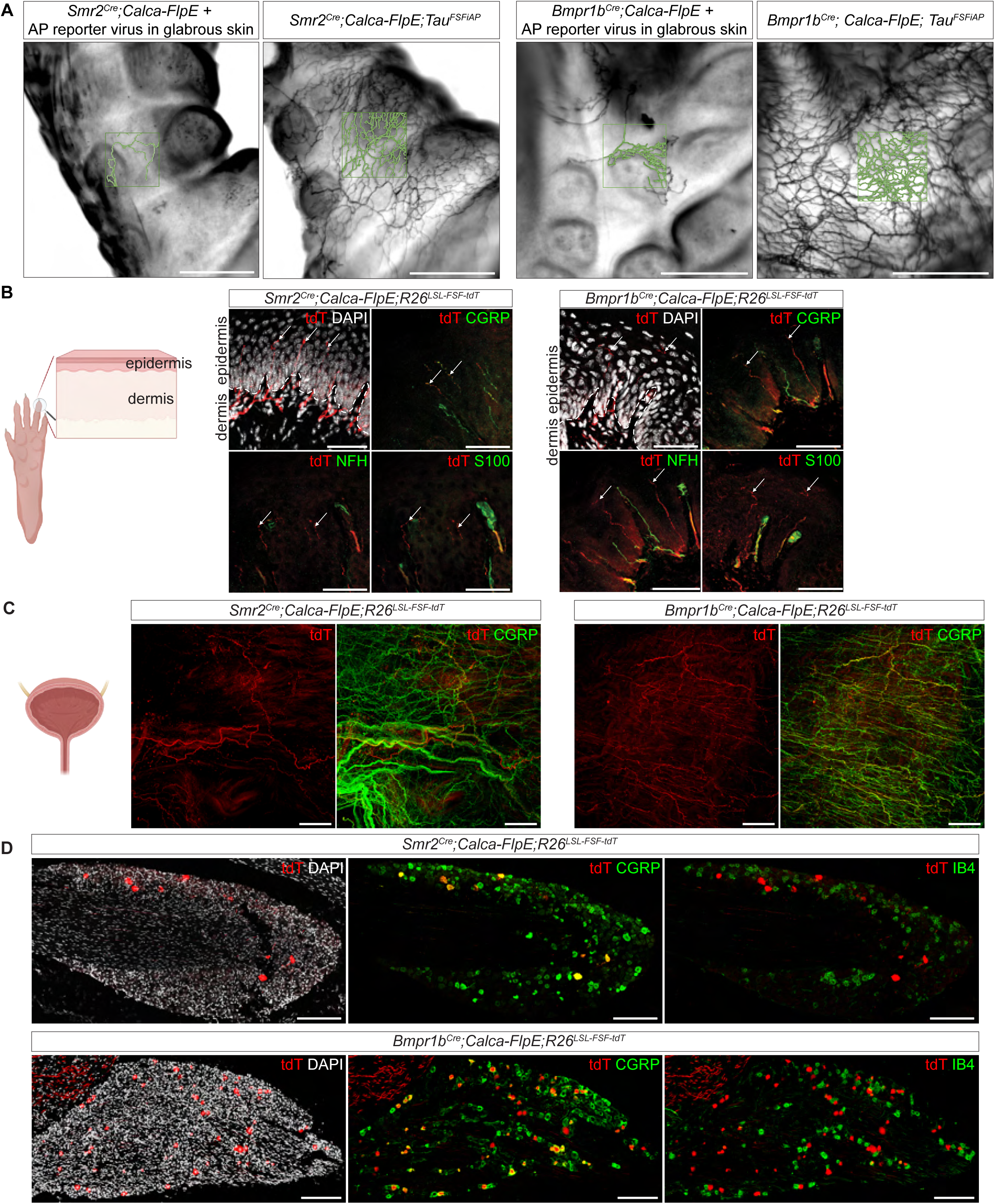
Peripheral morphology of genetically labeled Aδ-HTMRs, related to Figures 4 and 5. (A) Axons from multiple Aδ-HTMRs overlap to create a dense network of endings in glabrous skin. Representative image pairs exemplifying the process of overlap index quantification. In each pair of images, the left image is of sparsely labeled Aδ-HTMR axons, achieved with low-titer reporter virus injection (see Methods for details), and the right image is of densely labeled Aδ-HTMRs, achieved with constitutive genetic labeling. The green outlines are the axon reconstructions. Scale bars are 500 μm. (B) Genetically labeled Aδ-HTMRs penetrate the epidermis and form free nerve endings. Representative confocal images of *Smr2^Cre^; Calca-FlpE; R26^LSL-FSF-tdT^* and *Bmpr1b^Cre^; Calca-FlpE; R26^LSL-FSF-tdT^*glabrous skin sections. Reporter^+^ endings are pointed out using white arrows. Dashed lines indicate the border between the dermis and epidermis. Scale bars are 50 μm. (C) Genetically labeled Aδ-HTMRs innervate the walls of the urinary bladder. Representative confocal images from *Smr2^Cre^; Calca-FlpE; R26^LSL-FSF-tdT^* and *Bmpr1b^Cre^; Calca-FlpE; R26^LSL-FSF-tdT^* whole-mount stained bladders. The reporter signal represents a part of the CGRP^+^ signal and can be observed throughout the tissue. Scale bars are 200 μm. (D) Genetically labeled Aδ-HTMRs are found in the trigeminal ganglia. Representative confocal images of *Smr2^Cre^; Calca-FlpE; R26^LSL-FSF-tdT^* and *Bmpr1b^Cre^; Calca-FlpE; R26^LSL-FSF-tdT^* trigeminal ganglion sections. As in the DRG, reporter^+^ cells co-express CGRP and do not bind IB4. Scale bars are 200 μm.

**Figure S7.**
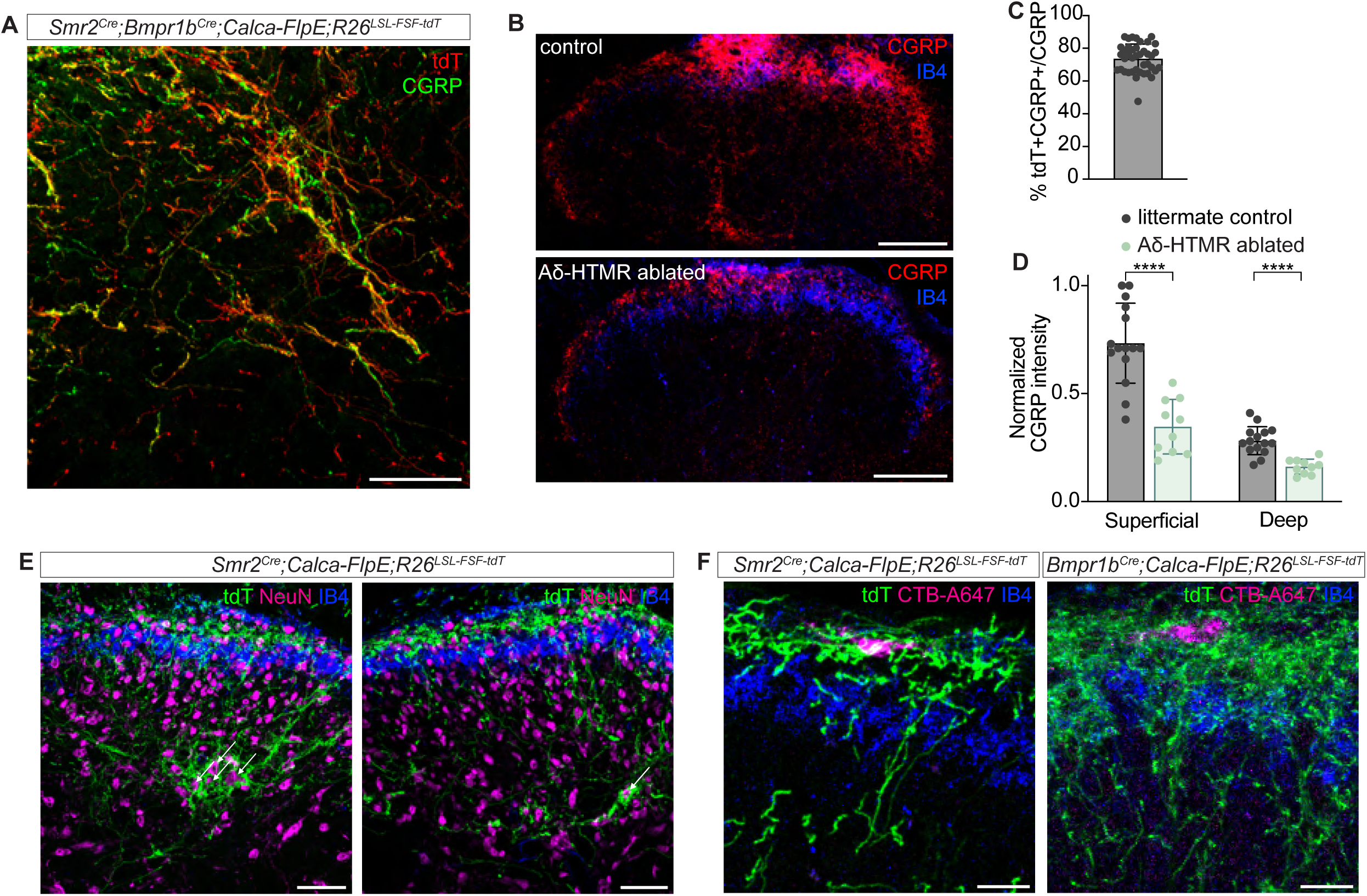
Central termination patterns of genetically labeled Aδ-HTMRs, related to Figure 6. (A) A representative confocal image of the *Smr2^Cre^; Bmpr1b^Cre^; Calca-FlpE; R26^LSL-FSF-tdT^* deep dorsal horn from a lumbar spinal cord section. This section was stained using both anti-tdTomato and anti-CGRP antibodies and shows that most CGRP^+^ axons are tdTomato^+^. Scale bar is 50 μm. (B) Representative confocal images of the control (wild type littermate) and Aδ-HTMR-ablated (*Smr2^Cre^; Bmpr1b^Cre^; Calca^iDTR^*) dorsal horn from lumbar spinal cord sections. Note that the CGRP signal is drastically diminished in the animal subjected to Aδ-HTMR ablation. Scale bar is 100 μm. (C) The majority of the CGRP signal overlaps with the Aδ-HTMR-reporter signal. Quantification related to the example shown in (A). Each dot represents a spinal cord section (n = 30 sections across 3 animals). (D) The CGRP signal is significantly diminished in animals subjected to Aδ-HTMR ablation. Quantification related to the example shown in (B). The Aδ-HTMR-ablated group consists of *Smr2^Cre^; Bmpr1b^Cre^; Calca^iDTR^* animals. Each dot represents a spinal cord section (n = 15 across 3 animals for controls, 10 across 2 animals for Aδ-HTMR-ablated group; *\*\*\*\*p* ≤ 0.0001, unpaired t test). (E) Aδ-HTMRs endings form nests around large cell bodies in the deep dorsal horn of the spinal cord. Representative confocal images from *Smr2^Cre^; Calca-FlpE; R26^LSL-FSF-tdT^* spinal cord sections. Nests of Aδ-HTMR endings are pointed out with white arrows. Scale bars are 50 μm. (F) Aδ-HTMR terminals are in close proximity to the lamina I anterolateral tract (ALT) projection neurons. Representative confocal images from *Smr2^Cre^; Calca-FlpE; R26^LSL-FSF-tdT^* and *Bmpr1b^Cre^; Calca-FlpE; R26^LSL-FSF-tdT^* spinal cord sections. ALT projection neurons were retrogradely labeled with CTB-A647 injection into the PBN. Scale bars are 20 μm.

## Methods

### Key resources table

| REAGENT or RESOURCE | SOURCE | IDENTIFIER |
| --- | --- | --- |
| **Antibodies** |  |  |
| Goat anti-mCherry (1:500, IHC) | OriGene | RRID:AB_2333093 |
| Rabbit anti-dsRed (1:500, IHC) | Takara Bio | RRID:AB_10013483 |
| Chicken anti-RFP (1:500, IHC) | Rockland | RRID:AB_10703148 |
| Goat anti-GFP (1:500, IHC) | US Biological Life Sciences | G8965-01E |
| Chicken anti-NFH (1:500, IHC, skin) | Aves Labs | RRID:AB_2313552 |
| Rabbit anti-NFH (1:500, IHC, DRG) | Sigma-Aldrich | RRID:AB_477272 |
| Rabbit anti-CGRP (1:500, IHC) | Immunostar | RRID:AB_572217 |
| Goat anti-TrkA (1:500, IHC) | R&D Systems | RRID:AB_2283049 |
| Rabbit anti-CalB (1:5000, IHC) | Swant | RRID:AB_3107026 |
| Rabbit anti-TrpV1 (1:500, IHC) | Alomone Labs | RRID:AB_2313819 |
| Rabbit anti-P2X3 (1:500, IHC) | Alomone Labs | RRID:AB_2313760 |
| Rabbit anti-S100 (1:500, IHC) | VWR/ProteinTech | RRID:AB_2254244 |
| Rabbit anti-NeuN (1:500, IHC) | Millipore | RRID:AB_10807945 |
| Goat anti-CD31 | R&D Systems | RRID:AB_2161028 |
| IB4 (Alexa Fluor 488 conjugated) | Thermo Fisher | I21411 |
| IB4 (Alexa Fluor 647 conjugated) | Thermo Fisher | I32450 |
| Donkey anti-goat (Alexa Fluor 546) | Thermo Fisher | RRID:AB_2534103 |
| Donkey anti-goat (Alexa Fluor 647) | Thermo Fisher | RRID:AB_11192020 |
| Donkey anti-goat (Alexa Fluor 488) | Thermo Fisher | RRID:AB_2534102 |
| Donkey anti-rabbit (Alexa Fluor 546) | Thermo Fisher | RRID:AB_2534016 |
| Donkey anti-rabbit (Alexa Fluor 647) | Thermo Fisher | RRID:AB_2536183 |
| Donkey anti-rabbit (Alexa Fluor 488) | Thermo Fisher | RRID:AB_2535792 |
| Donkey anti-chicken (Alexa Fluor 555) | Thermo Fisher | RRID:AB_2921071 |
| Donkey anti-chicken (Alexa Fluor 488) | Thermo Fisher | RRID:AB_2921070 |
| Donkey anti-chicken (Alexa Fluor 647) | Jackson | RRID:AB_2340380 |
| **Bacterial and virus strains** |  |  |
| AAV9-CAG-FLEX-hPLAP | BCH viral core | N/A |
| AAV2/9-CAG-CreON-FlpON-SEAP | BCH viral core | N/A |
| AAV2-retro-hSyn-FlpO | BCH viral core | N/A |
| rAAV1/2-Cre virus | Penn Viral Core | N/A |
| **Chemicals, peptides, and recombinant proteins** |  |  |
| Urethane | Sigma | U2500 |
| Isoflurane | Covetrus | 029405 |
| Tamoxifen | Sigma | T5648-1g |
| Paraformaldehyde (PFA), reagent grade, crystalline | Sigma | P6148-500G |
| Picric acid-formaldehyde (PAF) fixative (Zamboni) | Fisher | NC9335034 |
| Normal Donkey Serum | Jackson | 017-000-121 |
| Benzyl alcohol | Sigma | 108006-500 ml |
| Benzyl benzoate 99% | Fisher | AC105860010 |
| OCT | Fisher | 1437365 |
| Fluoromount-G | Southern Biotech | 0100-01 |
| DAPI-Fluoromount-G | Southern Biotech | 0100-20 |
| CTB-Alexa Fluor(R) 555 conjugate | Fisher | C34776 |
| CTB-Alexa Fluor(R) 647 conjugate | Fisher | C34778 |
| BCIP | Sigma | 11383221001 |
| NBT | Sigma | 11383213001 |
| **Critical commercial assays** |  |  |
| RNAscope™ Multiplex Fluorescent Reagent Kit v2 | ACD bio | 323100 |
| **Experimental models: Organisms/strains** |  |  |
| Mouse: *Smr2*<sup>Cre</sup> | Jax, Qi et al.<sup>23</sup> | RRID:IMSR_JAX:039562 |
| Mouse: *Bmpr1b*<sup>Cre</sup> | Jax, Sharma et al.<sup>28</sup> | RRID:IMSR_JAX:039561 |
| Mouse: *Sstr2*<sup>CreER</sup> | Jax, Qi et al.<sup>23</sup> | RRID:IMSR_JAX:039563 |
| Mouse: *MrgprD*<sup>CreER</sup> | Jax, Olson et al.<sup>31</sup> | RRID:IMSR_JAX:031286 |
| Mouse: *Cysltr2*<sup>Cre</sup> | Jax, Qi et al.<sup>23</sup> | RRID:IMSR_JAX:039985 |
| Mouse: *Trpm8*<sup>FlpO</sup> | Jax, Qi et al.<sup>23</sup> | RRID:IMSR_JAX:039564 |
| Mouse: *TrkB*<sup>CreER</sup> | Jax, Rutlin et al.<sup>32</sup> | RRID:IMSR_JAX:027214 |
| Mouse: *R26*<sup>LSL-FSF-tdTomato</sup> | Jax, Madisen et al.<sup>85</sup> | RRID:IMSR_JAX:021875 |
| Mouse: *R26*<sup>LSL-FSF-ReaChR::mCitrine</sup> | Jax, Madisen et al.<sup>86</sup> | RRID:IMSR_JAX:024846 |
| Mouse: *R26*<sup>LSL-ReaChR::mCitrine</sup> | Jax | RRID:IMSR_JAX:026294 |
| Mouse: *R26*<sup>LSL-tdTomato</sup> | Jax, Madisen et al.<sup>86</sup> | RRID:IMSR_JAX:007914 |
| Mouse: *TIGRE*<sup>LSL-GCaMP6f</sup> | Jax, Daigle et al.<sup>87</sup> | RRID:IMSR_JAX:030328 |
| Mouse: *TIGRE*<sup>LSL-FSF-GCaMP7s</sup> | Jax | RRID:IMSR_JAX:034112 |
| Mouse: *Tau*<sup>FSFiAP</sup> | Jax, Lehnert et al.<sup>88</sup> | RRID:IMSR_JAX:039981 |
| Mouse: *MrgprB4*<sup>Cre</sup> | Jax, Vrontou et al.<sup>33</sup> | RRID:IMSR_JAX:021077 |
| Mouse: *MrgprA3*<sup>Cre</sup> | Han et al.<sup>34</sup> | N/A |
| Mouse: *Calca-FlpE* | Choi et al.<sup>70</sup> | N/A |
| Mouse: *Avil*<sup>FlpO</sup> | Choi et al.<sup>70</sup> | N/A |
| Mouse: *Calca*<sup>iDTR</sup> | McCoy et al.<sup>89</sup> | N/A |
| Mouse: *Ptgfr*<sup>CreER</sup> | This study | N/A |
| **Oligonucleotides** |  |  |
| mm-Bmpr1b-C3 | ACD bio | 533941-C3 |
| mm-Smr2-C2 | ACD bio | 538931-C2 |
| mm-Calca | ACD bio | 420361 |
| mm-Piezo2-C3 | ACD bio | 1228811-C3 |
| mm-Ptgfr-C1 | ACD bio | 501841 |
| mm-Ntrk3-C3 | ACD bio | 423621-C3 |
| mm-Scn10a-C2 | ACD bio | 426011-C2 |
| mm-Prdm12-C3 | ACD bio | 524371-C3 |
| mm-Kit-C2 | ACD bio | 314151-C2 |
| mm-Chrna7-C3 | ACD bio | 465161-C3 |
| GCaMP | ACD bio | 557091 |
| **Software and algorithms** |  |  |
| MATLAB | Mathworks | RRID:SCR_001622 |
| Python | The Python Software Foundation | RRID:SCR_008394 |
| FIJI | NIH | RRID:SCR_002285 |
| Arduino IDE | Arduino | RRID:SCR_024884 |
| GraphPad Prism | GraphPad | RRID:SCR_002798 |
| ZEISS ZEN Microscopy Software | Zeiss | RRID:SCR_013672 |

### Mice

All mice used in this study were maintained on a mixed background. Adult mice (1.5-20 months old) were used, unless specified otherwise. Both male and female mice were used for all experiments. Mice were handled and housed in accordance with Harvard Medical School and IACUC guidelines. Mice were kept in a temperature- and humidity-controlled room with a 12-hour light/dark cycle, with food and water available *ad libitum*.

*Smr2^Cre^* (strain #039562, ^23^), *Bmpr1b^Cre^* (strain #039561, ^28^), *Sstr2^CreER^* (strain #039563, ^23^), *MrgprD^CreER^* (strain #031286, ^31^), *MrgprB4^Cre^*(strain #021077, ^33^), *Cysltr2^Cre^* (strain #039985, ^23^), *Trpm8^FlpO^* (strain #039564, ^23^), *TrkB^CreER^* (strain #027214, ^32^), *R26^LSL-FSF-tdTomato^* (strain #021875, ^85^), *R26^LSL-FSF-ReaChR::mCitrine^* (strain #024846), *R26^LSL-ReaChR::mCitrine^* (strain #026294), *R26^LSL-tdTomato^* (strain #007914, ^86^), *TIGRE^LSL-GCaMP6f^* (strain #030328, ^87^), *TIGRE^LSL-FSF-GCaMP7s^* (strain #034112), *Tau^FSFiAP^* (strain #039981, ^88^) mice are available at Jackson Laboratory. The following mouse lines have been previously described: *MrgprA3^Cre^* ^34^, *Calca-FlpE* ^70^, *Avil^FlpO^* ^70^, *Calca^iDTR^* ^89^. The *Ptgfr^CreER^* mouse line was generated at the Janelia Research Campus Gene Targeting and Transgenic Facility using CRISPR-based homologous recombination techniques in embryonic stem (ES) cells by inserting a CreER cassette into the first coding exon of the *Ptgfr* gene. Chimeras were produced via blastocyst injection, and germline transmission was confirmed through standard tissue genotyping PCR. The *Ptgfr^CreER^* mice can be bred as heterozygotes or homozygotes and express no overt phenotype. The *R26^FSF-ReaChR^* line was derived from the *R26^LSL-FSF-ReaChR::mCitrine^* line by germline excision of LSL cassette.

### Tamoxifen treatment

Tamoxifen (Sigma, T5648) was dissolved in sunflower oil and stored at -80°C until used. Tamoxifen was delivered intraperitoneally (i.p.). *Sstr2^CreER^* animals were treated with 2 mg at P21; *MrgprD^CreER^* animals were treated with 2 mg at P21; *Ptgfr^CreER^* animals were treated with 1 mg at P21; *TrkB^CreER^* animals were treated with 0.5 mg at P5. At least 3 weeks were allowed for Cre induction before using mice for experiments.

### DTX treatment

Diphtheria toxin (Sigma, D0564) (DTX) was dissolved in saline and stored at -20°C until used. DTX was administered i.p. over 10 days at a dose of 20 ng/g of mouse weight per day. Ablation efficiency was quantified with *in situ* hybridization (RNAScope™).

### Optical activation behavior

#### Setup

Mice were habituated to the room and holding chambers for 2 days before the experiment. On the day of the testing, mice were placed on top of a wire rack in holding chambers for 10 minutes to acclimate. 470 mN LED (Thorlabs, M470F3) was used with Ø1000 µm, 0.39 NA optic fiber (Thorlabs, M35L01) and 532 nm, f=7.86 mm, NA=0.51 collimator (Thorlabs, F240SMA-532) for optical stimulation. LED power was measured to be ∼130 mW/cm^2^. 5-ms light pulses were triggered using a custom MATLAB and Arduino program and delivered from below to the plantar surface of animal hindpaws. Each animal received 12 stimuli to alternating hindpaws, each stimulus 5 seconds apart. Mouse behavior was recorded with a high-speed camera (Edmund Optics, Cat. #11-506) at 200 fps.

#### Analysis

Videos of mouse behavior were viewed and analyzed manually. Stimuli were observed on the video, and the latencies to select behaviors were calculated by counting frames elapsed. The first 10 stimulus presentations were considered to calculate behavior frequency. The assay and quantification were performed blind to genotypes.

### Real-time place preference

#### Setup

One week prior to testing, animals were moved to the testing room and habituated to investigator handling by undergoing tail inking. Animals were group habituated to the black matte test chamber (12 in L x 6 in W x 7 in H), center divided with an arched doorway and allowed to freely explore for 5 minutes. The test chamber was mounted on an optically clear acrylic floor with an opaque 1-in border along the chamber walls on each side of the chamber to create a safety zone for escape from painful laser stimulation. A laser and three LED lights (Thorlabs, M470L5 #M01048055; LEDD1B driver #M01063661) were mounted below the floor, oriented to illuminate up through the floor of the stimulation side of the chamber. The LEDs were mounted 1/2 inch below the floor surface to create a 3-in halo of light stimulation overlapping each LED to fill the entire floor with light. To encourage travel to the light stimulation side, the illumination power was graded from 7-10 mW/cm^2^ closest to the door opening to 10-12.5 mW/cm^2^ in the center of the floor to 14-15.5 mW/cm^2^ along the far wall. During the room/chamber habituation, animals were also habituated to the LED light source at very low (∼0 mW) power. The room and test platform were lined with odorless 1-in thick soundproof panels (Bonded Logic Natural Fiber Acoustic Sound Absorbing panels, Model #60600-11212).

The test lasted 3 minutes and consisted of 5 stages.

Stage 1: 1 minute - exploration period. Mice were allowed to explore both sides of the chamber with no illumination.

Stage 2: 30 seconds - Light ON. The “laser” side of the chamber was illuminated.

Stage 3: 30 seconds - Light OFF. No illumination.

Stage 4: 30 seconds - Light ON. The “laser” side of the chamber was illuminated.

Stage 5: 30 seconds - Light OFF. No illumination.

Following test completion, the animal was removed from the testing chamber and returned to their home cage. The equipment was cleaned, and the system was reset for the next animal.

A custom Bonsai script was used to trigger the LEDs and lasers and Imaging source USB 2.0 camera. Mouse locomotion was recorded with a top-down camera at 60 fps.

#### Analysis

Animal locomotion was tracked, and time spent on each side was calculated with a custom MATLAB program. The assay and quantification were performed blind to genotypes. The relative preference for the laser side was defined as the difference between fraction of time spent on the laser side during ON periods and fraction of time spent on the laser side during the exploration (laser off) period.

### Pin prick assay

#### Setup

Mice were habituated to the room and holding chambers for 2 days before the experiment. Importantly, when the same animals were tested in any other assay, the pin-prick assay was done last. The same setup was used as for the optical activation behavior, with modifications. A house-made pinprick apparatus was added; upward movement of a sharp pin (FST, 26007-02) was triggered with a manual push-button. An accelerometer (SparkFun, ADXL335) was secured to the wire rack on which the mice were standing to pick up their movements. To ensure that data consist only of trials where the pin made strong contact with the paw skin, a circuit was set up to measure conductance between the pin and the mouse paw that rested on a conductive metal grid. A DAQ board (National Instruments, USB-6002) was used to synchronously record accelerometer signals along with button presses and skin contact signals. These data were collected for 3 minutes as an experimenter delivered multiple pinpricks to the plantar surface of animals’ hindpaws. Animal behavior was also recorded with a camera at 200 fps.

#### Analysis

Custom MATLAB code was used to read and analyze recorded analog data. Each push of a button was considered a putative trial. Data from all trials were visually inspected. The criteria for selecting a trial for further analysis included 1) pin-to-skin contact initiated after the start of the trial, 2) conductance value for pin-to-skin contact value exceeding 1.5 V, which was subjectively deemed to correspond to strong contact when analyzing corresponding videos, and 3) low baseline accelerometer values (movement) before the start of the trial. Stimulus onset was determined to be at the time of pin to skin contact. Movement onset was empirically determined to be at the time when accelerometer value reached 0.15 V. For trials that passed the quality control, the movement magnitude was computed as area under the curve for accelerometer values across 120 ms after stimulus onset. For presentation, accelerometer voltage readings were converted to arbitrary units (AU) such that 1 AU = maximum accelerometer voltage reading across all trials. The assay and quantification were performed blind to genotypes.

### General methods for open field, balance beam, temperature preference, and texture preference assays

The assays were performed as previously described^90^. The assays were run in the following order: open field, balance beam, temperature preference (cold), temperature preference (hot), texture aversion, pin prick. Mice were provided with chew snacks and always habituated to the testing room at least 30 minutes before handling. Animals were briefly group-habituated to the test chambers. The chambers were cleaned with ECOS unscented dish soap and 70% ethanol between animals. All assays and quantifications were performed blind to genotypes.

### Open field assay

The testing chamber was made of white acrylic (10 in L x 8 in W x 8 in H). Construction paper (Pacon® Tru-Ray) was placed on the floor of the chamber. After habituation, the mice were placed in the testing chamber alone for 5 minutes. Their activity was recorded with a top-down camera at 30 fps. A custom MATLAB program was used to track locomotion and calculate time traveled and time spent in the center of the chamber.

### Temperature preference assay

The testing chamber used was made of black acrylic (12 in L x 6 in W x 7 in H) with an arched doorway across the center and rested on 2 metal temperature control plates (TE Technology, TC-720). For habituation, both plates were set to 30°C. For cold temperature testing, one plate was set to 30°C (control) and the other to 18°C (cold). For hot temperature testing, one plate was set to 35°C (control) and the other to 46°C (hot). After habituation, the mice were placed in the testing chamber alone for 5 minutes. Their activity was recorded with a top-down camera at 30 fps. A custom MATLAB program was used to track locomotion and calculate time spent on either side of the chamber.

### Texture aversion assay

The testing chamber used was made of black acrylic (12 in L x 6 in W x 7 in H) with no divider and rested on an acrylic floor. For habituation, construction paper (Tru-Ray Heavyweight, Pacon) was placed on the floor. For testing, half of the floor was covered with construction paper, and the other half was covered with coarse 60-grit sandpaper (ceramic version, 26060PGP-4, 3M). After habituation, the mice were placed in the testing chamber alone for 10 minutes. Their activity was recorded with a top-down camera at 30 fps. A custom MATLAB program was used to track locomotion and calculate time spent on either side of the chamber.

### Balance beam assay

Animals were trained to walk on the beam for 2 days prior to testing. The balance beam was constructed of 1 m matte white acrylic with a flat 12 mm surface. The beam was mounted 50 cm above the bench top. A soft pad was stretched below the beam to serve as a soft-landing surface to cushion falls. A black matte acrylic box was placed at each end of the beam to provide a start and end enclosure for the animal. During the training, the animals were placed on the beam and encouraged to move across it towards an enclosure. On testing day, animals were placed into the enclosure on one end and allowed to cross. Four beam crossings were recorded with top-down and side cameras at 60 fps. Videos were manually analyzed to determine time to cross, number of slips, and number of stalls.

### Reflex assay

#### Transection surgery

Spinal transection was performed as previously described^91^. Mice were anesthetized with inhaled isoflurane and injected with carprofen at a dose of 5 mg per kg of mouse weight. Ophthalmic ointment was used to protect the eyes. The fur below the neck was shaved and removed with Nair. 100 μL of 2% lidocaine (Covertus, 2468) was injected subcutaneously around the incision. A skin incision was made over the thoracic segment of the spinal column, and the vertebral muscles were removed to expose the gap between spinal segments T9 and T10. Spring scissors were then inserted into the gap, and the spinal cord was cut. To ensure a complete cut, a curved blade was then inserted into the gap and moved laterally. Care was taken to avoid cuts to the underlying lungs. The incision was then sutured, and the mouse was allowed to recover for 6 hours before testing. A transection was deemed successful if lower limb paralysis was observed. Mice were euthanized after the testing.

#### Setup

The same general procedure as for the optogenetic activation behavior was used. The mice were tested the day before the spinal transection surgery and 6 hours after the transection. After the transection, the animals’ lower limbs were paralyzed, so the hindpaws were moved down through the wire rack for better stimulus access. Brush, light, and pinch stimuli were applied to the paws of the animals. Animal behavior was recorded with a high-speed camera at 200 fps. Only the animals with reduced responses to brushing and retained responses to pinching were considered for analysis.

#### Analysis

Videos of mouse behavior were analyzed manually, similar to optogenetic activation behavior. Frames were counted from stimulus onset to movement onset (latency) and from movement onset to movement conclusion (duration).

### Skin injections

Mice aged P7-10 were anesthetized with inhaled isoflurane. AAV9-CAG-FLEX-hPLAP (1.19E+13 gc/mL, custom-made by Janelia Viral Core) or AAV2/9-CAG-CreON-FlpON-SEAP (9.18E+13 gc/mL, custom-made by Boston Children’s Hospital Viral Core) virus was injected into the glabrous skin of hindpaws in a volume of 1.5-2 μL with a borosilicate glass pipette at various dilutions (1:100 to 1:150) for sparse labeling. To label glabrous innervating DRG neurons for loose patch electrophysiology, 2 μL of 2 μg/mL CTB-A555 (Fisher, C34776) was injected into the glabrous skin. A small amount of Fast Green dye was mixed into the virus or CTB solution to visualize the injection. Care was taken to avoid blood vessels and potential systemic spread of the virus.

### Perfusion

Mice were anesthetized with a high dose of inhaled isoflurane and transcardially perfused with ∼10 mL of 1X PBS and ∼15 mL of 4% PFA in 1X PBS. Unless specified otherwise, tissues were collected and post-fixed in 4% PFA rotating at 4°C overnight, then washed with 1X PBS 3 times for ∼10 minutes to avoid overfixation.

### Glabrous skin collection

This tissue collection protocol was used for immunohistochemistry but not whole-mount alkaline phosphatase staining. Mice were deeply anesthetized with ketamine (125 μg/g) and xylazine (12.5 μg/g). Glabrous skin was dissected off the paw while the mouse was anesthetized and placed in 1% PFA at 4°C for 2 hours. The skin samples were then washed in 1X PBS and microdissected further as described below. The samples were immediately processed. The mouse was euthanized or perfused, if other tissues were collected.

### Whole-mount alkaline phosphatase (AP) staining

The staining was performed as previously described^88^. 3 weeks after virus injection, mice were perfused and tissues were collected as described. Post-fixed and washed paws and spinal columns were used for these experiments. Glabrous skin was removed from the hindpaw in one piece, and any remaining fat, connective tissue, and hypodermis were cleaned off. Spinal cords were microdissected from spinal columns with DRG attached. All microdissected tissues were moved to fresh 1X PBS and incubated for 2 (skin) or 2.5 (spinal cord) hours at 67°C. Tissues were then washed in AP detection buffer (0.1 M Tris pH 9.5, 0.1 M NaCl, 50 mM MgCl_2_, 0.1% Tween-20) 3 times for 5 minutes at room temperature. Substrate was then provided (3.4 μL of NBT and BCIP in AP detection buffer), and the tissues were incubated in the dark with slight agitation for approximately 24 hours. To stop the reaction, tissues were then pinned flat on a sylgard plate to prevent curling and incubated in 4% PFA at room temperature for 1 hour. Finally, tissues were dehydrated with serial incubation in 50%, 70%, and 100% ethanol at room temperature for 1 hour each and stored in 100% ethanol until imaging at 4°C. Spinal cords were incubated in 100% methanol for a few hours prior to imaging. When ready, tissues were incubated in BABB (1 part Benzyl Alcohol, 2 parts Benzyl Benzoate) until clear and imaged in BABB under Zeiss AxioZoom stereoscope in brightfield mode.

### Single neuron morphological quantifications

Images were acquired as described above of single neurons in the skin and spinal cord for which a clear single axon could be traced out of the tissue. Images were viewed and analyzed in ImageJ with the SNT plugin. Individual peripheral and central arbors were reconstructed using SNT. For peripheral arbors, branchpoints were manually counted, and area was calculated by drawing a tight polygon surrounding the arbor. For central arbors, collaterals were manually counted, and the number of spinal segments was approximated based on the number of spinal roots.

### Overlap index quantification

Images from sparsely labeled (as described above) and densely labeled (from *Smr2^Cre^; Calca-Flp; Tau^FSFiAP^* and *Bmpr1b^Cre^; Calca-Flp; Tau^FSFiAP^* mice) skin were compared for this analysis. A square of defined area was drawn around similar hindpaw regions on two images. Then, all axons in the field of view were reconstructed, the reconstructed images were binarized, and black and white pixels were counted. Axon density was defined as the number of black pixels divided by the total number of pixels in the field of view. Axon density in the sparsely labeled image was divided by the axon density in the densely labeled image to get an overlap index value.

### Cryosectioning

In preparation for cryosectioning, microdissected tissues were placed in 30% sucrose at 4°C for 1-3 nights or until the tissue sank. Tissues were then placed in cryomolds in Optimal Cutting Temperature (OCT; Andwin Scientific Tissue-Tek™ CRYO-OCT Compound) compound and frozen over dry ice. Samples were sectioned at 20 μm (DRG) or 30 μm (all other tissue types) directly onto Superfrost Plus slides and dried at room temperature in the dark overnight.

Teeth within the mandibles were decalcified prior to the sucrose cryoprotection step in Morse’s solution (22.5% formic acid, 10% tri-sodium citrate in Milli-Q water) for 30-40 hours with rotation at room temperature as previously described^92^.

### Immunohistochemistry

All imaging was performed with an LSM-900 confocal microscope, unless otherwise specified.

#### Staining skin, kidney, and teeth samples (cryosections)

ImmEdge Pen was used to draw a border around the tissue on the slide. Slides were rehydrated for 5 minutes with 1X PBS and then washed 2 times for 5 minutes with 0.1% TritonX-100 in 1X PBS. The slides were then incubated in blocking solution (5% normal donkey serum, 0.1% TritonX-100 in 1X PBS) for 2 hours at room temperature. Primary antibodies were added in blocking solution, and slides were incubated overnight at 4°C. After washing with 0.1% TritonX-100 in 1X PBS, secondary antibodies were added in blocking solution, and the slides were incubated for 2 hours at room temperature. The slides were then washed again and mounted with DAPI Fluoromount (SouthernBiotech, 0100-20).

#### Staining DRG and spinal cord samples (cryosections)

ImmEdge Pen was used to draw a border around the tissue on the slide. Slides were washed with 1X PBS 3 times for 5 minutes and then incubated in blocking solution (5% normal donkey serum, 0.1% TritonX-100 in 1X PBS) for 2 hours at room temperature. Primary antibodies were then added in blocking solution and slides were incubated at 4°C overnight. Slides were washed 4 times for 5 minutes with 0.02% Tween-20 in 1X PBS. Secondary antibodies were added in blocking solution, and the slides were incubated for 2 hours at room temperature. Slides were washed again and mounted with DAPI Fluoromount (SouthernBiotech, 0100-20).

#### Staining sagittal spinal cord sections (free-floating)

The staining was performed in a 12-well plate. 60 µm cryosections were collected directly into the wells containing 1X PBS. All steps were performed with gentle agitation. Sections were incubated in blocking solution (5% normal donkey serum, 0.3% TritonX-1200 in 1X PBS) for 1 hour at room temperature and then transferred to blocking solution containing primary antibodies overnight at 4°C. After 3 washes with 1X PBS, sections were incubated in blocking solution containing secondary antibodies for 2 hours at room temperature. After 3 more washes, sections were mounted on a slide with DAPI Fluoromount (SouthernBiotech, 0100-20) using a paintbrush.

#### Staining kidney and bladder samples (whole-mount)

Post-fixed and washed tissues were used. Kidneys were bisected lengthwise to improve antibody penetration. Tissues were washed with 1% TritonX-100 in 1X PBS for 6-8 hours at room temperature, changing solution every 30 minutes. Tissues were then transferred into blocking solution (5% normal donkey serum, 20% DMSO, 1% TritonX-100 in 1X PBS) with primary antibodies and incubated for 3 nights with rotation at room temperature protected from light. After another 6-8 hours of washes, tissues were incubated in blocking solution with secondary antibodies for 3 nights with rotation at room temperature protected from light. The tissues were then washed for 6-8 hours again, dehydrated with serial incubation in 50%, 70%, and 100% methanol at room temperature for 1 hour each, and stored in 100% methanol. The tissues were then cleared by incubating in BABB. An imaging well was made on a tissue slide with vacuum grease, and a coverslip was placed on top. The bladder was cut to flatten.

#### Staining cranial meninges (whole-mount)

Skullcaps were post-fixed and washed as described. Meninges were carefully peeled off the skull. The meninges were incubated in blocking solution (20% normal donkey serum, 0.3% TritonX-100 in 1X PBS) for 1 hour at room temperature. Primary antibodies were then added to the blocking solution, and the tissue was incubated overnight at 4°C. After the tissue was washed with 0.3% TritonX-100 in 1X PBS 3 times for 1 hour at room temperature, secondary antibodies were added to the blocking solution, and the tissue was incubated overnight at 4°C. The tissues were washed again, then rinsed with 1X PBS 2 times and carefully mounted on slides with Fluoromount Gold. A paintbrush was used to flatten the meninges.

### Joint whole-mount immunolabeling and lightsheet microscopy imaging

Knee joints collected from male and female naïve mice aged 10 weeks (n = 2 knees/reporter line) were fixed and decalcified in Morse’s solution for 3 days. Decalcified knee joints were then processed for immunolabeling and tissue clearing as previously described^92,93^. In brief, decalcified knee joints were dehydrated and rehydrated with methanol gradient and decolorized with hydrogen peroxide. The samples were then immunolabeled with RFP antibody (Rockland, 600-401-379, 1:1000) for 2 weeks, followed by a secondary antibody conjugated with AF647 fluorophore (Thermo Fisher Scientific, A21245, 1:1000) to boost the RFP signal for 2 weeks. Immunolabeled samples were embedded in 1% agarose with the patellar tendon facing upwards. After agarose embedding, samples were subsequently dehydrated in methanol gradient, delipidated in dichloromethane, and cleared in dibenzyl ether. Cleared knee joints were imaged by ZEISS Lightsheet 7 (ZEISS) to visualize RFP positive nerves in 3D (Zen Blue 3.7). An air gap between the refractive index matching solution and objective was corrected by setting the objective correction collar at 1.56.

### Human skin immunohistochemistry

#### Biopsy

IRB-approved study was conducted at the Massachusetts General Hospital (Boston) between March 2021 and April 2021 to obtain skin biopsies from 4 healthy, adult subjects. Written, informed consent was obtained before participation.

Skin biopsies (3 mm) were obtained from distal palmar surface of 4^th^ or 5^th^ digits, processed, and analyzed according to consensus standards^94^. Tissue was placed in Zamboni’s fixative (Newcomer Supply) for 3 hours at room temperature, then washed in 1X PBS 3 times.

#### Immunohistochemistry

Glabrous skin samples were cryoprotected in 15% sucrose in 1X PBS overnight at 4°C, then transferred to 30% sucrose in 1X PBS overnight at 4°C. Tissue was embedded in OCT (1437365, Fisher), frozen using dry ice, and immediately cryosectioned (20 µm) and collected onto glass slides. Sections were allowed to dry overnight at 4°C. Slides not used for staining immediately were placed at -20°C. To reduce background, sections were first rehydrated in 1X PBS, washed once in 0.1% Triton X-100 in 1X PBS for 15 minutes, followed by 3 5-minute washes with 1X PBS, then incubated in Image-iT™ FX Signal Enhancer (ThermoFisher, I36933) for 45 minutes at room temperature. Sections were then washed 3 times for 5 minutes in 1X PBS and incubated in blocking solution (5% normal donkey serum, 0.1% TritonX-100 in 1X PBS) for 2 hours at room temperature. Slides were incubated with primary antibodies diluted in blocking solutions at 4°C overnight, washed 3 times for 20 minutes each with 0.1% TritonX-100 in 1X PBS, and then once for 5 minutes with 1X PBS. A second short block was performed for 30 minutes before being incubated with secondary antibodies diluted in blocking solutions for 1.5 hours at room temperature. Slides were then washed again 5 times for 10 minutes each with 0.1% TritonX-100 in 1X PBS, and then once for 5 minutes with 1X PBS. Finally, slides were mounted in DAPI Fluoromount (SouthernBiotech, 0100-20).

### DRG image quantification

Images were analyzed in ImageJ. Maximum intensity projections were obtained, and cells with clearly visible DAPI nuclei were analyzed. Ellipses were drawn around cells of interest, and area was measured; cell diameter was calculated from the area. The diameters for each cell type were then sorted, and values below the 20^th^ percentile were discarded to minimize the chance of including cell fragments. For Supplementary Figure 3P, large diameter was defined as >25 µm.

### Spinal cord image quantifications

For reporter and CGRP overlap quantifications, signals in the two channels were independently thresholded using the triangle method. Area occupied by pixels with above-threshold values was calculated and used for the overlap calculation.

For calculating the CGRP intensity in control and HTMR-ablated animals, sections from control and “ablated” animals were collected onto the same slides to minimize experimental variation. All sections were imaged using the same settings. Images were rotated such that most of the IB4 signal was horizontal. Superficial dorsal horn was defined to be above the IB4 band; deep dorsal horn was defined to be up 250 μm below the center of the IB4 band. Rectangles were drawn around the superficial and deep dorsal horn, and mean gray value in the CGRP channel was measured. Mean gray values were normalized to the highest mean gray value obtained from images on the same slide.

### *In situ* hybridization

Mice were deeply anesthetized with inhaled isoflurane, decapitated, and their spinal columns were extracted and placed on ice. Fine DRG microdissection was performed in dissection medium (DMEM:F12 (1:1) supplemented with 1% pen/strep and 12.5mM D-Glucose) atop ice. DRG were immediately placed in OCT compound in cryomolds and frozen in dry ice-cooled 2-methylbutane. Samples were cryosectioned at 20 μm thickness and stored at -80°C until further use. *In situ* hybridization was done using the ACDBio RNAScope™ platform (RNAscope™ H202 and Protease Reagents, RNAscope™ Multiplex Fluorescent Detection Kit v2, RNAscope™ probes, and Akoya Opal dyes). The following probes were used: *Bmpr1b* (Cat. # 533941), *Smr2* (Cat. # 538931), *Calca* (Cat. # 420361), *Piezo2* (Cat. # 1228811), *GCaMP* (Cat. # 557091), *Ptgfr* (Cat. # 501841), *Scn10a* (Cat. # 426011), *Prdm12* (Cat. # 524371), *Ntrk3* (Cat. # 423621), *Kit* (Cat. # 314151), *Chrna7* (Cat. # 465161). When combined with immunohistochemistry, same propocol as above (*Staining DRG and spinal cord samples (cryosections)*) was used, started after the last HRP block step of the RNAScope™ protocol.

For quantification, cells expressing one or several transcripts of interest were counted in ImageJ. Only cells with a visible DAPI nucleus were counted. Quantifications were done blind to genotypes.

### Retrograde labeling of spinal cord projection neurons

Mice were anesthetized with inhaled isoflurane and placed in a small animal stereotaxic frame (David Kopf Instruments). Sustained-release buprenorphine (0.1 μg/g) was injected subcutaneously before starting the surgery. Ophthalmic ointment was applied to the eyes, and body temperature was maintained around 37°C with a heating pad. A freshly thawed aliquot of 10 μg/mL CTB-A647 (Fisher, C34778) was used for injections. A small amount of Fast Green dye was mixed into the CTB to visualize the injection. A borosilicate glass pipette and a Microinject system (World Precision Instruments) were used for the injections. Mice were used for experiments 3 days after the injections.

#### DCN injections

DCN were exposed as previously described^88^. In a stereotaxic frame, the animal’s neck was bent 45°. Paraspinal muscles were retracted to expose the gap between the base of the skull and the C1 vertebra. The dura was nicked with a bent insulin syringe. A pipette was lowered 150-200 μm below the brain surface near the obex. 75 nL of CTB was injected in 2 sites bilaterally (4 injections total) at a rate of 50 nL/min.

For labeling DCN-projecting sensory neurons, rAAV1/2-Cre virus (Penn Viral Core, 2E+12 gc/mL) or AAV2-retro-hSyn-FlpO virus (Boston Children’s Hospital Viral Core, 2.06E+14 gc/mL) was injected into the DCN of P11 mice.

#### PBN injections

Injection was performed as previously described^70^. Skull surface was leveled, burr holes were drilled with a dental drill, and 150 nL of CTB was injected in 2 sites bilaterally (4 injections total) at a rate of 75 nL/min. For more reliable anterior-posterior axis PBN targeting, a correction factor F was used (F = (bregma - lambda)/4.21). The following coordinates were used: x = ±1.3-1.4, y = 5.0-5.2*F, z = 2.85-3.10 from the brain surface.

### Calcium imaging

#### Setup

The imaging and stimulus delivery were performed as previously described^23^. Mice were anesthetized with inhaled isoflurane, and their body temperature was maintained at 37°C throughout the experiment. The back hair was shaved off, and Nair was applied to remove the remaining hair. An incision was made over the lumbar spine and paravertebral muscles overlaying vertebrae L3-L5 were loosened and pulled away from the bone. The spine was stabilized at the L3-L4 vertebrae with a custom-made clamp. The bone covering L4 DRG was removed with Friedman-Pearson Rongeurs (FST, 16221-14). Bleeding was controlled with Surgifoam (Mckesson, Cat. # 1972) and cotton swabs. The right thigh hair was shaved. The right hindpaw was secured on the imaging platform, glabrous side up, with silicone putty.

The surgical preparation was then transferred to the platform under an upright epifluorescence microscope (Zeiss Axio Examiner) with 10X air objective (Zeiss Epiplan, NA = 0.20). 470 nm LED (Thorlabs, M470L5) with LED driver (Thorlabs, LEDD1B) was used as the light source. A CMOS Camera (Thorlabs, CS505MU1) was triggered at 10 fps with a 50 ms exposure time. All the recorded stimuli were synchronized with the camera and LED using a DAQ board (National Instruments, NI USB-6343).

#### Stimulation

Stimuli were applied to glabrous skin in the following order: airpuff (Dust-Off, DPSXL4A), brush (small paintbrush), von Frey filaments (in order of increasing force; North Coast Medical, NC12775), poking with closed forceps (FST, 11050-10), controlled indentation with a mechanical stimulator, pinch, and temperature stimuli. Pinch and temperature stimuli were then applied to the thigh.

For controlled indentation, an indenter (Aurora Scientific, 300C-I) with a custom-made 200 µm diameter tip was used. The paw was first stimulated manually to identify spots that gave rise to robust calcium responses, and an indenter was then placed on top of that spot. Multiple spots were tested. A custom-written MATLAB program was used to design and trigger the stimuli. Baseline was recorded for 2 s, indentation lasted for 4 s, and 6 s of decay time was allowed. The interstimulus interval was 10 seconds. Each force was presented 3 times.

For thermal stimuli, a Peltier device (13*12*2.5mm, TE-65–0.6–0.8, TE technology) was controlled by a Temperature Controller (TEC1089/PT1000, Meerstetter Engineering) with an RTD Platinum (Pt) thermistor (2952-P1K0.161.6W.A.010-ND, Digikey) mounted on the surface of the Peltier. To maximize the conductivity between the Peltier and skin, a small amount of thermal paste (Aeronaut, Thermal Grizzly) was evenly applied on the top of the Peltier surface, which was then gently pressed on the skin until good contact was formed. A thermocouple microprobe (Physitemp, IT-1E) was inserted between the Peltier device and the skin as a separate measurement of the applied temperature. Thirty seconds after contacting skin, temperature stimuli started from innocuous temperatures progressing to noxious temperatures, in the sequence of 35, 25, 20, 40, 15, 45, 10, 50, 5, 55°C. Stimulus structure was as follows:

1. Baseline (32°C) - 20 seconds
2. Target temperature - 20 seconds
3. Baseline - 50 seconds
4. Back to 1.

The change rate for all temperatures was kept at 5°C s^−1^.

### Calcium imaging analysis

The analysis was performed as previously described^23^. Motion correction and spatial high-pass filtering were conducted using a custom-written macro code that employed the ImageJ plugin “moco” and “Unsharp mask” filter, then regions of interest (ROIs) were manually selected in ImageJ. Intensity measurements were then analyzed in MATLAB with a custom-written code. In the calculation of ΔF/F, F was defined using baseline activity (average intensity before each stimulation). When a cell responded to controlled indentation of multiple spots, responses with the lowest force threshold were selected. Responses to three trials of force presentation were averaged, and the average trace was used for threshold and intensity calculations. Threshold was determined to be the lowest force that evoked a discernible calcium response. Response intensity was calculated as ΔF/F area under the curve (AUC) for 3 seconds of indentation. For the calculation of the percentage of cells responding to a given stimulus for Figure 2C, the number of cells responding to pinch was used as a denominator.

### *In vivo* loose patch targeted DRG recordings

The recordings were performed as previously described^95^. Mice were treated with dexamethasone (2 mg/kg i.p.) and anesthetized with an i.p. injection of urethane (1 g/kg); anesthesia was maintained with 1-1.5% inhaled isoflurane. The L4 DRG was exposed similarly to the procedure described in the calcium imaging section. The exposed DRG was immersed in external solution containing 140 mM NaCl, 3.1 mM KCl, 0.5 mM KH_2_PO_4_, 6 mM glucose, 1.2 mM CaCl_2_, 1.2 mM MgSO_4_ (pH adjusted to 7.4 with NaOH); the same solution was used to fill glass pipettes with a 20–30 μm tip diameter. Fluorescent cell bodies that were labeled with a genetic reporter and CTB-A555 were targeted for loose-seal cell-attached recordings. Extracellular action potentials were measured using a Multiclamp 700A amplifier (Axon Instruments) operated in the voltage clamp configuration. Electrophysiological data were digitized at 40 kHz with a Digidata 1550a (Molecular Devices), low-pass filtered at 10 kHz (four-pole Bessel filter), and acquired using pClamp (Molecular Devices, Version 10). Stimuli were delivered manually or with a mechanical stimulator, as described in the calcium imaging section. A 250 µm diameter indenter tip was used for controlled indentation. For optogenetic stimulation, the same equipment was used as described in the optical activation behavior section. Electrical stimuli were applied using a bipolar electrode to glabrous skin moistened with 0.9% saline. Conduction velocity was measured by electrically stimulating the skin and dividing the distance between stimulation site and cell body by the spike latency. Electrophysiological data were analyzed using custom-made Python code as previously described^95^.

### *In vivo* DRG recordings with sharp glass electrodes

The recordings were performed as previously described^96^. Mice were anesthetized with urethane (1.5 g/kg). L4 DRG was exposed similarly to how described in the calcium imaging section. 1 MΩ glass electrode filled with 0.9% saline was advanced into the right L4 DRG while optical search stimulus was applied to the glabrous hindpaw. Controlled indentations were applied using a custom-built mechanical stimulator as previously described. Electrical stimuli were applied using a bipolar electrode to glabrous skin moistened with 0.9% saline. Conduction velocity was measured by electrically stimulating the skin and dividing the distance between stimulation site and cell body by the spike latency. To test the presence of DCN-projecting collaterals, the DCN were exposed similar to how described in the retrograde labeling section. An optical fibre (400 µm diameter, 0.39 NA) was placed above the DCN, and blue light pulses were delivered to generate antidromic spikes (Thorlabs, M470F3). Recordings were amplified using a Multiclamp 700B commander (Molecular Devices) under the 100X AC differential amplification mode with an additional 20X gain. Data were collected at 20 kHz using a Digidata 1550B (Molecular Devices).

### Spinal cord MEA recordings

The recordings and subsequent analysis were performed as previously described^48^. Briefly, mice were anesthetized with urethane (1.5 g/kg i.p.) and their body temperature was maintained at 37°C throughout the experiment. After exposure, the T13 vertebra was retracted to reveal the dorsal surface of the L4-L5 segments of spinal cord. The dura was removed, and the spinal cord was covered with 0.9% saline. A 32-channel silicon probe (Neuronexus Cambridge Neurotech ASSY-37 H4 optrode) was inserted into the spinal cord and advanced up to ∼700 μm below the dorsal surface under visual guidance. Signals were amplified, filtered (0.1 – 7.5 kHz bandpass), and digitized (20 kHz) using a headstage amplifier and recording controller (Intan Technologies RHD2000 and Recording Controller). Data acquisition was controlled with open-source software (Intan Technologies Recording Controller version 2.07). Spike sorting was performed using JRCLUST.

### Whole cell recordings

#### Retrograde injections

Three days prior to the experiment, 200 nl of 5 mg/μL CTB-A555 was injected bilaterally into the PBN of *Smr2^Cre^; Calca-FlpE; R26^FSF-LSL-ReaChR::mCitrine^* or *Bmpr1b^Cre^; Calca-FlpE; R26^FSF-LSL-ReaChR::mCitrine^* mice aged P14-18. Whole-cell patch-clamp recordings were then performed on CTB^+^ neurons while optically stimulating the ReaChR-containing afferents, as previously described^70^.

#### Slice preparation

Mice (P17-21) were anesthetized with inhaled isoflurane (4% – 5%) while vertebral columns were dissected. Lumbar enlargements were dissected out from vertebral columns in an ice-cold choline solution (92 mM Choline Chloride, 2.5 mM KCl, 1.2 mM NaH_2_PO_4_, 30 mM NaHCO_3_, 20 mM HEPES, 25 mM Glucose, 5 mM Sodium Ascorbate, 2 mM Thiourea, 3 mM Sodium Pyruvate, 10 mM MgSO_4_*7H_2_O, 0.5 mM CaCl_2_*2H_2_O, pH = 7.3-7.4, Osm = 300-310 mOsm) including 1 μM (*R*,*S*)-3-(2-carboxypiperazin-4-yl)propyl-1-phosphonic acid (CPP; Abcam, ab120160) and 1 μM 2,3-dihydroxy-6-nitro-7-sulfamoyl-benzo[f]quinoxaline (NBQX; Abcam, ab120046) and embeded in 5% LMP agarose (Life Technology, 16520-100). The lumbar spinal cords were sliced in a transverse plane (300 μm) (Leica, VT1200S), and the slices were recovered at 34°C for 30 minutes in oxygenated (95% O2 and 5% CO2) holding solution (119 mM NaCl, 2.5 mM KCl, 1.0 mM NaH_2_PO_4_*H2O, 26 mM NaHCO_3_, 25 mM Glucose, 1.3 mM Sodium Ascorbate, 2 mM MgSO_4_*7H_2_O, 2 mM CaCl_2_*2H_2_O, pH = 7.3-7.4, Osm = 300-310 mOsm). After recovery, spinal cord slices were placed at room temperature for 30 minutes prior to recordings.

#### Whole-cell recording procedure

Spinal cord slices were placed in a recording chamber maintained at room temperature and perfused with oxygenated (95% O2 and 5% CO2) recording ACSF (2.5 mM CaCl_2_*2H_2_O, 1.0 mM NaH_2_PO_4_*H2O, 119 mM NaCl, 2.5 mM KCl, 1.3 mM MgSO_4_*7H_2_O, 26 mM NaHCO_3_, 25 mM Glucose, 1.3 mM Sodium L-ascorbate, pH = 7.3-7.4, Osm = 300-310 mOsm). Retrogradely labeled SPB neurons were visualized using a SliceScope Pro 6000 electrophysiology rig (Scientifica) equipped with IR-DIC optics (ThorLabs, M850L3-C4, with COP1-B Collimation Adapter), an LED to fluorescently identify stere^+^ cells (pE-300, Nikon CoolLED), a water-immersion objective (Olympus LUMPlanFLN 40X), and a camera (Teledyne QImaging, Rolera Bolt™ CMOS camera). For the cells within deep laminae, fluorescence from ReaChR::mCitrine^+^ afferents was used as a guide to target neurons in nests, i.e. surrounded by fluorescent axons. The pipette resistance ranged from 3 to 5 MΩ, and the electrodes were filled with an intracellular solution containing biocytin (130 mM Cs-gluconate, 10 mM HEPES, 1 mM EGTA, 0.1 mM CaCl_2_, 5 mM TEA, 1 mM QX-314, 0.2 D600, 0.4% w/v biocytin, pH 7.3, 295 mOsm). To avoid biocytin spillover, pipettes were front-filled with intracellular solution not containing biocytin. Whole-cell capacitance was left uncompensated (∼5 pF) and the series resistance was constantly monitored and compensated online (60-80%). Signals were acquired using a Multiclamp 700A amplifier (Molecular Devices). The data were low-pass filtered at 2 kHz and digitized at 20 kHz (Molecular Devices, Digidata 1440A). For ReaChR-assisted circuit mapping, primary afferent synaptic terminals were stimulated with wide-field white LED illumination with a TxRd filter through the 40x objective (pE-300, Nikon CoolLED, 5 ms pulse width, light intensity = 12.35 mW). Current amplitude, latency, and jitter were analyzed using Clampfit (version 10, Molecular Devices). Cells were recorded first without additional drugs, and then in a solution containing 1 μM tetrodotoxin (TTX) (Tocris, 1069), followed by the addition of 500 μM 4-aminopyridine (4-AP) (Tocris, 940) to the bath.

#### Cell fills

Recordings were maintained for 40 minutes to allow biocytin to perfuse the cell. Subsequently, the slices were fixed in 4% PFA at 4°C overnight and processed for immunohistochemistry. Briefly, slices were washed in 0.3% TritonX-100 in 1X PBS 5 times for 30 minutes, then transferred to blocking solution (5% normal donkey serum, 0.3% TritonX-100 in 1X PBS) for 2 hours at room temperature. Slices were then incubated in blocking solution with primary antibodies for 72 hours, washed as described above and incubated in blocking solution with secondary antibodies for 48 hours. Slices were washed again and then dehydrated with serial incubation in 50%, 75%, and 100% methanol in PBS at room temperature for 10 minutes each and kept in 100% methanol overnight for full dehydration. Before imaging, slices were cleared in BABB solution for 2 hours.

### Quantification and statistical analysis

Statistical analyses were performed in GraphPad Prism 10.2.3 and MATLAB. Data normality was tested with the Shapiro-Wilk test. Comparison between two independent groups was done with unpaired t-test (normally distributed data) and Mann-Whitney test (non-parametric data). Comparison between more than two independent groups was done with one-way ANOVA with Tukey’s multiple comparisons post-hoc test. A permutation test on the mean difference was used to compare calcium response intensity distributions. A p-value ≤ 0.05 was considered statistically significant.

## Supplemental information

### Document S1. Figures S1–S7 and supplemental references

**Video S1. Aδ-HTMRs evoke a fast and robust nocifensive behavior, related to Figure 1**. Behavioral response to optical activation of glabrous skin-innervating Aδ-HTMR in an *Smr2^Cre^; Calca-FlpE; R26^LSL-FSF-ReaChR::mCitrine^* mouse. Stimulus is a 5-ms light pulse. Video is acquired at 200 fps and replayed at 5 fps.

**Video S2. Pinprick paradigm, related to Figure 3**. Pin is pushed up into the mouse paw with a button push. Video is acquired at 200 fps and replayed at 5 fps.

**Video S3. Aδ-HTMRs drive fast reflexive paw movement in spinalized animals, related to Figure 7**. Behavioral response to optical activation of glabrous skin-innervating Aδ-HTMR in an *Smr2^Cre^; Calca-FlpE; R26^LSL-FSF-ReaChR::mCitrine^* mouse after spinal transection. Mouse hindlimbs are paralyzed by the transection and pulled through the wire rack for easy optical access and unrestrained movement. Stimulus is a 5-ms light pulse. Video is acquired at 200 fps and replayed at 5 fps.

**Video S4. C-heat thermoreceptors drive slow reflexive paw movement in spinalized animals, related to Figure 7**. Behavioral response to optical activation of glabrous skin-innervating C-heat thermoreceptors in an *Sstr2^CreER^; R26^LSL-ReaChR::mCitrine^* mouse after spinal transection. Mouse hindlimbs are paralyzed by the transection and pulled through the wire rack for easy optical access and unrestrained movement. Stimulus is a 5-ms light pulse. Video is acquired at 200 fps and replayed at 5 fps.

**Video S5. C-heat thermoreceptors drive a unique delayed paw movement in spinalized animals, related to Figure 7**. An example of extended behavioral response to optical activation of glabrous skin-innervating C-heat thermoreceptors in an *Sstr2^CreER^; R26^LSL-ReaChR::mCitrine^* mouse after spinal transection. Note that the initial withdrawal-like movement is followed by delayed twisting of the paw in absence of additional stimulation. Stimulus is a 5-ms light pulse. Video is acquired at 200 fps and replayed at 25 fps.

## References

1. Finnerup, N.B. and Jensen, T.S. (2006) Mechanisms of disease: mechanism-based classification of neuropathic pain-a critical analysis. Nat Clin Pract Neurol. 2(2): p. 107–15.

2. Vanelderen, P., Rouwette, T., Kozicz, T., Roubos, E., Van Zundert, J., Heylen, R., and Vissers, K. (2010) The role of brain-derived neurotrophic factor in different animal models of neuropathic pain. European Journal of Pain. 14(5): p. 473.e1–473.e9.

3. Dai, D.-W., Xu, Z., Chen, X., Yuan, L., Zhang, A.-J., Zhang, P.-Q., Lu, Y.-M., and Deng, Z.-F. (2014) Distinct roles of neuregulin in different models of neuropathic pain. Neurological Sciences. 35(4): p. 531–536.

4. Cuevas-Diaz Duran, R., Li, Y., Garza Carbajal, A., You, Y., Dessauer, C.W., Wu, J., and Walters, E.T. (2023) Major Differences in Transcriptional Alterations in Dorsal Root Ganglia Between Spinal Cord Injury and Peripheral Neuropathic Pain Models. J Neurotrauma. 40(9-10): p. 883–900.

5. Bangash, M.A., Alles, S.R.A., Santana-Varela, S., Millet, Q., Sikandar, S., De Clauser, L., Ter Heegde, F., Habib, A.M., Pereira, V., Sexton, J.E., Emery, E.C., Li, S., Luiz, A.P., Erdos, J., Gossage, S.J., Zhao, J., Cox, J.J., and Wood, J.N. (2018) Distinct transcriptional responses of mouse sensory neurons in models of human chronic pain conditions. Wellcome Open Research. 3: p. 78.

6. Renthal, W., Tochitsky, I., Yang, L., Cheng, Y.-C., Li, E., Kawaguchi, R., Geschwind, D.H., and Woolf, C.J. (2020) Transcriptional Reprogramming of Distinct Peripheral Sensory Neuron Subtypes after Axonal Injury. Neuron. 108(1): p. 128–144.e9.

7. Perl, E.R. (2011) Pain mechanisms: A commentary on concepts and issues. Progress in Neurobiology. 94(1): p. 20–38.

8. Peirs, C., Williams, S.-P.G., Zhao, X., Arokiaraj, C.M., Ferreira, D.W., Noh, M.-C., Smith, K.M., Halder, P., Corrigan, K.A., Gedeon, J.Y., Lee, S.J., Gatto, G., Chi, D., Ross, S.E., Goulding, M., and Seal, R.P. (2021) Mechanical Allodynia Circuitry in the Dorsal Horn Is Defined by the Nature of the Injury. Neuron. 109(1): p. 73–90.e7.

9. Woolf, C.J. and Ma, Q. (2007) Nociceptors—Noxious Stimulus Detectors. Neuron. 55(3): p. 353–364.

10. Dubin, A.E. and Patapoutian, A. (2010) Nociceptors: the sensors of the pain pathway. Journal of Clinical Investigation. 120(11): p. 3760–3772.

11. Perl, E.R. (1968) Myelinated afferent fibres innervating the primate skin and their response to noxious stimuli. The Journal of Physiology. 197(3): p. 593–615.

12. Adriaensen, H., Gybels, J., Handwerker, H.O., and Van Hees, J. (1983) Response properties of thin myelinated (A-delta) fibers in human skin nerves. J Neurophysiol. 49(1): p. 111–22.

13. Koltzenburg, M., Stucky, C.L., and Lewin, G.R. (1997) Receptive properties of mouse sensory neurons innervating hairy skin. J Neurophysiol. 78(4): p. 1841–50.

14. Nagi, S.S., Marshall, A.G., Makdani, A., Jarocka, E., Liljencrantz, J., Ridderstrom, M., Shaikh, S., O’Neill, F., Saade, D., Donkervoort, S., Foley, A.R., Minde, J., Trulsson, M., Cole, J., Bonnemann, C.G., Chesler, A.T., Bushnell, M.C., McGlone, F., and Olausson, H. (2019) An ultrafast system for signaling mechanical pain in human skin. Sci Adv. 5(7): p. eaaw1297.

15. Peirs, C. and Seal, R.P. (2016) Neural circuits for pain: Recent advances and current views. Science. 354(6312): p. 578–584.

16. Ma, Q. (2022) A functional subdivision within the somatosensory system and its implications for pain research. Neuron.

17. Boada, M.D. and Woodbury, C.J. (2008) Myelinated Skin Sensory Neurons Project Extensively throughout Adult Mouse Substantia Gelatinosa. The Journal of Neuroscience. 28(9): p. 2006–2014.

18. Burgess, P.R. and Perl, E.R. (1967) Myelinated afferent fibres responding specifically to noxious stimulation of the skin. J Physiol. 190(3): p. 541–62.

19. Yu, H., Nagi, S.S., Usoskin, D., Hu, Y., Kupari, J., Bouchatta, O., Yan, H., Cranfill, S.L., Gautam, M., Su, Y., Lu, Y., Wymer, J., Glanz, M., Albrecht, P., Song, H., Ming, G.-L., Prouty, S., Seykora, J., Wu, H., Ma, M., Marshall, A., Rice, F.L., Li, M., Olausson, H., Ernfors, P., and Luo, W. (2024) Leveraging deep single-soma RNA sequencing to explore the neural basis of human somatosensation. Nature Neuroscience. 27(12): p. 2326–2340.

20. Yang, L., Xu, M., Bhuiyan, S.A., Li, J., Zhao, J., Cohrs, R.J., Susterich, J.T., Signorelli, S., Green, U., Stone, J.R., Levy, D., Lennerz, J.K., and Renthal, W. (2022) Human and mouse trigeminal ganglia cell atlas implicates multiple cell types in migraine. Neuron.

21. Kupari, J. and Ernfors, P. (2023) Molecular taxonomy of nociceptors and pruriceptors. PAIN. 164(6): p. 1245–1257.

22. Djouhri, L. (2016) Adelta-fiber low threshold mechanoreceptors innervating mammalian hairy skin: A review of their receptive, electrophysiological and cytochemical properties in relation to Adelta-fiber high threshold mechanoreceptors. Neurosci Biobehav Rev. 61: p. 225–38.

23. Qi, L., Iskols, M., Shi, D., Reddy, P., Walker, C., Lezgiyeva, K., Voisin, T., Pawlak, M., Kuchroo, V.K., Chiu, I.M., Ginty, D.D., and Sharma, N. (2024) A mouse DRG genetic toolkit reveals morphological and physiological diversity of somatosensory neuron subtypes. Cell. 187(6): p. 1508–1526.e16.

24. Rau, K.K., McIlwrath, S.L., Wang, H., Lawson, J.J., Jankowski, M.P., Zylka, M.J., Anderson, D.J., and Koerber, H.R. (2009) Mrgprd Enhances Excitability in Specific Populations of Cutaneous Murine Polymodal Nociceptors. Journal of Neuroscience. 29(26): p. 8612–8619.

25. Beaudry, H., Daou, I., Ase, A.R., Ribeiro-Da-Silva, A., and Séguéla, P. (2017) Distinct behavioral responses evoked by selective optogenetic stimulation of the major TRPV1+ and MrgD+ subsets of C-fibers. Pain. 158(12): p. 2329–2339.

26. Warwick, C., Cassidy, C., Hachisuka, J., Wright, M.C., Baumbauer, K.M., Adelman, P.C., Lee, K.H., Smith, K.M., Sheahan, T.D., Ross, S.E., and Koerber, H.R. (2021) MrgprdCre lineage neurons mediate optogenetic allodynia through an emergent polysynaptic circuit. Pain. 162(7): p. 2120–2131.

27. Sherrington, C.S., The integrative action of the nervous system. 1906, New Haven, London, Oxford: Yale University Press; Henry Frowde; University Press. xvi, 411 p.

28. Sharma, N., Flaherty, K., Lezgiyeva, K., Wagner, D.E., Klein, A.M., and Ginty, D.D. (2020) The emergence of transcriptional identity in somatosensory neurons. Nature. 577(7790): p. 392–398.

29. Usoskin, D., Furlan, A., Islam, S., Abdo, H., Lönnerberg, P., Lou, D., Hjerling-Leffler, J., Haeggström, J., Kharchenko, O., Kharchenko, P.V., Linnarsson, S., and Ernfors, P. (2015) Unbiased classification of sensory neuron types by large-scale single-cell RNA sequencing. Nature Neuroscience. 18(1): p. 145–153.

30. Von Buchholtz, L.J., Ghitani, N., Lam, R.M., Licholai, J.A., Chesler, A.T., and Ryba, N.J.P. (2020) Decoding Cellular Mechanisms for Mechanosensory Discrimination. Neuron.

31. Olson, W., Abdus-Saboor, I., Cui, L., Burdge, J., Raabe, T., Ma, M., and Luo, W. (2017) Sparse genetic tracing reveals regionally specific functional organization of mammalian nociceptors. eLife. 6: p. e29507.

32. Rutlin, M., Ho, C.-Y., Victoria, Cassidy, C., Bai, L., C, and David (2014) The Cellular and Molecular Basis of Direction Selectivity of Aδ-LTMRs. Cell. 159(7): p. 1640–1651.

33. Vrontou, S., Wong, A.M., Rau, K.K., Koerber, H.R., and Anderson, D.J. (2013) Genetic identification of C fibres that detect massage-like stroking of hairy skin in vivo. Nature. 493(7434): p. 669–673.

34. Han, L., Ma, C., Liu, Q., Weng, H.-J., Cui, Y., Tang, Z., Kim, Y., Nie, H., Qu, L., Patel, K.N., Li, Z., McNeil, B., He, S., Guan, Y., Xiao, B., Lamotte, R.H., and Dong, X. (2013) A subpopulation of nociceptors specifically linked to itch. Nature Neuroscience. 16(2): p. 174–182.

35. Cavanaugh, D.J., Chesler, A.T., Jackson, A.C., Sigal, Y.M., Yamanaka, H., Grant, R., O’Donnell, D., Nicoll, R.A., Shah, N.M., Julius, D., and Basbaum, A.I. (2011) *Trpv1*Reporter Mice Reveal Highly Restricted Brain Distribution and Functional Expression in Arteriolar Smooth Muscle Cells. The Journal of Neuroscience. 31(13): p. 5067–5077.

36. Stirling, L.C., Forlani, G., Baker, M.D., Wood, J.N., Matthews, E.A., Dickenson, A.H., and Nassar, M.A. (2005) Nociceptor-specific gene deletion using heterozygous NaV1.8-Cre recombinase mice. Pain. 113(1-2): p. 27–36.

37. Patil, M.J., Hovhannisyan, A.H., and Akopian, A.N. (2018) Characteristics of sensory neuronal groups in CGRP-cre-ER reporter mice: Comparison to Nav1.8-cre, TRPV1-cre and TRPV1-GFP mouse lines. PLOS ONE. 13(6): p. e0198601.

38. Huang, J., Polgár, E., Solinski, H.J., Mishra, S.K., Tseng, P.-Y., Iwagaki, N., Boyle, K.A., Dickie, A.C., Kriegbaum, M.C., Wildner, H., Zeilhofer, H.U., Watanabe, M., Riddell, J.S., Todd, A.J., and Hoon, M.A. (2018) Circuit dissection of the role of somatostatin in itch and pain. Nature Neuroscience. 21(5): p. 707–716.

39. Luo, W., Enomoto, H., Rice, F.L., Milbrandt, J., and Ginty, D.D. (2009) Molecular Identification of Rapidly Adapting Mechanoreceptors and Their Developmental Dependence on Ret Signaling. Neuron. 64(6): p. 841–856.

40. Liu, Y., Rutlin, M., Huang, S., Barrick, C.A., Wang, F., Jones, K.R., Tessarollo, L., and Ginty, D.D. (2012) Sexually Dimorphic BDNF Signaling Directs Sensory Innervation of the Mammary Gland. Science. 338(6112): p. 1357–1360.

41. Bai, L., Brendan, Liu, J., Nicole, Travis, Pann, Cassidy, C., C, and David (2015) Genetic Identification of an Expansive Mechanoreceptor Sensitive to Skin Stroking. Cell. 163(7): p. 1783–1795.

42. Shields, S.D., Ahn, H.-S., Yang, Y., Han, C., Seal, R.P., Wood, J.N., Waxman, S.G., and Dib-Hajj, S.D. (2012) Nav1.8 expression is not restricted to nociceptors in mouse peripheral nervous system. PAIN®. 153(10): p. 2017–2030.

43. Li, L., Rutlin, M., Abraira, V.E., Cassidy, C., Kus, L., Gong, S., Jankowski, M.P., Luo, W., Heintz, N., Koerber, H.R., Woodbury, C.J., and Ginty, D.D. (2011) The functional organization of cutaneous low-threshold mechanosensory neurons. Cell. 147(7): p. 1615–27.

44. Le Pichon, C.E. and Chesler, A.T. (2014) The functional and anatomical dissection of somatosensory subpopulations using mouse genetics. Frontiers in Neuroanatomy. 8.

45. Arcourt, A., Gorham, L., Dhandapani, R., Prato, V., Taberner, F.J., Wende, H., Gangadharan, V., Birchmeier, C., Heppenstall, P.A., and Lechner, S.G. (2017) Touch Receptor-Derived Sensory Information Alleviates Acute Pain Signaling and Fine-Tunes Nociceptive Reflex Coordination. Neuron. 93(1): p. 179–193.

46. Ghitani, N., Barik, A., Szczot, M., Thompson, J.H., Li, C., Le Pichon, C.E., Krashes, M.J., and Chesler, A.T. (2017) Specialized Mechanosensory Nociceptors Mediating Rapid Responses to Hair Pull. Neuron. 95(4): p. 944–954.e4.

47. Abraira, V. and Ginty, D. (2013) The Sensory Neurons of Touch. Neuron. 79(4): p. 618–639.

48. Chirila, A.M., Rankin, G., Tseng, S.-Y., Emanuel, A.J., Chavez-Martinez, C.L., Zhang, D., Harvey, C.D., and Ginty, D.D. (2022) Mechanoreceptor signal convergence and transformation in the dorsal horn flexibly shape a diversity of outputs to the brain. Cell. 185(24): p. 4541–4559.e23.

49. Chen, C.-Y., Do Nascimento, L.F., De Faria, F.M., Carballo, G., Kaczmarczyk, L., Broman, J., Nagi, S., Olausson, H., Jackson, W.S., Szczot, M., and Larsson, M., A fast nociceptive subsystem mediating rapid reflexive behavior but not affective pain. 2025, openRxiv.

50. Dhaka, A., Murray, A.N., Mathur, J., Earley, T.J., Petrus, M.J., and Patapoutian, A. (2007) TRPM8 Is Required for Cold Sensation in Mice. Neuron. 54(3): p. 371–378.

51. Wolfson, R.L., Abdelaziz, A., Rankin, G., Kushner, S., Qi, L., Mazor, O., Choi, S., Sharma, N., and Ginty, D.D. (2023) DRG afferents that mediate physiologic and pathologic mechanosensation from the distal colon. Cell. 186(16): p. 3368–3385.e18.

52. Neubarth, N.L., Emanuel, A.J., Liu, Y., Springel, M.W., Handler, A., Zhang, Q., Lehnert, B.P., Guo, C., Orefice, L.L., Abdelaziz, A., DeLisle, M.M., Iskols, M., Rhyins, J., Kim, S.J., Cattel, S.J., Regehr, W., Harvey, C.D., Drugowitsch, J., and Ginty, D.D. (2020) Meissner corpuscles and their spatially intermingled afferents underlie gentle touch perception. Science. 368(6497): p. eabb2751.

53. Bhuiyan, S.A., Nagi, S.S., Sankaranarayanan, I., Semizoglou, E., Usoskin, D., Yang, L., Yu, H., Arendt-Tranholm, A., Bertels, Z., Bhatia, P., Bouchatta, O., Boyer, K., Cervantes, A., Chalif, J., Chintalapudi, H., Cicalo, A., Copits, B., Cronin, C., Curatolo, M., Dong, X., Dougherty, P.M., Dourson, A., Funk, G., Gabriel, K., Griesemer, D.S., Guo, H., Gupta, P., Hofstetter, C., Horton, P., Hsieh, A., Inturi, N.N., Jain, A., Jayakar, S., Johnston, B., Kim, R., Krauter, D., Kupari, J., Lemen, J., Lesnak, J.B., Liu, W., Lopez, I., Lu, Y., Macmillan, H.J., Mazhar, K., Meriau, P., Moffitt, J.R., Moreno, M.M., Mwirigi, J.M., Naz, H., O’Brein, J., Payne, M., Del Rosario, J., Rosen, S.F., Shiers, S., Simpson, E., Slivicki, R., Stone, J.R., Tavares-Ferreira, D., Uhelski, M., Woolf, C.J., Xu, Q., Yi, J., Yousuf, M.S., Zhu, D., Cavalli, V., Zhao, G., Olausson, H., Ernfors, P., Gereau, R.W., Luo, W., Price, T.J., and Renthal, W., A Reference Atlas of the Human Dorsal Root Ganglion. 2025, openRxiv.

54. Hartono, F., Tanjung, C., K, E.B., Marpaung, D., Ananditya, T., and Budisantoso, A.B. (2020) Catastrophic results due to unrecognizing of congenital insensitivity to pain with anhidrosis in children with multiple long bones fractures: A case report of 27 years follow-up of two siblings. Int J Surg Case Rep. 73: p. 213–217.

55. Bennett, D.L. and Woods, C.G. (2014) Painful and painless channelopathies. Lancet Neurol. 13(6): p. 587–99.

56. Chen, Y.-C., Auer-Grumbach, M., Matsukawa, S., Zitzelsberger, M., Themistocleous, A.C., Strom, T.M., Samara, C., Moore, A.W., Cho, L.T.-Y., Young, G.T., Weiss, C., Schabhüttl, M., Stucka, R., Schmid, A.B., Parman, Y., Graul-Neumann, L., Heinritz, W., Passarge, E., Watson, R.M., Hertz, J.M., Moog, U., Baumgartner, M., Valente, E.M., Pereira, D., Restrepo, C.M., Katona, I., Dusl, M., Stendel, C., Wieland, T., Stafford, F., Reimann, F., von Au, K., Finke, C., Willems, P.J., Nahorski, M.S., Shaikh, S.S., Carvalho, O.P., Nicholas, A.K., Karbani, G., McAleer, M.A., Cilio, M.R., McHugh, J.C., Murphy, S.M., Irvine, A.D., Jensen, U.B., Windhager, R., Weis, J., Bergmann, C., Rautenstrauss, B., Baets, J., De Jonghe, P., Reilly, M.M., Kropatsch, R., Kurth, I., Chrast, R., Michiue, T., Bennett, D.L.H., Woods, C.G., and Senderek, J. (2015) Transcriptional regulator PRDM12 is essential for human pain perception. Nature Genetics. 47(7): p. 803–808.

57. Lischka, A., Lassuthova, P., Çakar, A., Record, C.J., Van Lent, J., Baets, J., Dohrn, M.F., Senderek, J., Lampert, A., Bennett, D.L., Wood, J.N., Timmerman, V., Hornemann, T., Auer-Grumbach, M., Parman, Y., Hübner, C.A., Elbracht, M., Eggermann, K., Geoffrey Woods, C., Cox, J.J., Reilly, M.M., and Kurth, I. (2022) Genetic pain loss disorders. Nature Reviews Disease Primers. 8(1).

58. Boada, D.M., Martin, T.J., Peters, C.M., Hayashida, K., Harris, M.H., Houle, T.T., Boyden, E.S., Eisenach, J.C., and Ririe, D.G. (2014) Fast-conducting mechanoreceptors contribute to withdrawal behavior in normal and nerve injured rats. Pain. 155(12): p. 2646–2655.

59. Kuehn, E.D., Meltzer, S., Abraira, V.E., Ho, C.-Y., and Ginty, D.D. (2019) Tiling and somatotopic alignment of mammalian low-threshold mechanoreceptors. Proceedings of the National Academy of Sciences. 116(19): p. 9168–9177.

60. Nencini, S. and Ivanusic, J. (2017) Mechanically sensitive Aδ nociceptors that innervate bone marrow respond to changes in intra-osseous pressure. The Journal of Physiology. 595(13): p. 4399–4415.

61. Thai, J., Kyloh, M., Travis, L., Spencer, N.J., and Ivanusic, J.J. (2020) Identifying spinal afferent (sensory) nerve endings that innervate the marrow cavity and periosteum using anterograde tracing. Journal of Comparative Neurology. 528(11): p. 1903–1916.

62. Dye, S.F., Vaupel, G.L., and Dye, C.C. (1998) Conscious neurosensory mapping of the internal structures of the human knee without intraarticular anesthesia. Am J Sports Med. 26(6): p. 773–7.

63. Abraira, V.E., Kuehn, E.D., Chirila, A.M., Springel, M.W., Toliver, A.A., Zimmerman, A.L., Orefice, L.L., Boyle, K.A., Bai, L., Song, B.J., Bashista, K.A., O’Neill, T.G., Zhuo, J., Tsan, C., Hoynoski, J., Rutlin, M., Kus, L., Niederkofler, V., Watanabe, M., Dymecki, S.M., Nelson, S.B., Heintz, N., Hughes, D.I., and Ginty, D.D. (2017) The Cellular and Synaptic Architecture of the Mechanosensory Dorsal Horn. Cell. 168(1-2): p. 295–310.e19.

64. Traub, R.J. and Mendell, L.M. (1988) The spinal projection of individual identified A-delta- and C-fibers. J Neurophysiol. 59(1): p. 41–55.

65. Kokai, E., Alsulaiman, W.A., Dickie, A.C., Bell, A.M., Goffin, L., Watanabe, M., Gutierrez-Mecinas, M., and Todd, A.J. (2022) Characterisation of deep dorsal horn projection neurons in the spinal cord of the Phox2a::Cre mouse line. Mol Pain. 18: p. 17448069221119614.

66. Warwick, C., Salsovic, J., Hachisuka, J., Smith, K.M., Sheahan, T.D., Chen, H., Ibinson, J., Koerber, H.R., and Ross, S.E. (2022) Cell type-specific calcium imaging of central sensitization in mouse dorsal horn. Nature Communications. 13(1).

67. Yarmolinsky, D.A., Zeng, X., Mackinnon-Booth, N., Greene, C.A., Kim, C., Cheng, Y.-T., Lenfers Turnes, B., and Woolf, C.J. (2025) Differential modification of ascending spinal outputs in acute and chronic pain states. Neuron. 113(8): p. 1223–1239.e5.

68. Ahanonu, B., Crowther, A., Kania, A., Rosa-Casillas, M., and Basbaum, A.I. (2024) Long-term optical imaging of the spinal cord in awake behaving mice. Nature Methods. 21(12): p. 2363–2375.

69. Häring, M., Zeisel, A., Hochgerner, H., Rinwa, P., Jakobsson, J.E.T., Lönnerberg, P., La Manno, G., Sharma, N., Borgius, L., Kiehn, O., Lagerström, M.C., Linnarsson, S., and Ernfors, P. (2018) Neuronal atlas of the dorsal horn defines its architecture and links sensory input to transcriptional cell types. Nature Neuroscience. 21(6): p. 869–880.

70. Choi, S., Hachisuka, J., Brett, M.A., Magee, A.R., Omori, Y., Iqbal, N.-U.-A., Zhang, D., Delisle, M.M., Wolfson, R.L., Bai, L., Santiago, C., Gong, S., Goulding, M., Heintz, N., Koerber, H.R., Ross, S.E., and Ginty, D.D. (2020) Parallel ascending spinal pathways for affective touch and pain. Nature. 587(7833): p. 258–263.

71. Boyle, K.A., Gradwell, M.A., Yasaka, T., Dickie, A.C., Polgár, E., Ganley, R.P., Orr, D.P.H., Watanabe, M., Abraira, V.E., Kuehn, E.D., Zimmerman, A.L., Ginty, D.D., Callister, R.J., Graham, B.A., and Hughes, D.I. (2019) Defining a Spinal Microcircuit that Gates Myelinated Afferent Input: Implications for Tactile Allodynia. Cell Reports. 28(2): p. 526–540.e6.

72. Gautam, M., Yamada, A., Yamada, A.I., Wu, Q., Kridsada, K., Ling, J., Yu, H., Dong, P., Ma, M., Gu, J., and Luo, W. (2024) Distinct local and global functions of mouse Aβ low-threshold mechanoreceptors in mechanical nociception. Nature Communications. 15(1).

73. Krauter, D., Kupari, J., Usoskin, D., Su, J., Hu, Y., Zhang, M.-D., and Ernfors, P. (2025) Spatial organization, chromatin accessibility and gene-regulatory programs defining mouse sensory neurons. Communications Biology. 8(1).

74. Bhuiyan, S.A., Xu, M., Yang, L., Semizoglou, E., Bhatia, P., Pantaleo, K.I., Tochitsky, I., Jain, A., Erdogan, B., Blair, S., Cat, V., Mwirigi, J.M., Sankaranarayanan, I., Tavares-Ferreira, D., Green, U., McIlvried, L.A., Copits, B.A., Bertels, Z., Del Rosario, J.S., Widman, A.J., Slivicki, R.A., Yi, J., Sharif-Naeini, R., Woolf, C.J., Lennerz, J.K., Whited, J.L., Price, T.J., Robert, W.G.I., and Renthal, W. (2024) Harmonized cross-species cell atlases of trigeminal and dorsal root ganglia. Science Advances. 10(25).

75. Szczot, M., Bouchatta, O., Brodzki, M., Manouze, H., Carballo, G., Yu, H., Kindström, E., De-Faria, F., Thorell, O., Kao, A., Liljencrantz, J., Karlsson, C., Capitán, M.M., Ng, K., Frangos, E., Ragnemalm, B., Saade, D., Bharucha-Goebel, D., Szczot, I., Moore, W., Terejko, K., Cole, J., Bönnemann, C., Gerling, G., Mahns, D., Larsson, M., Luo, W., Marshall, A., Chesler, A., Olausson, H., and Nagi, S., PIEZO2-dependent rapid pain system in humans. 2024, Springer Science and Business Media LLC.

76. 76. Gatto, G., Bourane, S., Ren, X., Di Costanzo, S., Fenton, P.K., Halder, P., Seal, R.P., and Goulding, M.D. (2021) A Functional Topographic Map for Spinal Sensorimotor Reflexes. Neuron. 109(1): p. 91–104.e5.

77. Todd, A.J. (2010) Neuronal circuitry for pain processing in the dorsal horn. Nat Rev Neurosci. 11(12): p. 823–36.

78. Basbaum, A. (2022) History of Spinal Cord “Pain” Pathways Including the Pathways Not Taken. Frontiers in Pain Research. 3.

79. Browne, T.J., Smith, K.M., Gradwell, M.A., Iredale, J.A., Dayas, C.V., Callister, R.J., Hughes, D.I., and Graham, B.A. (2021) Spinoparabrachial projection neurons form distinct classes in the mouse dorsal horn. PAIN. 162(7).

80. Knuepfer, M.M. and Schramm, L.P. (1987) The conduction velocities and spinal projections of single renal afferent fibers in the rat. Brain Res. 435(1-2): p. 167–73.

81. De Groat, W.C. (2006) Integrative control of the lower urinary tract: preclinical perspective. British Journal of Pharmacology. 147(S2): p. S25–S40.

82. Strassman, A.M. and Levy, D. (2006) Response properties of dural nociceptors in relation to headache. J Neurophysiol. 95(3): p. 1298–306.

83. Cadden, S.W., Lisney, S.J., and Matthews, B. (1983) Thresholds to electrical stimulation of nerves in cat canine tooth-pulp with A beta-, A delta- and C-fibre conduction velocities. Brain Res. 261(1): p. 31–41.

84. Koutsioumpa, C., Santiago, C., Jacobs, K., Lehnert, B.P., Barrera, V., Hutchinson, J.N., Schmelyun, D., Lehoczky, J.A., Paul, D.L., and Ginty, D.D. (2023) Skin-type-dependent development of murine mechanosensory neurons. Developmental Cell. 58(20): p. 2032–2047.e6.

85. Madisen, L., Aleena, Shimaoka, D., Amy, Nathan, Li, L., Alexander, Niino, Y., Egolf, L., Monetti, C., Gu, H., Mills, M., Cheng, A., Tasic, B., Thuc, Susan, Benucci, A., Nagy, A., Miyawaki, A., Helmchen, F., Ruth, Knöpfel, T., Edward, R, Carandini, M., and Zeng, H. (2015) Transgenic Mice for Intersectional Targeting of Neural Sensors and Effectors with High Specificity and Performance. Neuron. 85(5): p. 942–958.

86. Madisen, L., Zwingman, T.A., Sunkin, S.M., Oh, S.W., Zariwala, H.A., Gu, H., Ng, L.L., Palmiter, R.D., Hawrylycz, M.J., Jones, A.R., Lein, E.S., and Zeng, H. (2010) A robust and high-throughput Cre reporting and characterization system for the whole mouse brain. Nature Neuroscience. 13(1): p. 133–140.

87. Daigle, T.L., Madisen, L., Hage, T.A., Valley, M.T., Knoblich, U., Larsen, R.S., Takeno, M.M., Huang, L., Gu, H., Larsen, R., Mills, M., Bosma-Moody, A., Siverts, L.A., Walker, M., Graybuck, L.T., Yao, Z., Fong, O., Nguyen, T.N., Garren, E., Lenz, G.H., Chavarha, M., Pendergraft, J., Harrington, J., Hirokawa, K.E., Harris, J.A., Nicovich, P.R., McGraw, M.J., Ollerenshaw, D.R., Smith, K.A., Baker, C.A., Ting, J.T., Sunkin, S.M., Lecoq, J., Lin, M.Z., Boyden, E.S., Murphy, G.J., Da Costa, N.M., Waters, J., Li, L., Tasic, B., and Zeng, H. (2018) A Suite of Transgenic Driver and Reporter Mouse Lines with Enhanced Brain-Cell-Type Targeting and Functionality. Cell. 174(2): p. 465–480.e22.

88. Lehnert, B.P., Santiago, C., Huey, E.L., Emanuel, A.J., Renauld, S., Africawala, N., Alkislar, I., Zheng, Y., Bai, L., Koutsioumpa, C., Hong, J.T., Magee, A.R., Harvey, C.D., and Ginty, D.D. (2021) Mechanoreceptor synapses in the brainstem shape the central representation of touch. Cell. 184(22): p. 5608–5621.e18.

89. McCoy, E.S., Taylor-Blake, B., and Zylka, M.J. (2012) CGRPα-Expressing Sensory Neurons Respond to Stimuli that Evoke Sensations of Pain and Itch. PLoS ONE. 7(5): p. e36355.

90. Huey, E.L., Turecek, J., Delisle, M.M., Mazor, O., Romero, G.E., Dua, M., Sarafis, Z.K., Hobble, A., Booth, K.T., Goodrich, L.V., Corey, D.P., and Ginty, D.D. (2025) The auditory midbrain mediates tactile vibration sensing. Cell. 188(1): p. 104–120 e18.

91. Qi, L., Iskols, M., Greenberg, R.S., Xiao, J.Y., Handler, A., Liberles, S.D., and Ginty, D.D. (2024) Krause corpuscles are genital vibrotactile sensors for sexual behaviours. Nature. 630(8018): p. 926–934.

92. Thai, J., Fuller-Jackson, J.P., and Ivanusic, J.J. (2024) Using tissue clearing and light sheet fluorescence microscopy for the three-dimensional analysis of sensory and sympathetic nerve endings that innervate bone and dental tissue of mice. Journal of Comparative Neurology. 532(1).

93. Ko, F.C., Fullam, S., Lee, H., Chan, K., Ishihara, S., Adamczyk, N.S., Obeidat, A.M., Soorya, S., Miller, R.J., Malfait, A.M., and Miller, R.E. (2025) Clearing-enabled light sheet microscopy as a novel method for three-dimensional mapping of the sensory innervation of the mouse knee. J Orthop Res. 43(3): p. 632–639.

94. Lauria, G., Hsieh, S.T., Johansson, O., Kennedy, W.R., Leger, J.M., Mellgren, S.I., Nolano, M., Merkies, I.S.J., Polydefkis, M., Smith, A.G., Sommer, C., and Valls-Solé, J. (2010) European Federation of Neurological Societies/Peripheral Nerve Society Guideline on the use of skin biopsy in the diagnosis of small fiber neuropathy. Report of a joint task force of the European Fe-deration of Neurological Societies and the Peripheral Ne. European Journal of Neurology. 17(7): p. 903.

95. Emanuel, A.J., Lehnert, B.P., Panzeri, S., Harvey, C.D., and Ginty, D.D. (2021) Cortical responses to touch reflect subcortical integration of LTMR signals. Nature. 600(7890): p. 680–685.

96. Turecek, J. and Ginty, D.D. (2024) Coding of self and environment by Pacinian neurons in freely moving animals. Neuron. 112(19): p. 3267–3277 e6.

