## Supplementary material for "Fast-conducting mechanonociceptors uniquely engage reflexive and affective pain circuitry to drive protective responses": Table 1

| Glabrous-innervating primary sensory neuron subtypes | | | | | |
| --- | --- | --- | --- | --- | --- |
| Driver line | Physiology  (mechanical and thermal) | Other common names | Withdrawal response | Nocifensive response* | Place aversion |
| *Smr2^Cre^* | Aδ-high threshold mechanoreceptors | Myelinated nociceptors/ peptidergic nociceptors | ✔ | ✔ | ✔ |
| *Bmpr1b^Cre^* | Aδ-high threshold mechanoreceptors | Myelinated nociceptors/ peptidergic nociceptors | ✔ | ✔ | ✔ |
| *Sstr2^CreER^* | C heat thermoreceptors | peptidergic nociceptors | ✔ | ✔ | ✔ |
| *MrgprD^CreER^* | C heat and high-threshold mechanoreceptors | nonpeptidergic nociceptors | ✔ | ✗ | ✗ |
| *MrgprA3^Cre^* | C heat and high-threshold mechanoreceptors | nonpeptidergic nociceptors/ pruriceptors | ✗ | ✗ | ✗ |
| *MrgprB4^Cre^* | C heat and high-threshold mechanoreceptors | nonpeptidergic nociceptors/ pruriceptors | ✗ | ✗ | ✗ |
| *Cysltr2^Cre^* | C heat and high-threshold mechanoreceptors | nonpeptidergic nociceptors/ pruriceptors/ *Sst^+^* | ✔ | ✗ | ✗ |
| *Trpm8^FlpO^* | C cold thermoreceptors |  | ✔ | ✗ | ✗ |
| *Ptgfr^CreER^* | Aβ high-threshold mechanoreceptors |  | ✔ | ✗ | ✗ |
| *TrkB^CreER^* | Aβ low-threshold mechanoreceptors |  | ✗ | ✗ | ✗ |

**Table 1. Glabrous skin-innervating primary sensory neuron types.** A list of transcriptionally and physiologically defined DRG neuron subtypes and genetic driver lines used to manipulate them for the gain-of-function optogenetic screen described in Figure 1. Results of the screen are summarized. We define a “nocifensive” response as one that includes robust paw shaking or jumping.
